# Parallel processing chains span cytoarchitectures to organize association cortex

**DOI:** 10.64898/2026.04.21.717753

**Authors:** Evan M. Gordon, Aishwarya Rajesh, Roselyne J. Chauvin, Alyssa Labonte, Babatunde Adeyemo, Ally Dworetsky, Charles J. Lynch, Samuel R. Krimmel, Philip Cho, Anxu Wang, Noah J. Baden, Kristen M. Scheidter, Julia Monk, Athanasia Metoki, Jianxun Ren, Tomoyuki Nishino, Youngeun Park, Emily Rafka, John R. Pruett, Adam Kepecs, Hesheng Liu, Damien A. Fair, Conor Liston, Choong-Wan Woo, Benjamin P. Kay, Scott Marek, Steven E. Petersen, Chad M. Sylvester, Rebecca F. Schwarzlose, Marcus E. Raichle, Timothy O. Laumann, Nico U.F. Dosenbach

## Abstract

Task fMRI^1^ and electrophysiology^2^ have revealed distributed, linked cortical patches with shared category preferences (e.g., faces, objects, places)^1,3–5^, smaller than cytoarchitectonic areas. Resting-state functional connectivity (RSFC) similarly showed that somato-cognitive action network (SCAN) nodes interleave with effectors (foot, hand, mouth), subdividing the precentral gyrus^6^. Here, using multiple precision functional mapping (PFM) modalities (RSFC, task, lags), we discovered that most of association cortex is organized like face processing and SCAN, with small, discrete patches interconnected into chains. Such patch-chains densely tile prefrontal cortex but are largely absent from primary cortex. Cortico-striatal connectivity is organized such that patches of the same chain connect to the same striatal location. Within chains, infra-slow fMRI signals are ordered in time. RSFC-defined chains align with task fMRI localizers (e.g., visual, motor, pain). Chains are absent at birth and emerge in the first year of life, suggesting their formation is at least partially experience-driven. Cytoarchitectonic areas are subdivided by patches, and patches in the same chain are distributed across different cytoarchitectures. Chains represent parallel ordered processing streams that are separated by information domain and behavioral goals, not cytoarchitectonics. Functional subdivision of architectonics into smaller patches, interlinked to form cross-architecture chains, enable greater parallelization and flexible specialization of processing.

## MAIN

Historically, cortical organization was viewed as a set of functions each localized to a particular cortical territory^7^, such as speech production in the left inferior frontal gyrus^8^.

Later lesion studies suggested that higher-order functions involve multiple distributed regions^9–11^. This perspective was expanded by human task-based functional magnetic resonance imaging (fMRI)^12^ and resting-state functional connectivity (RSFC), which observed that complex tasks evoked activity in multiple functionally networked regions, distributed across cortical lobes^13–18^. This led to the influential view that the brain is organized into large-scale functional networks^10^, with cognitive functions requiring multiple processing operations that are implemented via the interaction of centimeter-scale, functionally specialized regions^19,20^.

The basic functional units of this organization were most commonly argued to be architectonic cortical areas^21^. First described by Brodmann in 1909^22^, cytoarchitectonic areas cover the cerebral cortex as discrete centimeter-scale regions^23^ that dissociate from each other early in postnatal development via the input-dependent discretization of initially continuous cortical gradients^24–27^. Each cortical area has a distinct intrinsic cellular composition, laminar organization, and circuitry^28,29^, that is thought to be optimized for a specific type of computation. Cortical areas connect to subcortical structures including the striatum, thalamus, cerebellum, and hippocampus^30–32^, which have relatively more homogenous cytoarchitecture^33^. Projections from many cortical areas often converge on a single location within a subcortical structure^34,35^.

Human task fMRI and RSFC studies investigating cognitive functions in association cortex traditionally averaged data across individuals^36^, and so identified networked functional responses at a spatial scale (centimeters) similar to that of architectonic divisions^12–14,37^. Therefore, most prior work, including our own, presumed that functionally distinct cortical regions described by neuroimaging would correspond to single or multiple adjacent centimeter-scale cytoarchitectonic cortical areas^28,29,38–43^. However, functional and architectonic parcellations of cortex only ever achieved reasonable convergence in primary sensory and motor cortex, not in association regions, with especially poor alignment in frontal cortex^40,44^ (see Supplementary Fig. 1). We and others presumed these structure-function inconsistencies to be a limitation of the available neuroimaging data, which would resolve with further advances in resolution and signal quality^28,40,41^.

The functional organization of visual processing systems, which is most effectively examined using individual-specific localizer methods rather than group-averaging^1,45^, never supported the existence of centimeter-scale functional regions or a strict function-to-architectonic convergence^1,3,4,46–54^. The temporal cortex of humans and macaques is tiled by small patches, only a few millimeters across, that selectively respond to specific categories of visual stimuli such as faces or objects^1,3,4,46–53,55^. Patches are networked to process stimuli in a partially ordered processing stream progressing from featural and view-dependent, to holistic and view-invariant, to stimulus identity and familiarity^3,56–60^.

Recently, visual category preferences have been reported beyond classical visual regions in temporal, prefrontal, and parietal cortex^61,62^. Most critically, patches divide rather than converge with architectonic divisions^3,5,54^. Important rodent work has described similar mismatches between architectonics and electrophysiological measures^63^. Together, these observations point to intrinsic connectivity rather than architectonics as a primary organizing principle of cortical function^63,64^.

In humans, intrinsic connectivity can be comprehensively characterized using precision functional mapping (PFM), which uses RSFC to detail functional connections across the whole brain within single individuals, thus avoiding pitfalls associated with group averaging^44,65–71^. Contrary to our expectations, PFM did not resolve the mismatch between function and architectonics but instead deepened it. Classical cytoarchitectonic Brodmann areas span the full length of the central sulcus (Brodmann area [BA] 4,3,2,1), with borders separating anterior motor (BA 4, precentral gyrus) from posterior somatosensory (BA 3,2,1) areas^22,72^. By contrast, localizer task fMRI and RSFC revealed an orthogonal functional organization that subdivides the putatively architectonically uniform precentral gyrus^6,13,14^. The somato-cognitive action network (SCAN), a chain of functionally connected action regions within BA 4, alternates with effector-specific regions for movement of the foot, hand, and mouth^6^. Other PFM studies revealed interdigitated functional regions in frontal^62,73–76^ and parietal cortex^70,71,77,78^ much smaller than group-averaged centimeter-scale functional responses or architectonic areas.

Motivated by the deep insights into brain function provided by research on category-selective visual processing patches (e.g., faces, places)^1,5,45,46,61^ and by the discovery that the SCAN chain of action regions interrupts effector-specific motor cortex^6^, we set out to characterize the functional organization of human cerebral cortex in greatest possible detail. Using functional connectivity, we systematically identified chains as multiple non-adjacent functionally connected patches. We developed a measure of catenation (from the Latin *catena*, meaning chain) to comprehensively map the density of these chains across cortex. Chain maps were then compared to task fMRI localizers and architectonic divisions, and their cortico-subcortical connectivity was detailed. These analyses utilized an extensive collection of PFM datasets (*n* = 48, totaling 160.8 total hours of functional connectivity data); these included ultra-short TR and 7 Tesla data, and ranged in ages from newborn to adult, enabling us to examine questions of signal timing and development. Results were verified in individual-specific data from the Human Connectome Project (HCP, n∼ 1,000)^79,80^ and the Adolescent Brain Cognitive Development Study (ABCD, n∼ 4,000)^81^.

### Chains of functional connectivity patches

Individual-specific functional connectivity maps revealed chains of cortical patches similar to SCAN (Fig. 1a) at many seed locations (see examples highlighted in Fig. 1b-d, Extended Data Fig. 1 and Supplementary Figs. 2-6). Replicating our prior work^82^, a seed placed in the center of the precentral gyrus (MNI:-35, 15, 45) identified the superior middle and inferior SCAN nodes (Fig. 1a), which alternate with effector-specific motor regions for control of the foot, hand, and mouth^82^. Seeds placed in dorsolateral (Fig. 1b; MNI:-38, 7, 36) and dorsomedial (Fig. 1c; MNI: 7, 26, 48) frontal cortex also showed chained connectivity patterns. A seed in superior parietal cortex (Fig. 1d; MNI:-30,-55, 50) revealed a chain of functionally connected nodes running from occipital cortex to posterior middle frontal gyrus, consistent with the dorsal visual pathway^83^. Extended Data Fig. 1 and Supplementary Fig. 2 show additional examples of functional connectivity chains in the same exemplar participant and demonstrate how each chain fits within a known functional network. Similar connectivity chains were identified in all individuals (*n* = 27 PFM data sets, Supplementary Figs. 3-6). These functional connectivity chains were reliable within-individual, based on split-quarter replication (Supplementary Fig. 7). Seed maps revealed strongly dissociated and, in some cases, anticorrelated connectivity in closely adjacent cortical locations (Fig. 1e, Supplementary Fig. 8). While connectivity chains were apparent in every individual (Supplementary Figs. 3-6), in most cases they were impossible to identify in data that had first been group-averaged (Fig. 1e-f, Extended Data Fig. 2), with SCAN being a notable exception^6^. Group-averaged data exhibited similar overall distributions of connectivity but lacked the spatial specificity to visualize discrete chain patches.

**Fig. 1.**
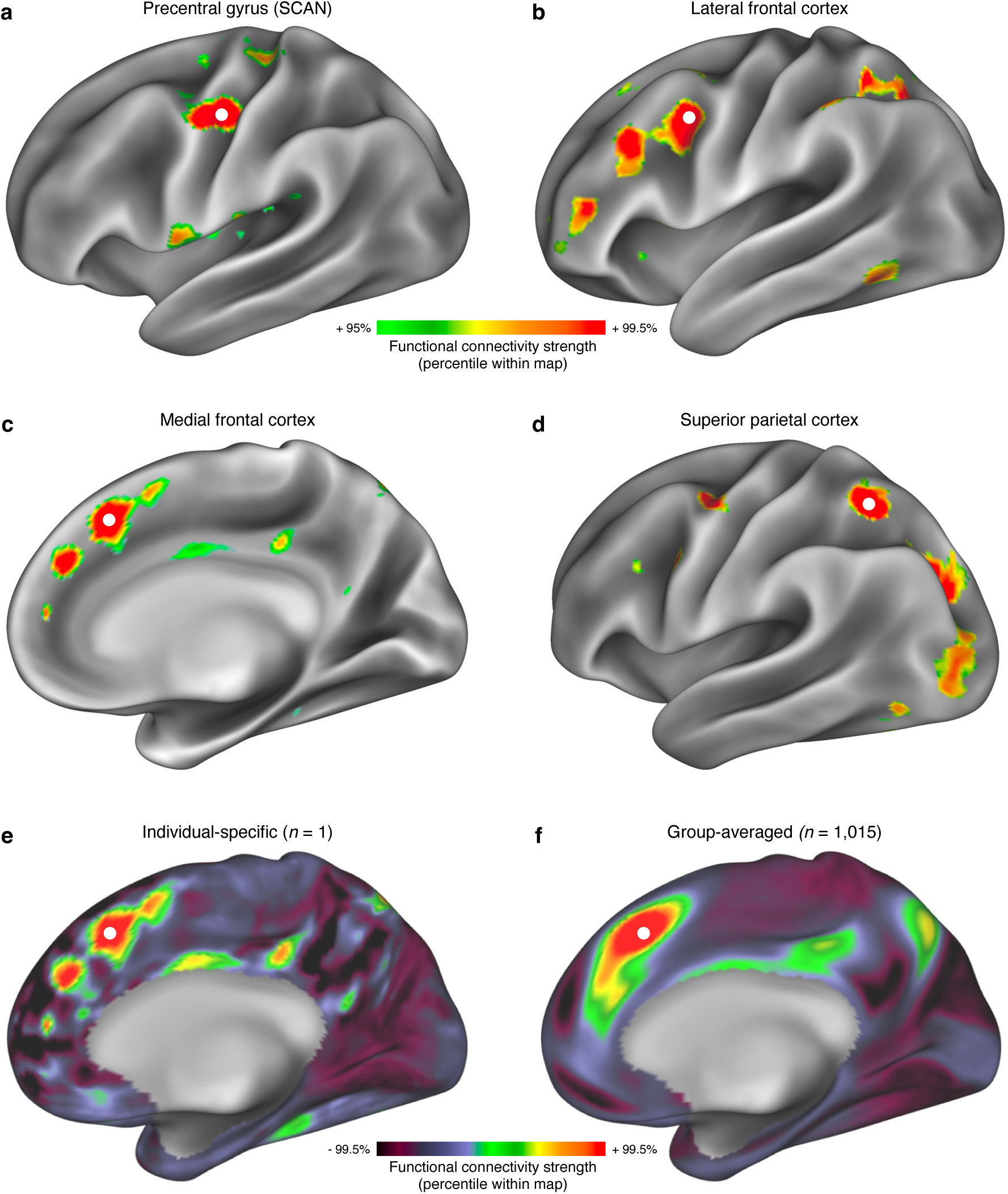
Chains of functional connectivity. a-d, Functional connectivity seed maps (thresholded to retain the top 5% of vertices), in a single exemplar individual (P1, 21-year-old female, 477 min resting-state fMRI). **a,** Somato-cognitive action network (SCAN) in precentral gyrus^6^ (seed MNI coordinates [x, y, z] = [-35, 15, 45]); **b,** lateral frontal cortex ([-38, 7, 36]); **c**, medial frontal cortex ([7, 26, 48]), and **d**, superior parietal cortex ([-30,-55, 50], approximate area LIPv^41^). These maps revealed chains of functionally connected patches. See Extended Data Fig. 1, Supplementary Fig. 2 for additional examples of functional connectivity chains in exemplar participant P1. See Supplementary Figs. 3-6 for matched chains in all *n* = 27 individual study participants. **e-f,** Unthresholded functional connectivity seed maps. See Supplementary Fig. 8 for all unthresholded connectivity seed maps. **e,** Chains were easily observed in individual-specific data from highly-sampled participants (here P1; seed: [7, 26, 48]); **f,** but not in group-averaged data from the Human Connectome Project dataset (HCP, *n* = 1,015; same seed). See Extended Data Fig. 2 for more examples of chains obscured by group-averaging of functional connectivity data.

### Chained connectivity in association cortex

To characterize the spatial distribution of functional connectivity chains, we quantified the chained nature of connectivity seeded from each cortical vertex (Fig. 2, Extended Data Fig. 1) using a catenation score (from *catena*, Latin for chain). In every individual, all connectivity maps seeded from every point in cortex were compared to a set of chain-like patterns (see Methods, Supplementary Fig. 9). This procedure successfully identifies chained patches (Extended Data Fig. 1).

**Fig. 2.**
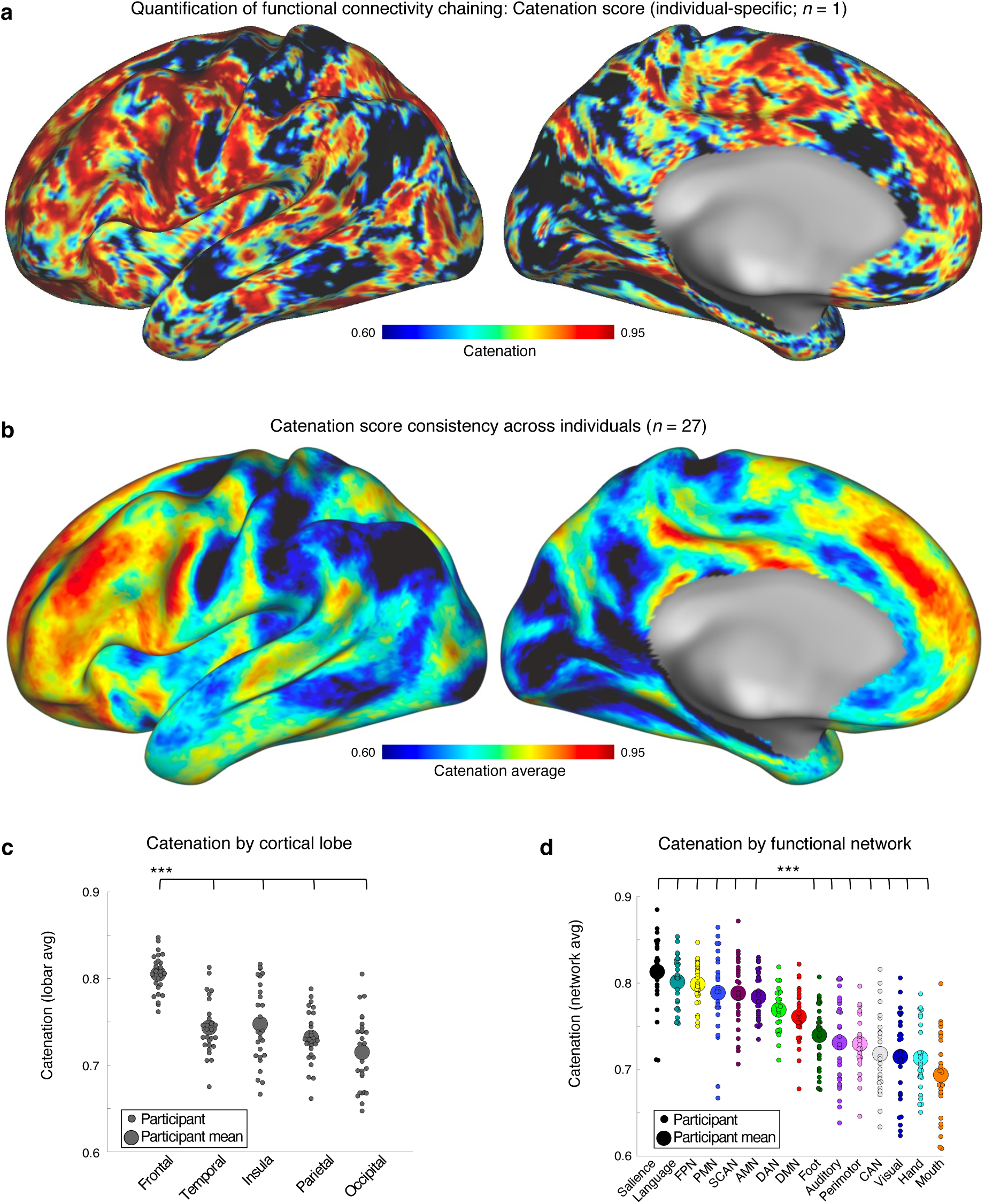
Quantification of functional connectivity chaining patterns via catenation scores. The presence of chained functional connectivity was quantified using the catenation score, which assesses the strength of fit of connectivity patterns from every cortical vertex against a set of template chain-like connectivity patterns (see Methods, Supplementary Fig. 9). Catenation score maps are shown for **a,** the individual exemplar participant (P1) and **b**, averaged across *n* = 27 repeatedly sampled individuals. See Supplementary Fig. 11 for vertexwise statistical testing of catenation map in **b** across participants. See Extended Data Fig. 5 for similar results using an alternative measure. **c,** Frontal cortex exhibited greater catenation than temporal, insular, parietal and occipital cortex. **d**, Most association networks, including salience, language, fronto-parietal (FPN), parietal memory (PMN), somato-cognitive action (SCAN), and action-mode (AMN) networks, exhibited significantly greater catenation than primary networks including auditory, visual, and motor foot, hand, and mouth networks, as well as the premotor and contextual association (CAN) networks. ***: *P* < 0.001 in two-tailed paired *t*-tests, Bonferroni-corrected for multiple comparisons.

Catenation was strongest in association cortex, especially in the frontal lobes (Fig. 2a, exemplar participant; Fig. 2b, average catenation maps; Supplementary Fig. 10, catenation in each participant; Supplementary Fig. 11, statistical testing against a null model). Frontal cortex was more catenated than temporal, insula, parietal, and occipital cortex in every individual (Fig. 2c; all paired *t*-tests: *t*s(26) > 8.15, *P*s < 10^-6^ after Bonferroni correction for multiple comparisons). Most association networks, including salience, language, fronto-parietal, parietal memory, somato-cognitive action (SCAN), and action-mode (AMN), were more catenated than all primary networks (auditory, visual, and somatomotor foot, hand, and mouth), perimotor, and the contextual association network (CAN) (Fig. 2d; all paired *t*-tests: *t*s(26) > 4.52, *P*s < 0.001 after Bonferroni correction for multiple comparisons). Group-averaging fMRI data obscured catenation (Fig 1f; Extended Data Fig. 4), but we were able to calculate catenation scores on individual-specific Human Connectome Project (HCP; *n* = 1,015) and Adolescent Brain Cognitive Development Study (ABCD; *n* = 3,928) data and then average the respective catenation maps. These analyses showed the pattern of catenation to be nearly identical for the PFM, HCP (*r_spatial_* = 0.83, *P_spin_* < 0.001), and ABCD (*r_spatial_* = 0.78, *P_spin_* < 0.001) datasets (Extended Data Fig. 5).

Maps of Moran’s I^84^, an alternative metric that quantifies the spatial frequency of connectivity maps rather than explicitly identifying chains, produced an almost identical pattern (catenation vs Moran’s I spatial correlations: exemplar participant P1: *r*_spatial_ = - 0.35, *P*_spin_ < 0.001; participant average: *r*_spatial_ =-0.32, *P*_spin_ = 0.004; Extended Data Fig. 5), with reduced Moran’s I values indicating more distributed connectivity in association relative to primary cortex.

Thus, distributed functional connectivity chains are common throughout much of association, but not primary cortex, with greatest concentration in the frontal lobes. Association cortex without chains was limited to the parietal nodes of the default-mode (DMN) and context association (CAN) networks, which are functionally connected to the anterior and middle hippocampus^85^.

### Connectivity chains tile prefrontal cortex

High catenation scores throughout prefrontal cortex imply that many functional connectivity chains lie near each other. To investigate this possibility, we placed connectivity seeds across all of medial (examples in Fig. 3a) and lateral (examples in Extended Data Fig. 6a) prefrontal cortex. We found that most of prefrontal cortex is segregated into functional connectivity chains that closely abut each other. In the exemplar participant (P1), at least six distinct chains tile the medial superior frontal cortex (Fig. 3a for seed-based connectivity patterns; Fig. 3b for algorithmically identified connectivity chains (see Methods); Extended Data Fig. 6a,b for lateral prefrontal chains). Across PFM participants, closely juxtaposed connectivity chains tile much of the prefrontal cortex (mean ± SD = 63% ± 9% of frontal cortex across participants). These connectivity chains fall within multiple canonical large-scale functional networks including language, fronto-parietal, action-mode, salience, and default-mode (see Fig. 3b; Extended Data Fig. 6b; Extended Data Fig. 1; Supplementary Fig. 2). Thus, a given region of cortex will contain multiple distinct but interlocking chains belonging to different larger canonical networks. Multiple distinct chains belonging to the same larger functional network can also be represented in the same region of cortex (Fig. 3b, FPN chains; see also Extended Data Fig. 1).

**Fig. 3.**
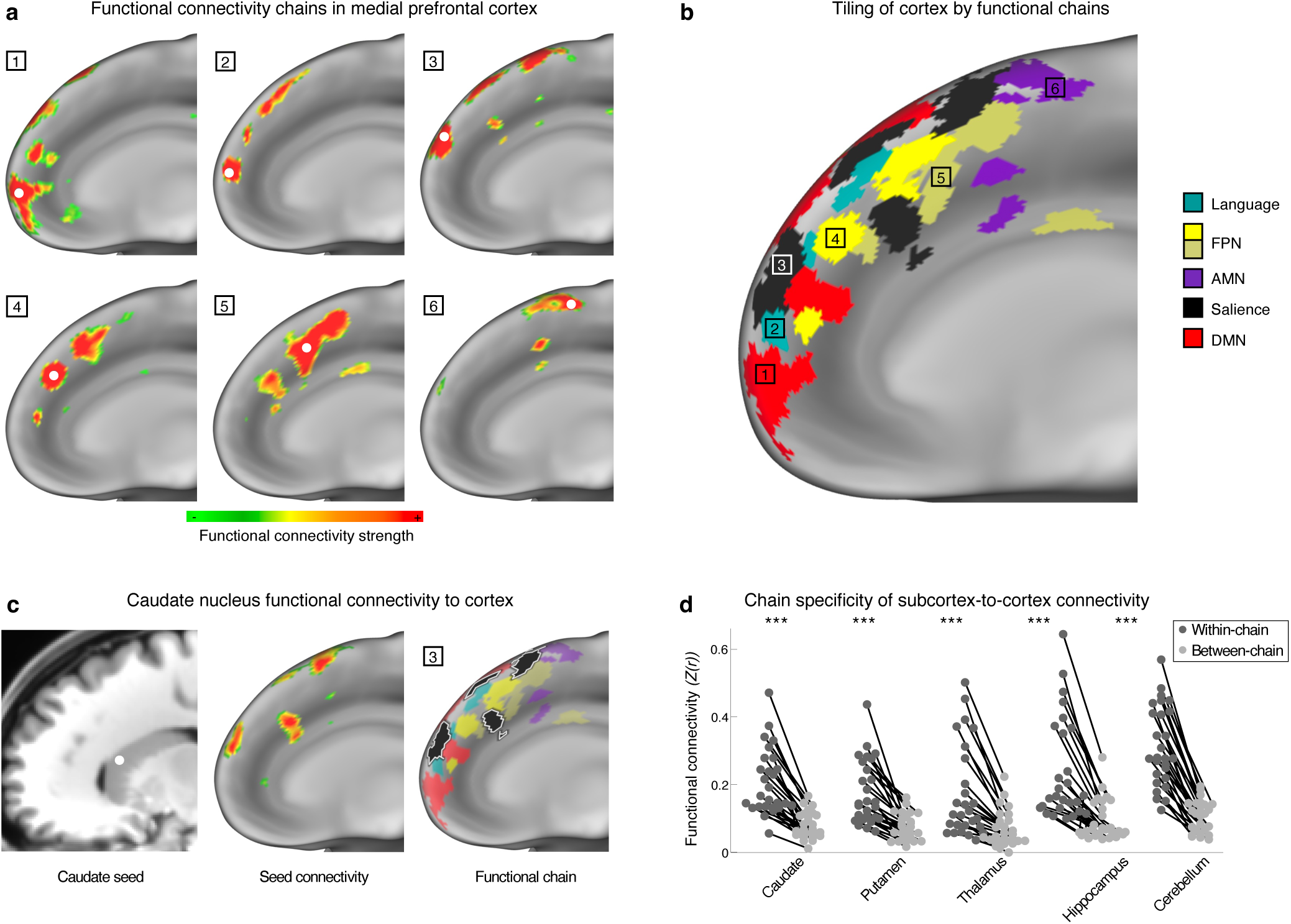
Density and subcortical connectivity of functional chains tiling prefrontal cortex. **a**, Connectivity seeds (1-6) placed in medial prefrontal cortex of the exemplar participant (P1) revealed a series of discrete, interleaved functional connectivity chains. **b**, A community detection algorithm (see Methods) defined the same chains (1-6) in a data-driven manner, revealing that connectivity chains closely abut each other and together tile most of medial frontal cortex. Chains are color-coded according to the large-scale canonical functional networks they are part of (language = teal, fronto-parietal network (FPN) = yellows, action-mode network (AMN) = purple, salience = black, default-mode network (DMN) = red. See Extended Data Fig. 6 for examples in lateral prefrontal cortex in P1. **c,** A seed located in left dorsal caudate [MNI: 15, 14, 12] (P1) connects to the same chain (#3) identified via cortical seed in **a** and community detection algorithm in **b**. **d,** For all subcortical locations, functional connectivity to patches of the most strongly connected cortical chain (dark grey dots) was stronger than connectivity to other patches interspersed with that chain (light gray dots). Each dot represents the average within-or between-chain connectivity across voxels within one subcortical structure in a single participant (*n* = 27); lines connect measures from the same individual. ***: *P* < 0.001 in two-tailed paired *t*-tests, Bonferroni-corrected for multiple comparisons.

In each participant, a mean of 136.7 (SD: ± 11.3) distinct chains were identified; each chain contained 3.4 patches (SD: ± 0.6). Across participants, chain patches averaged 175.9 (SD: ± 144.1) mm^2^ in cortical surface area and were spaced at an average inter-patch Euclidean distance of 22.5 ± 8.4mm, and a geodesic distance of 39.0 ± 12.9 mm along the unfolded cortical sheet. Patches were larger in anterior parietal and superior temporal lobes (see Supplementary Fig. 12 for maps of patch size and inter-patch distance). These distances are long to be spanned by unmyelinated intracortical axons but short to be linked by major axonal fiber bundles in deep white matter (e.g., superior longitudinal fasciculus). Hence, we investigated whether functional connectivity chains may be preferentially supported by the superficial white matter system (also known as u-fibers), a diffuse collection of short-range fibers running underneath association cortex^86–88^. To test this idea, we identified white matter regions within the superficial u-fiber white matter system using the O’Donnell Research Group (ORG) atlas^89^ (Extended Data Fig. 7a) and computed the catenation of cortex directly above superficial white matter (Extended Data Fig. 7b). Average catenation of cortex overlaying superficial white matter was significantly higher than for the null model (spin test: *P* = 0.026, Extended Data Fig. 7c), suggesting that catenated cortex may be preferentially supported by the superficial white matter system.

In addition to cortico-cortical connections, cortex is linked to subcortical structures including striatum, hippocampus, and cerebellum. These connections often appear to be many-to-one, such that projections from multiple cortical areas converge onto a single subcortical target^34,35^. We hypothesized that if cortico-cortical connections are organized into chains, then subcortico-cortical connectivity should also reflect cortical catenation, such that a given subcortical location should be preferentially functionally connected to patches within the same chain, rather than to interleaved cortical patches belonging to other chains. As expected, functional connectivity seeds placed in subcortical structures exhibited strong and specific connectivity to multiple patches within the same cortical chain (e.g., dorsal caudate connected to chained patches in cingulate and superior frontal gyrus; Fig. 3c). Voxels within a subcortical structure exhibited stronger connectivity to a specific chain than to other patches in the same region. This chain specificity was true for every individual in caudate, putamen, thalamus, hippocampus, and cerebellum; all statistical comparisons were significant (Fig. 3d; two-tailed paired t-tests: all *t*s(26) > 6.10; all *P*s (Bonferroni-corrected) < 10^-5^). Thus, functional connectivity between cortex and subcortex is organized by chain.

Highly catenated cortex exhibited stronger connectivity to striatum than low-catenation cortex (paired *t*-tests: PFM – *t*(26) = 3.70, *P* = 0.001; HCP – *t*(1014) = 42.4, *P* < 0.001; ABCD – *t*(3927) = 91.8, *P* < 0.001), while low-catenation regions exhibited stronger connectivity to hippocampus than highly catenated regions (paired *t*-tests: PFM – *t*(26) = 2.37, *P* = 0.025; HCP – *t*(1014) = 52.5, *P* < 0.001; ABCD – *t*(3927) = 58.1, *P* < 0.001) (Extended Data Fig. 8). Hippocampal connectivity to cortical regions with chains (such as medial PFC) is thus organized by chain (Fig. 3d), but the bulk of hippocampal connections are to uncatenated cortex (e.g. angular gyrus and medial parietal cortex; Fig. 2b).

Seeding connectivity from specific cortical chains revealed strong connectivity to single locations in the striatum. By contrast, cortical chains were functionally connected to multiple regions in the cerebellum arranged in a chain-like pattern (Extended Data Fig. 6c). Across participants, functional connectivity seeded from cortical chains rarely exhibited multiple discrete connectivity foci (i.e. a chain-like pattern) in caudate, putamen, thalamus, and hippocampus (chain-like patterns observed in <25% of each structure) (Extended Data Fig. 6d). This suggests that cortico-subcortical connectivity is broadly many-nodes-to-one for striatum and thalamus. By contrast, on average more than 60% of the cerebellum exhibited multiple discrete connectivity foci, more than any other tested subcortical structure (*t*s(26) > 14.2, *P*s (corrected) < 10^-12^). This suggests a many-to-many organization for cortico-cerebellar connections.

### Chains are functional processing streams

While connectivity chains are most prevalent in prefrontal cortex, its complex organization and function are not yet well characterized. By contrast, the functions of visual processing regions are better understood. Imaging, electrophysiological, and anatomical research in humans and macaques has revealed that inferior temporal cortex contains multiple spatially segregated patches that selectively process faces, objects, and scenes^1,4,5,46,47,51,53,57^. Patches with the same category-selectivity (e.g., faces) are anatomically connected to each other^91^. These functionally segregated, category-selective patch systems tile large sections of temporal cortex^3^. Thus, we tested whether functional connectivity chains in temporal cortex correspond to these category-selective patch systems (faces, objects, places).

Functional connectivity and task localizer fMRI data collected at 7 Tesla were examined in eight highly sampled participants from the Natural Scenes Database (NSD)^66,90,92^. In each participant, a visual category functional localizer task^93^ identified a distributed system of face-selective patches in posterior inferior and lateral temporal cortex (Fig. 4a; Supplementary Fig. 13) consistent with past human and non-human primate research^1,4,51^. Functional connectivity seeded from the anterior inferior temporal face patch (likely fusiform face area (FFA)^1^; see Methods) delineated a connectivity chain (Fig. 4b, Supplementary Fig. 13) that converged with the face patches (Fig. 4c, Supplementary Fig. 13). Functional connectivity from FFA to other face patches was stronger than a rotated null model in 7 of 8 participants (*P*s < 0.05 – 0.001; Fig. 4d). Similar congruence between connectivity chains and task fMRI localizer maps were observed in all cases. Specifically chains in temporal and occipital cortex overlapped with activations for other visual stimulus categories (objects, Extended Data Fig. 9a; places, Extended Data Fig. 9b), and frontal cortex AMN chains overlapped with pain activations (Extended Data Fig. 9c) (*P*s < 0.05 – 0.001 for each activation pattern in each individual participant).

**Figure 4.**
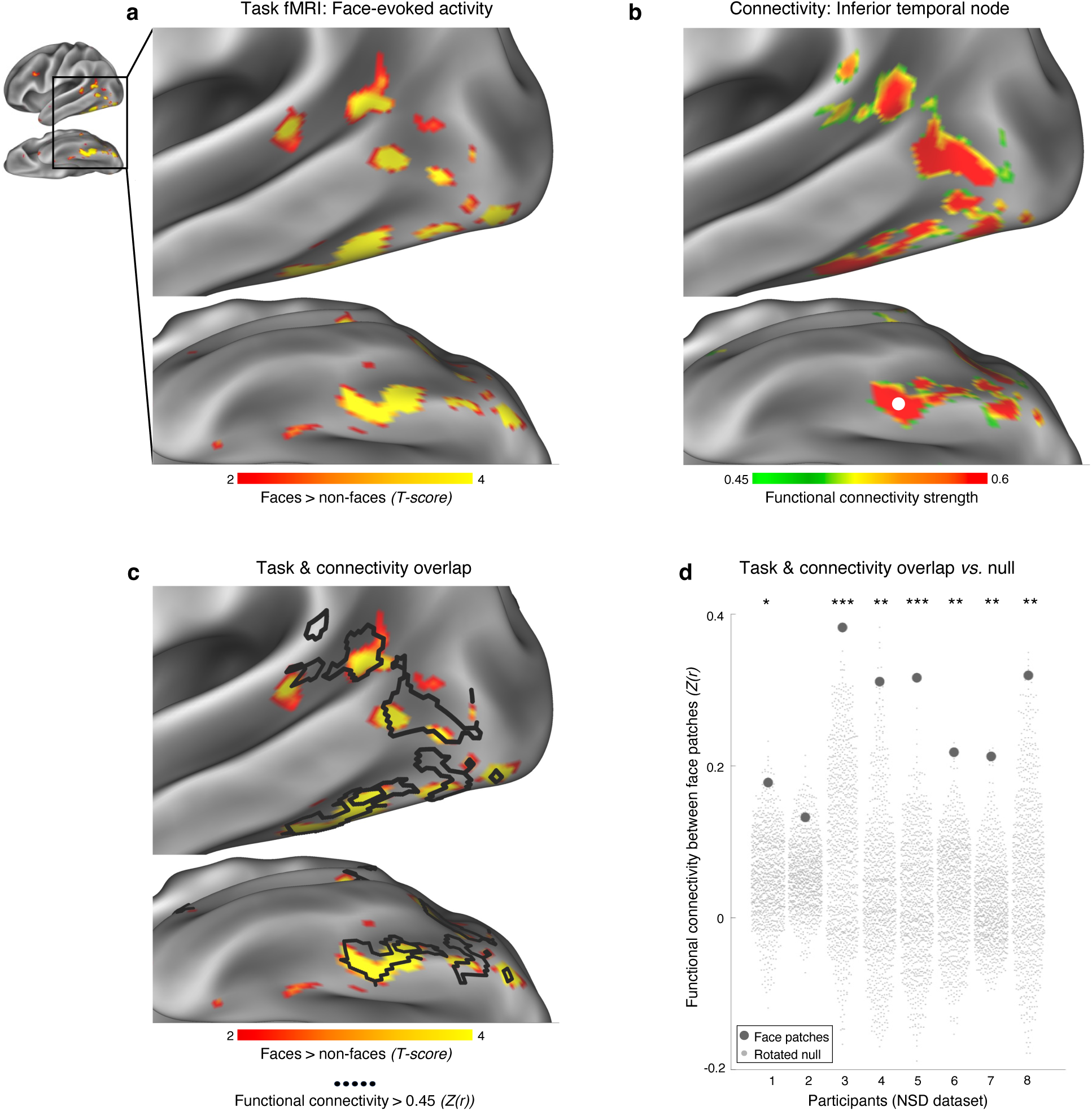
Task fMRI activations map onto functional connectivity chains. a, Task contrast localizing activity related to viewing faces (face stimuli > all other stimulus types, thresholded T > 2.0) demonstrates a patchy pattern of activity in posterior lateral and inferior temporal cortex (participant NSD05 from the Natural Scenes Dataset [NSD^90^]), consistent with prior work^1,4,51^. **b,** A functional connectivity seed placed in the anterior face-active region in inferior temporal cortex (from **a**, location convergent with putative fusiform face area [FFA^1^], thresholded Z(r) > 0.45) demonstrated a chained connectivity pattern convergent with that of face-evoked activity. **c,** Overlap of face-evoked activity (from **a,** red-yellow) with functional connectivity (black outline). See Supplementary Fig. 13 for other participants and Extended Data Fig. 9 for other task contrasts. **d,** Task fMRI and functional connectivity converged, such that connectivity seeded from the anterior face-active region of inferior temporal cortex (putative FFA) to other face-active regions (dark gray dots) was stronger than would be expected by chance (null rotated face patches: light gray dots), in seven of eight participants (Natural Scenes Database (NSD) dataset). *** indicates *P* < 0.001; ** indicates *P* < 0.01; * indicates *P* < 0.05.

Category-selective visual processing is known to be directed, with regions processing simpler visual features generally feeding forward into proximal regions that compute stimulus identity (with some reversed projections for feedback)^57^. It is possible that chains across the brain similarly represent directed functional processing streams. If so, their organization should also be ordered, with stronger connectivity between proximal patches (see example in Extended Data Fig. 10a), rather than an undirected net of mutually interconnected patches. Indeed, connectivity between patches of a chain was stronger when they were closer together. This effect was significant across all participants (connectivity vs distance: *r* =-0.34, *P* < 0.001; Extended Data Fig. 10b) and within each participant (*r*s =-0.22 –-0.49; *P*s < 10^-6^, Bonferroni corrected for multiple comparisons; Extended Data Fig. 10c). This within-chain effect was significantly stronger than the connectivity-distance relationship for cross-chain patches (paired t-test: *t*(26) = 7.42, *P* < 0.001). The preferential connectivity between adjacent links of a chain supports the notion that they not only look like chains but also function as directed processing streams.

Ordered processing is thought to be represented by a lagged temporal structure of brain activity, known to propagate in waves along sections of cortex^6,94–101^. In medial prefrontal cortex, chains run anterior-to-posterior parallel to the curvature of the cingulate, suggesting a specific ordering (see Fig. 3b). Therefore, we tested whether the spatial order of medial prefrontal patches predicted within-chain fMRI signal lags, following^102^. To improve temporal lag detection, ultra-short repetition time (TR 0.46 seconds) functional connectivity data were collected in an individual (27 scanning sessions totaling 169 minutes). Consistent with ordered processing, medial prefrontal chains exhibited posterior-to-anterior ordering of infra-slow activity (<0.1 Hz), significant in 3 out of 4 chains (three one-way ANOVAs testing for main effect of region: *P*s < 0.001; fourth ANOVA: *P* = 0.5; tests Bonferroni corrected for multiple comparisons; Extended Data Fig. 11a). Previous studies have linked lags in infra-slow (<0.1 Hz) activity with coordinating signals enabling propagation of higher-frequency delta activity (0.5 - 4 Hz) in the opposite direction^102^ (see Extended Data Fig. 11b). Thus, these results are consistent with higher-frequency activity moving from anterior to posterior patches within medial prefrontal chains, consistent with the known propagation of signals transforming intentions to actions^103,104^. In data with longer repetition times, the same posterior-to-anterior infra-slow signal ordering (anterior-to-posterior in delta) was also present across all medial prefrontal chains in all participants (PFM data: *r* =-0.49, *P* < 0.001; Extended Data Fig. 11c; TRs = 1.1 – 2.4 seconds). Other chains with a similar curvilinear appearance—those running from the occipito-parietal to superior parietal cortex, following the putative dorsal visual stream^105^—exhibited anterior-to-posterior infra-slow fMRI signal temporal ordering (Supplementary Fig. 14), suggesting posterior-to-anterior propagation of high-frequency neural activity, consistent with electrophysiological recordings preceding movement^106^.

### Functional chain development

Functional connectivity chains may reflect category-, modality, or behavioral objective-specific processing streams. Thus, they may require exposure and activity-dependent tuning in order to segregate from neighboring chains^107^, and should therefore emerge on an extended timescale after birth (as in^6^). Midline prefrontal chains were not visible in a newborn (Fig. 5a, left) and were possibly just beginning to emerge in an 11-month-old infant (Fig. 5a, 2^nd^ from left), but were robust in a 9-year-old (Fig. 5a, 3^rd^ from left). Similar differences were observed with other cortical seeds. Notably, chains in the 11-month-old were more evident in motor and visual regions, including chains seeded from putative fusiform face area (Supplementary Fig. 15), consistent with the early development of facial recognition abilities^108^ and category-selective ventral temporal responses^109^. Whole-brain catenation maps (Fig. 5b) confirmed the relative absence of functional chains in the newborn, followed by increasing catenation in infancy, while the 9-year-old child already showed largely adult-like catenation. Statistical comparisons of catenation maps across participants verified extensive differences between the adult and the newborn and infant, but only small differences between child and adult brains (Fig. 5c for vertexwise two-tailed two-sample *t*-tests, thresholded at *P* < 0.001 vertexwise and cluster-size corrected to *P* < 0.05). Thus, functional chains begin to emerge in the first year of life and are refined during childhood.

**Figure 5.**
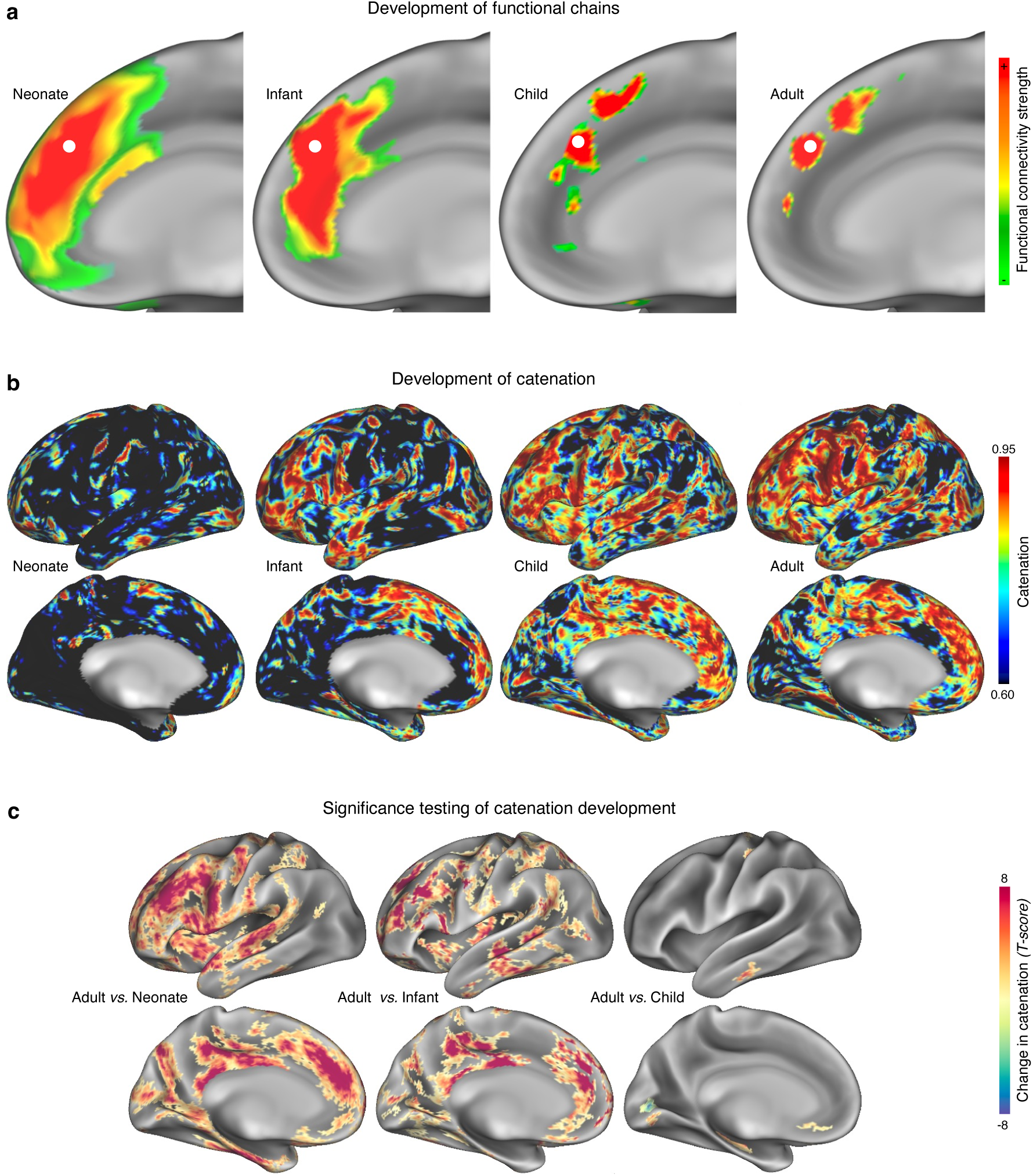
Development of connectivity chains. a, Neonate (37 days old, left), infant (11 months old, left middle), child (9 years old, right middle), and adult (21 years old, right), functional connectivity seeded from medial prefrontal cortex. In the neonate and infant there is no functional chain detectable, in contrast to the child and adult. **b,** Maps of catenation score in the same example participants as in **a**. Catenation was greatly reduced in the neonate and infant compared to the child and the adult. **c,** Statistical testing of the catenation score map changes with age (from **b**), using two-tailed, two-sample vertexwise *t*-tests. Compared to adults, catenation was significantly reduced in the neonates and infant, but not in children. *T*-values are thresholded at *P* < 0.001 vertexwise and cluster-size corrected to *P* < 0.05.

### Chains subdivide and link distinct architectonic areas

Anatomically, the cerebral cortex is divided into discrete cytoarchitectonic areas^22^. However, stimulus-selective patches in temporal cortex, which correspond to functional connectivity chains (Fig. 4), form subdivisions within multiple neighboring architectonic areas (see refs. ^3,54^ for category-selective visual patch vs cytoarchitectonic area comparison in macaques; Supplementary Fig. 16 for how visual patches align with architectonic areas in the present data). Thus, we considered whether subdivision of architectonic areas by functional connectivity chains represents a general principle of brain organization, such that distinct patches of the same functional chain fall into different architectonic areas.

To test this idea, we compared connectivity chains in each participant against the most complete extant map of human architectonic areas, which delineates 140 discrete architectonic areas and currently covers 77% of the cortex^72^. Architectonic areas often contained patches of multiple adjacent chains (see Fig. 6a for an example participant).

**Figure 6.**
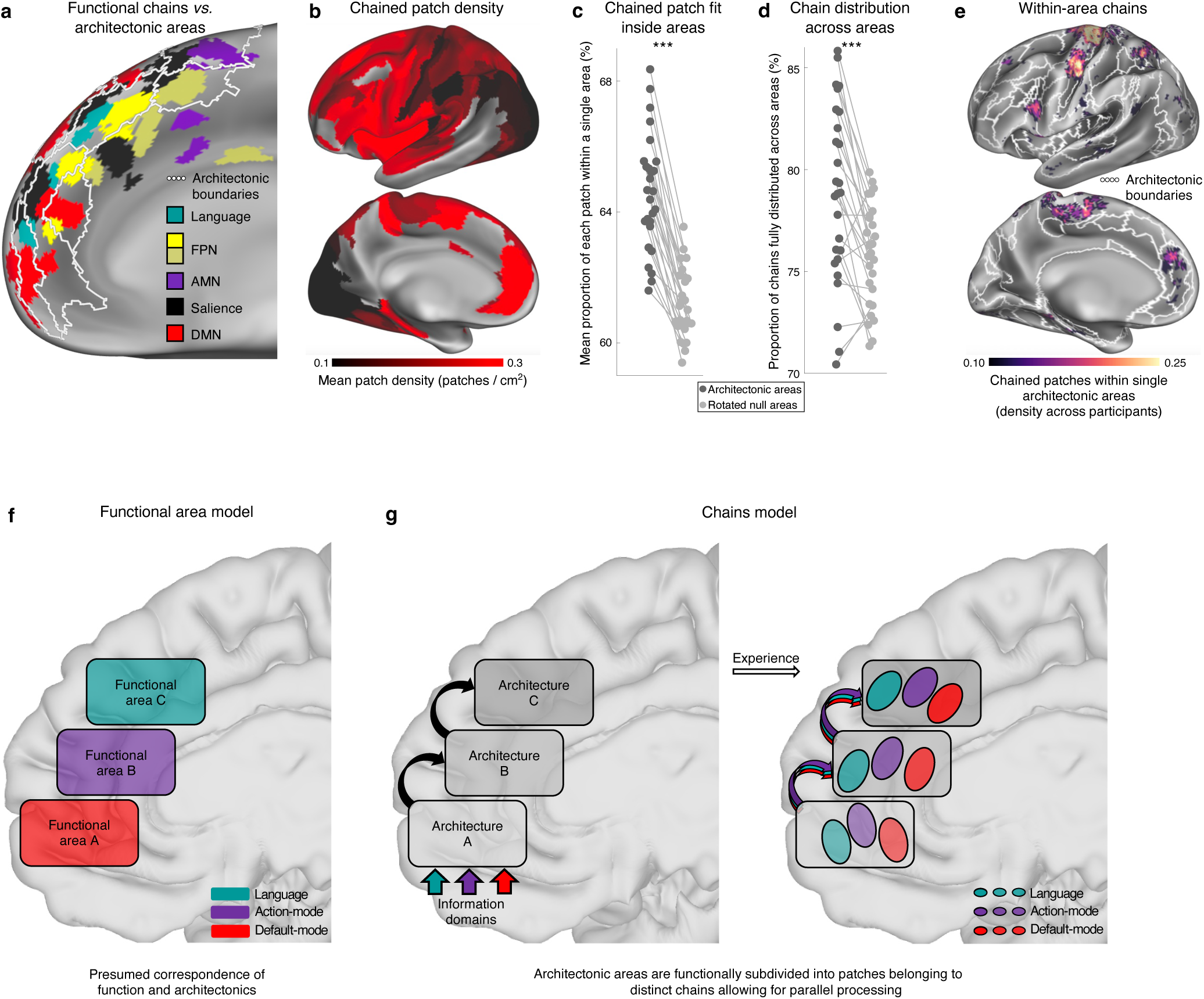
Connectivity chains subdivide and are distributed across architectonic areas. a, Spatially separate patches within connectivity chains (colors) in medial frontal cortex (exemplar participant P1) represent regions within different a priori architectonic areas (white borders; from^72^). **b,** Average density of chained patches within each a priori architectonic area. Areas in frontal cortex contained more densely packed chains of patches. Blank portions of the map indicate as-yet-uncharted regions in which architectonic boundaries have not yet been described as in^72^. **c,** Across participants, patches on average fit better into single cytoarchitectonic areas (dark gray dots: average patch-to-area fit within each participant) than they did into null rotated areas (light gray dots); ***: *P* < 0.001. **d,** Patches within a chain were more distributed across different a priori cytoarchitectonic areas than would be expected by chance. Across participants, connectivity chains were more distributed over architectonic areas (dark gray dots: percent of cross-area chains within each participant) than would be expected by chance (light gray dots: null rotated architectonic areas). ***: *P* < 0.001. **e,** Across individuals, chains that had multiple patches within a single architectonic area (white borders; from^72^) were most commonly located in precentral gyrus and converged with the somato-cognitive action network^6^. **f,** Prior model of cortical organization held by us and many others. Functionally homogenous architectonic areas (colored boxes) process a uniform domain of information and belong to distinct large-scale networks. **g,** Proposed updated model of cortical organization articulating the relationship between functional chains and architectonic areas. Left: each architectonic area (gray box) conducts a specific set of processing operations. These operations are each applied to multiple inputs (colored arrows) carrying information in different domains and then fed forward to the next area in a processing stream. Right: With repeated exposure, architectonic areas subdivide and functionally specialize to operate in parallel on different information domains. Functionally specialized sub-areal divisions are linked across areas, forming connectivity chains.

Across cytoarchitectonic areas and individuals, we observed an average of 2.5 ± 1.9 (mean ± SD) patches per area. Patch density was highest in frontal cortex (Fig. 6b), consistent with the topography of catenation (Fig. 2b). On average, patches were ∼10x denser in frontal than primary cortex, and 2-3x denser in frontal than ventral temporal or superior parietal cortex.

Patches within a chain generally fit within single architectonic areas (Fig 6a). Across participants, a mean ± SD of 64.5 ± 1.8% of each patch fit within a single architectonic area (Fig. 6c); in every individual this fit was better than in null model versions of architectonic areas (paired t-test: *t*(26) = 15.3, *P* < 0.001). This fit could never be perfect due to the inability to define cytoarchitectonic divisions separately within each individual.

Importantly, patches within a chain were distributed such that each patch fell into a different architectonic area (true for mean ± SD = 79.4 ± 4.1% of chains; Fig. 6d). Across individuals, chained patches were more distributed across architectonic areas than in null model versions of architectonic areas (paired t-test: *t*(26) = 4.89, *P* < 0.001). Functional connectivity chains thus seem to link subdivisions of different architectonic areas, with each chain patch potentially carrying out a different type of computational step^28^, likely in an ordered series (Extended Data Figs. 10,11). Adjacent functional patches within the same architectonic area, but belonging to different chains, likely carry out similar computations, but in different information domains or categories and towards different processing and behavioral objectives^61^. Only a few cortical locations had architectonic areas that contained multiple patches of a single chain, mostly SCAN within precentral gyrus^6^ (Fig. 6e).

### Human cortex is functionally organized by chains, not areas

As a general principle of human brain organization, association cortex (and particularly prefrontal cortex) is tiled by interlocking chains consisting of small functional patches.

Larger architectonic areas are functionally subdivided by these chains, containing patches from multiple large-scale networks (Fig. 6g). Patches within a given architectonic area belonging to different chains likely carry out similar computational processes—for which the area’s cellular circuitry is optimized^28,54^—but apply them for different purposes and feed their outputs forward to different targets, as demonstrated in the visual^3,47,57,91,110^ and limbic^111^ systems. These findings expand on recent rodent research demonstrating that cortical activity is organized more by its connectivity than by traditional cytoarchitectonics^63^.

Understanding patches as functional subdivisions of architectonic areas, chained into parallel processing streams, helps explain their spatial distribution on the cortical sheet. Architectonic areas emerge by the discretization of initially continuous cortical gradients^24–26^. Since the originating gradient is continuous and linear, running from high to low concentration of morphogens^24,25^, the discretized architectonic areas must be contiguous rather than distributed. As a result, a processing stream that requires multiple operations enabled by different cytoarchitectonic circuitries cannot be contiguous (Fig. 6f) but must be distributed (Fig. 6g).

These insights help resolve a longstanding disconnect between structural and functional measures of cortical organization^64^. A pervasive, but not universal^54^, assumption of systems neuroscience was that fMRI-derived signals should converge with traditional architectonic areas (Fig. 6f), and thus may be suitable for noninvasively defining those areas^39–43^. However, functionally derived areal parcellations have never properly aligned with architectonic divisions^64^, often subdividing uniform architectonic areas such as primary motor cortex into somatotopic (and non-somatotopic) functional divisions^6,40,112^ or describing split parcels that seem implausible matches for contiguous architectonic areas (e.g., area 55b^41^). Task-derived functional regions similarly fail to converge with architectonics^54^. In individuals, functionally derived parcellations identify many more (almost two-fold) putative functional areas than could be present in architectonics^44,65^, suggesting that function may subdivide (Fig. 6g) rather than converge with architectonics^54^. This framework aligns with evidence in mice that interconnected^113^ and functionally similar^114^ neurons are distributed across architectonic areas^63^. In humans, up to two-thirds of neurons within local brain regions do not participate in active epileptic seizures^115^, while micro-sleeps selectively engage subsets of neurons at small spatial scales that are synchronized across larger scales^116^, suggesting the presence of segregated chain-like local circuits synchronized with distant regions.

Few chains failed to span multiple architectonic areas (Fig. 6d). The highest concentration of chains that violated this principle was in the SCAN^6^ regions of precentral gyrus (Fig. 6e). This could indicate that the SCAN has a unique relationship between structure and function not well explained by the catenation framework, or that primary motor cortex (BA 4) has unrecognized architectonic divisions spanned by the SCAN. The latter possibility is supported by findings of topographically variable intracortical myelin content within Area 4^6,117^.

The strong functional connectivity between patches of the same chain may be supported by the superficial white matter system (Extended Data Fig. 7). First described in 1878^118^, the superficial white matter fibers, also known as short association fibers^87^ or U-fibers^88^, course directly underneath association cortex and link regions whose close proximity does not require long-distance projection into deep white matter fascicles^86^.

Dissection, radiographic tracing, and diffusion MRI in humans and macaques have identified chains of multiple superficial white matter fibers running from occipital to inferior temporal cortex^119,120^ that likely connect category-selective visual processing chains^1,47,91,121^ (Fig. 4). Similar chains of superficial white matter fibers have been described across the brain^122–124^, including those termed *plis de passage* that colocalize with the SCAN chain^125^.

### Chains represent parallel streams for distinct processing objectives

Functional connectivity chains represent temporally ordered processing streams that help parallelize cognitive functions across diverse information domains. Visual patch systems conduct an ordered series of distinct operations on a specific category of stimuli^3,46,57,126^. The discovery of functional chains throughout association cortex suggests that most higher-order cognition operates on similar principles, such that each patch applies an architecture-specific computational step to achieve an overall processing objective.

Chains are functionally segregated from each other within cytoarchitectonic areas (ref. ^54^, Supplementary Fig. 8), which may help minimize unwanted interference between parallel, simultaneous processes. In temporal cortex, such an organization allows ultra-rapid detection (20 ms for faces^127^) and simultaneous recognition of object types within the same visual scene^90^. In auditory cortex it enables simultaneous processing of inputs for speech and non-speech content^128^. This is akin to two assembly lines with shared machinery producing different products within the same factory, such as plastic toys and plastic cutlery.

The processing objectives accomplished by chains likely encompass many of the major functions of association cortex. Here we demonstrate convergence between connectivity chains and processing of faces (Fig. 4), objects, scenes, and pain (Extended Data Fig. 9), as well as circuits for whole-body movement (Fig. 1a)^6^. Prior work has also identified distributed, chain-like sets of regions responsive to a wide variety of domain-or objective-specific tasks. These include prefrontal, parietal, and insular regions selective for visual categories^61,62^, stimulus modality (visual vs auditory working memory)^75,76^, language function^74,129^, memory processing^130^, episodic memory vs social cognition^74,77^, somatotopic tuning^78^, and cognitive control^74^. Chains in non-visuotopic visual cortex are selective for foveolar vision^131^. Distributed, functionally coupled regions linking motor and premotor cortices evoke naturalistic facial gestures^132^. Structurally-coupled chain-like functional patches have also been described in the macaque limbic system^111^. Task-driven activity patterns often exhibit a nearly identical spatial topology to chains highlighted here^61,74,76,133^, as well as being strongly functionally interconnected^6,70,74,129,133^, indicating close correspondence to functional connectivity chains.

The conceptualization of functional chains as domain-and objective-specific ordered processing streams provides the key insight into why the patches are strongly functionally connected to each other across cytoarchitectonic boundaries. Both faces and objects, for example, likely require detailed featural processing within the inferior temporal lobe. Each architectonic area is optimized for a particular type of computation by its laminar organization and distribution of cell types^29^. When a portion of an area becomes specialized to provide featural processing for faces, it should functionally dissociate from other portions of the area providing featural processing for other objects (e.g. bodies, as in^47,134^), but associate with a face-specific subdivision of an adjacent area enacting the next stage of the ordered processing stream. This preferential coactivation, enabled by preferential anatomical projections^91,111,121^, produces temporally correlated chains of focal activity that cross architectonic boundaries.

### Experience-and use-driven functional chain development

Chains are absent from neonatal brains but begin to emerge in the first year of life and are approximately adult-like by age 9 (Fig. 5, Supplementary Fig. 15). This is consistent with work suggesting that functional connectivity in neonates is highly dissimilar to that of adults, exhibiting spatially nonspecific signal spread around any seed region, but that connectivity becomes more adult-like by age two^6,42,135^. Interestingly, neonatal functional connectivity may more closely reflect cytoarchitectonic areas than adult connectivity, as neonatal proto-areas have not yet been functionally subdivided into chains. The timing of initial catenation is also consistent with the first emergence of visual category-selective brain responses^109,136^.

The process of subdividing a uniform cytoarchitectonic field into functional connectivity chains may be experience-and use-driven. In the visual system, category-selective patches do not develop without exposure to the object class^107^, and intensive exposure to new object classes can induce formation of new patches^137^. Category-selective sharpening of neural response curves with stimulus exposure^138^ may be the circuit-level mechanism by which chains segregate from each other. Concomitant with this segregation, white matter fibers underlying temporal cortex patches segregate from being architectonically organized to being chain-specific^121^. A similar process may help shape functional connectivity chains throughout most of association cortex, with exposure to categories of function required for chain development. As an example, segregation of the precentral gyrus into effector-specific motor regions and the chain of SCAN patches, which occurs during the first year^6^, may be scaffolded by early movements during infancy and may ultimately enable intentional body movement for goal directed behavior^139^. Chains representing cognitive or emotional processing may emerge in frontal cortex^61,74,76,111^ with exposure to complex stimuli and situations.

### Chain-specific striatal loops for outcome-based feedback

Functional connections from all patches of a chain converge on a single location within striatum, especially in the caudate (Extended Data Fig. 6c,d). This discovery is consistent with tract-tracing work describing a many-to-one organization of cortico-striatal projections^34,35^ and supports the concept of striatum as serving an integrative, funneling role^30,140^. Parallel cortico-striato-thalamo-cortical loops projecting through striatum and recurring through thalamus^141,142^ are thought to provide outcome-related feedback for thoughts and actions^30^. Precision mapping indicates that each striatal location is not broadly integrative (Fig 3c,d) but instead seems to provide training signals for a specific functional chain. Stimulation work suggests the thalamocortical outputs of these loops are also chain-specific^143^. Thus, chains could represent a hidden computational layer whose outputs receive reward/outcome prediction error signals for rapid learning^140^.

### Patch-specific cerebellar loops for adaptive control

Cerebellar cytoarchitecture is homogeneous, consisting of the same type of circuit likely computing a common transform^33^, yet it is sharply sub-divided according to function and cortical connectivity^144–149^. Unlike striatum, the cerebellum showed a many-to-many functional connectivity pattern with cortex (Fig. 3c, Extended Data Fig. 6c,d), consistent with the known multiple cerebellar representation of both sensory/motor^150,151^ and cognitive functions^144,145,151^. Cortico-ponto-cerebellar-thalamo-cortical loops are thought to provide adaptive control of ongoing cortical functions^152^. Chains in the cerebellum may therefore indicate that cerebellar adjustments, in contrast to the striatum, modify the outputs of patch-specific processes.

### Cortical chains boost adaptability and breadth of cognitive abilities

It has long been argued that parallel processing streams^9,10^ must be a key feature of human brain organization allowing for particularly complex and flexible function^153^. The use-and experience-driven development of functional chains that subdivide and interlink cytoarchitectonic areas may be a major mechanism for achieving this parallelization.

Postnatal subdivision of cytoarchitectonic areas provides a means for flexibly adapting human abilities to new environmental demands, including categorizing and processing new types of information^140^, without the need to evolve fundamentally new circuit types, while the large primate neocortex provides the physical space to implement many of these segregated chains in parallel. Chains are thus a vehicle for rapidly and flexibly expanding the breadth of cognition by simply growing a larger brain.

Surprisingly, functional chains are three times more prevalent in human prefrontal than temporal cortex (Fig 6b), where patch systems were first discovered^1^. Thus far patches have been associated with category-or domain-selective processing of sensory stimuli^51,53,58^. By contrast, prefrontal cortex is widely held to support the highest-order, most abstract, most flexible cognitive processes^154,155^. This seeming contradiction may be resolved by the recent model proposed by Feldmann Barrett and Miller^140^, which holds categorization to be a core computational strategy of the brain, effectively simplifying and generalizing information across short and long timespans to reduce its dimensionality for complex processing operations. Such categorization-based dimensionality reduction should be implemented most commonly in prefrontal functions^140^.

While patch systems are well characterized in non-human primates^46,53,156^, functional chains may take particular advantage of the rapidly enlarged prefrontal cortex of humans. The flexible function of prefrontal cortex^101,157^ may be enabled by chains, which in temporal cortex can rapidly shift representation coding^55,158^. Thus, chains may also lie at the heart of flexible human specialist skills such as social networking, language, and tool use, while powering our ability to adapt to novel living conditions and technologies. In prefrontal cortex, chains belonging to distinct cognitive domains (e.g., default-mode network^159–161^ vs action-mode network^162^) are often immediately adjacent, suggesting that even disparate seeming functions such as self-referential thought and goal-directed action require similar computations instantiated by subdivisions of the same architectonic areas (Fig. 6).

Therefore, detailed exploration of chains may help better illuminate prefrontal cortex function in general.

At the other extreme, the functional chains framework may shed light on the fundamental differences between catenated (prefrontal) and uncatenated association cortex (angular gyrus, medial parietal cortex, retrosplenial cortex). If chains enable the use of a specific cytoarchitecture in multiple parallel processing streams, then angular gyrus, medial parietal cortex, and retrosplenial cortex may each participate in only a single processing stream, similar to primary cortex. Interestingly, these regions are all functionally connected to the hippocampus, which like sensory organs generates de-novo sequences^163,164^. Their processing of hippocampal inputs may thus have similarities to primary cortex receiving information from sensory organs.

Since chains process distinct information types, they may also represent promising targets for novel neuromodulatory therapies for specific brain disorders. It was recently shown, for example, that hyperconnectivity between the SCAN chain (precentral gyrus) and subcortex serves as a biomarker for Parkinson’s, and that successful re-normalization of SCAN connectivity via neuromodulation improves clinical symptoms^165,166^. Targeting condition-appropriate chain-specific cortico-subcortical circuits (such as those recently described in depression^167,168^) could relieve symptoms by modifying signal propagation within those circuits^169^.

A key question moving forward is the extent to which functional chains and patch systems may represent a universal, potentially scale-free cortical organizational principle. Sub-millimeter chains responsive to color, motion, and spatial frequency are present in early visual cortex^170–172^. Smaller functionally linked chain-like organizational structures termed the superficial patch system have been observed at the columnar scale^173–175^. The large distributed networks present in the human brain^13,14^, which include functionally connected regions widely distributed across lobes, may represent a larger-scale version of the same organizational principle. A common organization across spatial scales could suggest the existence of scale-free developmental and computational principles underlying biological intelligence.

## Acknowledgements

This work was supported by NIH grants NS140256 (EMG, NUFD), MH096773 (NUFD, DAF), MH134966 (CMS, TOL, and EMG), MH129616 (TOL), MH121518 (SM), HD109454 (RFS), MH118362 (JRP), HD088125 (JRP), HD055741 (JRP), MH121462 (JRP), MH116961 (JRP), MH129426 (JRP), and MH120989 (CJL); by Michael J. Fox Foundation grant MJFF-026625 (EMG, NUFD); by the National Spasmodic Dysphonia Association (EMG); by the Taylor Family Foundation (TOL); by the Intellectual and Developmental Disabilities Research Center (NUFD); by the Kiwanis Foundation (NUFD); by the Washington University Hope Center for Neurological Disorders (EMG, BPK, NUFD); by the McDonnell Center for Systems Neuroscience (AR); and by Mallinckrodt Institute of Radiology pilot funding (EMG, NUFD).

We would like to thank Doris Tsao, Winrich Freiwald, and Nancy Kanwisher for very helpful comments provided on drafts of this work.

## Author Contributions

Conception: E.M.G. and N.U.F.D. Design: E.M.G., T.O.L., and N.U.F.D. Data acquisition, analysis, and interpretation: E.M.G., A.R., R.J.C., A.L., B.A., A.D., C.J.L., S.R.K., P.C., A.W., N.J.B., K.M.S., J.M., A.M., J.R., T.N., J.R.P., H.L., B.P.K., C.L., C-W.W., S.M., R.F.S., C.M.S., S.E.P., M.E.R., T.O.L., and N.U.F.D. Manuscript writing and revision: E.M.G., S.M., R.F.S., C.M.S., T.O.L., and N.U.F.D.

## METHODS

### Participants

#### Individual-specific Precision Functional Mapping (PFM) data – healthy Washington University adults (single-echo)

##### Participants

Data were collected from three healthy, right-handed, adult participants (ages 35, 25, and 27; 1 female) as part of a study investigating effects of arm immobilization on brain plasticity (data previously published in). Two of the participants are authors (NUFD and ANN). Written informed consent was obtained from all participants. The study was approved by the Washington University School of Medicine Human Studies Committee and Institutional Review Board. Written informed consent was provided by each participant. The primary data employed here was collected either prior to the immobilization intervention (Participants 1, 3) or two years afterwards (Participant 2). This data is available on OpenNeuro^176^ (https://openneuro.org/datasets/ds002766/versions/3.0.2) For details concerning data acquisition and processing, see ^177^; this processing was followed precisely except that data were smoothed with a Gaussian kernel of σ = 1.7 mm (FWHM = 4mm). After controlling for motion, participants retained 338-356 minutes of data.

### Individual-specific PFM data – healthy and post-TBI Washington University adults (multi-echo)

#### Participants

Data were collected from six healthy adult participants with no history of neurological injury (ages 24-64, 4 female). Five of these participants underwent five separate scanning visits within 8 weeks. The sixth underwent 25 scanning visits over 26 weeks.

Data were also collected from four adult participants (ages 21-32, two female) beginning 1-3 weeks after a hospital admittance for a complicated mild Traumatic Brain Injury (Glascow Coma Score^178^ between 13 and 15, with indication on clinical MRI scan) and continued for up to 26 weeks post-injury, with visit frequency up to weekly. The four participants were scanned over 26, 18, 16, and 7 visits. Effects observed in the present study were not influenced by time since injury.

The study was approved by the Washington University School of Medicine Human Studies Committee and Institutional Review Board. Written informed consent was provided by each participant.

### MRI acquisition

In each visit, the participant was scanned using a Siemens Prisma 3T scanner on the Washington University Medical Campus. Each visit included collection of a high-resolution T1-weighted MP-RAGE image (TE = 1.81 ms, TR = 2500 ms, flip angle = 8°, 208 slices with 0.8 mm isotropic voxels) and T2-weighted image (TE = 564 ms, TR = 3200 ms, flip angle = 120°, 208 slices with 0.8 mm isotropic voxels).

Either two 12 minute 30 second long runs (TBI patients, one control) or five 10 minute long runs (five controls) of resting-state fMRI were collected using a multi-echo blood oxygen level-dependent (BOLD) contrast sensitive gradient echo-planar sequence (flip angle = 68°, resolution = 2.0mm isotropic, 72 slices, TEs = [14.2ms, 38.9ms, 63.7ms, 88.4ms, 113.1ms], TR = 1761ms, multiband factor = 6, IPAT = 2). In each session, three pairs of spin echo EPI images (TR = 8.0s, TE = 66ms, flip angle = 90°) with opposite phase encoding directions (AP and PA) but identical geometrical parameters were acquired to correct spatial distortions in the BOLD data.

### MRI processing

Data for all processing and analysis were converted to nifti using dcm2niix^179^.

#### Structural processing

For each participant, FSL FAST was applied to each of the T1-and T2-weighted images to standardize tissue contrast and segmentation (Zhang et al., 2001). All of a participant’s T1 images were linearly cross-registered to each other and averaged across sessions to create a mean T1 image using 4dfp tools (https://readthedocs.org/projects/4dfp/); this procedure was then repeated for the T2 images. The mean T1 image was then linearly registered to MNI atlas space using 4dfp tools. The mean T2 image was linearly registered to the mean T1 image and through to MNI space.

Generation of cortical surfaces from the MRI data followed a procedure similar to that previously described in ^146^. First, anatomical surfaces were generated from the participant’s average T1-and T2-weighted images in native volumetric space using FreeSurfer’s default recon-all processing pipeline (version 7.0^180,181^). This pipeline conducted brain extraction, segmentation, generation of white matter and pial surfaces, inflation of the surfaces to a sphere, and surface shape-based spherical registration of the participant’s original surface to the fsaverage surface ^180,181^. The fsaverage-registered left and right hemisphere surfaces were brought into register with each other using deformation maps from a landmark-based registration of left and right fsaverage surfaces to a hybrid left-right fsaverage surface (‘fs_LR’; ^23^). These fs_LR spherical template meshes were input to a flexible Multi-modal Surface Matching (MSM) algorithm using sulcal features to register templates to the atlas mesh ^182^. These newly registered surfaces were then down-sampled to a 32,492 vertex surface (fs_LR 32k) for each hemisphere.

These various surfaces in native stereotaxic space were then transformed into MNI space by applying the previously calculated T1-to-atlas transformation.

#### Functional processing

For each echo in each run, both phase and magnitude BOLD images were collected and were input into the NORDIC denoising procedure, which enhances image quality by removing zero-mean, unstructured thermal noise, thereby improving the SNR without sacrificing spatial precision^183,184^. Thermal noise levels were implemented as 1/sqrt(2).

Slice-time correction was then applied to each echo using 4dfp tools. Echoes were averaged and movement parameters were computed from the echo-average timecourse. The bias field was computed from the first echo and was removed from all echoes using FSL FAST, and a temporal average BOLD image was computed from this slice-time corrected, motion-corrected, debiased timeseries. Susceptibility distortion fields were computed from the collected spin-echo field maps using FSL TOPUP, and this field was used to undistort the temporal average BOLD image. The undistorted temporal average image was then linearly registered to the participant’s session-average T2 image.

Subsequently, the motion parameters, the undistortion warp, the BOLD->T2 registration, the T2->T1 registration (from *Structural Processing*, above), and the T1->MNI registration (above) were concatenated to resample each echo timeseries into MNI space in a single interpolation using FSL’s applywarp tool. The five MNI-space echoes were then optimally combined using the weighted summation approach^185^, i.e.,

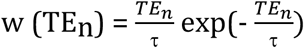

where *TE_n_* is time of the n^th^ echo and time constant *τ* matches expected relaxation time 𝑇^∗^. The echo-combined MNI-space timeseries was then mode-1000 normalized using 4dfp tools.

Additional preprocessing steps to reduce spurious variance unlikely to reflect neuronal activity were executed as recommended in ^186,187^. First, motion estimates were re-computed from the original (pre-slice time correction) timeseries. These motion estimate time courses were filtered to retain effects occurring below 0.1 Hz in order to eliminate “pseudomotion” induced by breathing-related motion of the chest altering the B0 field ^188^. Motion contaminated volumes were then identified by frame-by-frame displacement (FD). Frames with FD > 0.08mm were flagged as motion-contaminated.

After computing the temporal masks for high motion frame censoring, the data were processed with the following steps: (i) demeaning and detrending, (ii) linear interpolation across censored frames using so that continuous data can be passed through (iii) a band-pass filter (0.005 Hz < f < 0.01 Hz) without re-introducing nuisance signals ^189^ or contaminating frames near high motion frames.

Next, the filtered BOLD time series underwent a component-based nuisance regression approach ^146^. Nuisance regression using time series extracted from white matter and cerebrospinal fluid (CSF) assumes that variance in such regions is unlikely to reflect neural activity. Variance in these regions is known to correspond largely to physiological noise (e.g., CSF pulsations), arterial pCO2-dependent changes in T2*-weighted intensity and motion artifact; this spurious variance is widely shared with regions of interest in gray matter. We also included the mean signal averaged over the whole brain as a nuisance regressor. Global signal regression (GSR) has been controversial.

However, the available evidence indicates that GSR is a highly effective de-noising strategy ^186,190^.

Nuisance regressors were extracted from white matter and ventricle masks, first segmented by FreeSurfer ^191^, then spatially resampled in register with the fMRI data. Voxels surrounding the edge of the brain are particularly susceptible to motion artifacts and CSF pulsations ^192,193^; hence, a third nuisance mask was created for the extra-axial compartment by thresholding the temporal standard deviation image (SD > 2.5%), excluding a dilated whole brain mask. Voxelwise nuisance time series were dimensionality reduced as in CompCor ^194^, except that the number of retained regressors, rather than being a fixed quantity, was determined, for each noise compartment, by orthogonalization of the covariance matrix and retaining components ordered by decreasing eigenvalue up to a condition number of 30 (max eigenvalue / min eigenvalue > 30). The retained components across all compartments formed the columns of a design matrix, X, along with the global signal, its first derivative, and the six time series derived by retrospective motion correction. The columns of X are likely to exhibit substantial co-linearity. Therefore, to prevent numerical instability owing to rank-deficiency during nuisance regression, a second-level SVD was applied to XXT to impose an upper limit of 250 on the condition number. This final set of regressors was applied in a single step to the filtered, interpolated BOLD time series, with censored data ignored during beta estimation. Censored frames were then excised from the data for all subsequent analyses.

Nuisance-regressed BOLD fMRI volumetric timeseries were sampled to each participant’s original mid-thickness left and right-hemisphere surfaces (generated as the average of the white and pial surfaces) using the ribbon-constrained sampling procedure available in Connectome Workbench 1.5 ^195^. This procedure samples data from voxels within the gray matter ribbon (i.e., between the white and pial surfaces) that lie in a cylinder orthogonal to the local mid-thickness surface weighted by the extent to which the voxel falls within the ribbon. Voxels with a timeseries coefficient of variation 0.5 standard deviations higher than the mean coefficient of variation of nearby voxels (within a 5 mm sigma Gaussian neighborhood) were excluded from the volume to surface sampling, as described in ^79^. Once sampled to the surface, timecourses were deformed and resampled from the individual’s original surface to the 32k fs_LR surface in a single step using the deformation map generated above (in *Structural processing*). This resampling allows point-to-point comparison between each individual registered to this surface space.

These surfaces were then combined with volumetric subcortical and cerebellar data into the CIFTI format using Connectome Workbench, creating full brain timecourses excluding nongray matter tissue. Subcortical (including accumbens, amygdala, brainstem, caudate, hippocampus, pallidum, putamen, and thalamus) and cerebellar voxels were selected based on the FreeSurfer segmentation of the individual participant’s native-space average T1, transformed into MNI space, and manually inspected. Finally, the BOLD timecourses were smoothed with a geodesic 2D (for surface data) or Euclidean 3D (for volumetric data) Gaussian kernel of σ = 1.7 mm. After controlling for motion, participants retained 131-475 minutes of data.

### Individual-specific PFM data – healthy Cornell University adults

#### Participants

Data were collected from four healthy adult participants (ages 29, 38, 24, and 31; all male) as part of a previously published study ^196^. Two of the participants are authors (CJL and JDP). The study was approved by the Weill Cornell Medicine Institutional Review Board. Written informed consent was provided by each participant. A portion fo this data is available on OpenNeuro^176^ (https://openneuro.org/datasets/ds005118/versions/1.0.0).

For details concerning data acquisition and processing, see ^196^. As above, data were smoothed with a Gaussian kernel of σ = 1.7 mm (FWHM = 4mm). After controlling for motion, participants retained 306-614 minutes of data.

### Individual-specific PFM data – healthy Stanford University adult

#### Participant

Data were collected from a single adult male (age 45 at study onset) who is generally healthy aside from mild psoriasis and a history of anxiety disorder as part of a previously published study^44^. The University of Texas Office of Research Support reviewed the procedure for collecting the primary subject data and determined that it did not meet the requirements for human subjects research as defined by the Common Rule (45 CFR 46) or FDA Regulations (21 CFR 50 and 56), and thus, institutional review board (IRB) approval was not necessary. Transfer of these data to Washington University for analysis was performed with the approval of the Washington University IRB. This data is available on OpenNeuro^176^ (https://openneuro.org/datasets/ds000031/versions/2.0.2).

For details concerning data acquisition and processing, see^44^. As above, data were smoothed with a Gaussian kernel of σ = 1.7 mm (FWHM = 4mm). After controlling for motion, the participant retained 816 minutes of data.

### Individual-specific PFM data – healthy Sungkyunkwan University control

#### Participants

Data were collected from a single healthy adult male (age=39; author C.W.) scanned repeatedly over 28 scanning visits. The study protocol was approved by the institutional review board (IRB) of Sungkyunkwan University. Written informed consent was provided by the participant.

#### MRI acquisition

In each visit, the participant was scanned using a 3T Siemens Prisma scanner in the Center for Neuroscience Imaging Research at Sungkyunkwan University. 26 of these visits included collection of one of resting-state fMRI run as either an 8 minute 13 second (in four sessions) or a 6 minute 22 second (in 22 sessions) BOLD gradient echo-planar sequence (flip angle = 44°, resolution = 2.7mm isotropic, 56 slices, TE =27.2ms, TR = 460ms, multiband factor = 8), as well as two pairs of spin echo EPI images (TR = 7.22s, TE = 73ms, flip angle = 90°) with opposite phase encoding directions (AP and PA) but identical geometrical parameters were acquired to correct spatial distortions in the BOLD data.

After the resting-state BOLD run, each session included collection of fMRI during a pain stimulation experiment. For this, a randomized 45-47° Celsius (0.5° Celsius increments) thermal nociceptive stimulus was applied to the volar surface of the left forearm and the target temperature was maintained for 7 seconds (ramp-up/down=2.5 seconds). Each session included two pain stimulation runs of approximately 12 minutes; across runs, the stimulation site was moved to a different position on the volar surface of the left forearm to minimize sensitization. Each run comprised 12 pain trials, and each trial consisted of 20 seconds of a short film, followed by a pre-stimulation period lasting between 4-6 seconds, then a pain stimulation lasting 12 seconds, followed by a post-heat period of 3-5 seconds, and finally a 5 second period in which the participant rated the pain intensity of the stimulus (range 0.0 [not painful] – 1.0 [maximally painful]). The inter trial period was 4 seconds.

Finally, the second scanning session also included collection of a high-resolution T1-weighted MP-RAGE image (TE = 2.34 ms, TR = 2400 ms, flip angle = 8°, 224 slices with 0.7 mm isotropic voxels).

#### MRI processing

Data for all processing and analysis were converted to nifti using dcm2niix^179^.

#### Structural processing

FSL FAST was applied to the T1-weighted image to standardize tissue contrast and segmentation (Zhang et al., 2001). The T1 image was linearly registered to MNI atlas space using 4dfp tools. Generation of cortical surfaces from the MRI data followed the procedure described in “Individual-specific PFM data – healthy and post-TBI Washington University adults” above.

### Functional processing

Slice-time correction was applied to each run using 4dfp tools, and movement parameters were computed from the resulting timecourse. The bias field was computed and removed from the data using FSL FAST, and a temporal average BOLD image was computed from this slice-time corrected, motion-corrected, debiased timeseries.

Susceptibility distortion fields were computed from the collected spin-echo field maps using FSL TOPUP, and this field was used to undistort the temporal average BOLD image. The undistorted temporal average image was then linearly registered to the T1 image. Subsequently, the motion parameters, the undistortion warp, the BOLD->T1 registration, and the T1->MNI registration (from *Structural Processing*, above), were concatenated to resample each echo timeseries into MNI space in a single interpolation using FSL’s applywarp tool. The MNI-space timeseries was then mode-1000 normalized using 4dfp tools.

For resting-state data, filtering, motion control, and nuisance regression were implemented following the procedures described in “Individual-specific PFM data – healthy and post-TBI Washington University adults” above. After controlling for motion, the participant retained 169 minutes of resting-state data. All data was then sampled to the cortical surface and into the CIFTI format following the procedure described in “Individual-specific PFM data – healthy and post-TBI Washington University adults” above.

#### Task analysis

Individual trials were classified as being painful or not painful based on the pain ratings (trials rated below 0.2 out of 1.0 were classified as not painful). Painful trials, non-painful trials, and movie visualization were each modeled as separate conditions with FSL FEAT^197^ using a gamma convolution (phase=0 seconds, standard deviation=3 seconds, mean lag=6 seconds). Vertexwise beta estimates for each condition were estimated in each run. A paired t-test across all runs contrasted the beta values estimated in the painful vs the nonpainful conditions.

### Individual-specific PFM data – healthy University of Minnesota controls (Natural Scenes Dataset)

#### Participants

Data were collected from eight healthy young adults (6 female; age 19-32 years) scanned repeatedly over 39-49 scanning visits as part of a previously published study^90^. The study was approved by the University of Minnesota Institutional Review Board.

Written informed consent was provided by each participant.

For details concerning data acquisition and some processing, see^90^. This data is publicly available (https://registry.opendata.aws/nsd/). Here, we downloaded 1) processed and Freesurfer-segmented structural data, 2) fully analyzed BOLD functional localizer maps, and 3) minimally processed resting-state BOLD timecourses (slice-time corrected, motion-corrected, and registered to structural image).

#### MRI processing

Freesurfer surfaces were registered to the fs_LR 32k space, and functional localizer maps were sampled to the cortical surface and into the CIFTI format, all following procedures described in “Individual-specific PFM data – healthy and post-TBI Washington University adults” above. Denoising and surface-processing of the resting-state BOLD data was conducted as described in^168^. After controlling for motion, participants retained 76-169 minutes of resting-state data.

### Individual-specific PFM data – Neonates

#### Participants

Data were collected from eight sleeping, healthy full-term neonatal participants (average postmenstrual age (PMA) 42.7 weeks). The study was approved by the Washington University School of Medicine Human Studies Committee and Institutional Review Board. Written informed consent was provided by a parent.

#### MRI acquisition

Participants were scanned while asleep over the course of 2-5 consecutive days using a Siemens Prisma 3T scanner on the Washington University Medical Campus. Every session included collection of a high-resolution T2-weighted spin-echo image (TE = 563 ms, TR = 4500 ms, flip angle = 120°, 208 slices with 0.8 x 0.8 x 0.8 mm voxels). In each session, resting-state fMRI runs were collected using various scanning parameters across each of the individual neonates (Table S1). The number of BOLD runs collected in each session depended on the ability of the neonate to stay asleep during that scan; in total, an average of 99.5 minutes of data were collected in each neonate. A pair of spin echo EPI images with opposite phase encoding directions (AP and PA) but identical geometrical parameters and echo spacing were also acquired during each session.

#### MRI processing

Structural and functional processing followed the pipeline described in^43^. The primary difference from the adult datasets is that segmentation, surface delineation, and atlas registration were conducted using a T2-weighted image (the single highest quality T2 image, as assessed via visual inspection) rather than a T1-weighted image, due to the inverted image contrast observed in neonates.

### Individual-specific PFM data – Infant

#### Participant

Data were collected from one healthy sleeping infant age 11 months. The study was approved by the Washington University School of Medicine Human Studies Committee and Institutional Review Board. Written informed consent was provided by a parent. For details concerning data acquisition and processing, see^6^.

### Individual-specific PFM data – Children

#### Participant

Data were collected from twelve healthy children ages 9-11. The study was approved by the Washington University School of Medicine Human Studies Committee and Institutional Review Board. Written informed consent was provided by a parent and assent was given by the participant.

#### MRI acquisition

Participants were scanned over the course of two to four sessions using a Siemens Prisma 3T scanner on the Washington University Medical Campus. Data collection was identical to that described in “Individual-specific PFM data – Treatment-Resistant Depression (TRD) Washington University adults” above, except each session included collection of 2-4 BOLD fMRI runs, which were collected as either 10 minute or 16 minute sequences during movie watching rather than during rest. All sequence parameters were otherwise identical.

#### MRI processing

All processing was identical to that described in “Individual-specific PFM data – Treatment-Resistant Depression (TRD) Washington University adults” above. After controlling for motion, participants retained 77-151 minutes of data, retaining 54% to 90% of collected data.

### Group datasets

#### Adolescent Brain Cognitive Development (ABCD) Study

Twenty minutes (4 x 5 minute runs) of resting-state fMRI data, as well as high-resolution T1-weighted and T2-weighted images, were collected from 3,928 9-10 year old participants (51% female), who were selected as the participants with at least 8 minutes of low-motion data from a larger scanning sample. Data collection was performed across 21 sites within the Unites States, harmonized across Siemens, Philips, and GE 3T MRI scanners. See ^81^ for details of the acquisition parameters. Data processing was conducted using the ABCD-BIDS pipeline (https://github.com/DCAN-Labs/abcd-hcp-pipelines); see ^198^ for details.

#### Human Connectome Project (HCP)

One hour (4 x 15 minutes run) of resting-state fMRI data were collected from 1200 healthy adults ages 25-37 years old. See^79,80^ for data acquisition details. Minimally processed volumetric resting-state data (including motion correction, fieldmap-based unwarping, and alignment to MNI atlas space) were downloaded. Additional processing steps were applied to this data following^186,187^, which included regression of subject specific white matter, ventricle and global signal average signals, as well as removal of motion artifact by first filtering the estimated motion parameters to remove respiration-associated pseudomotion and then excluding frames with filtered framewise displacement greater than 0.04mm^188^. At this stage 185 participants were excluded from further analysis due to either excessive motion^188^ or the presence of radiofrequency coil artifact (as detailed here: https://wiki.humanconnectome.org/docs/HCP%20Subjects%20with%20Identified%20Quality%20Control%20Issues%20(QC_Issue%20measure%20codes%20explained).html), leaving a final sample size of 1015 participants. Processed data was then sampled to the individual-specific cortical surface and into the CIFTI format following the procedure described in “Individual-specific PFM data – healthy and post-TBI Washington University adults” above.

### Analyses

#### Functional connectivity computation

For each single-participant dataset, a vertex/voxelwise functional connectivity matrix was calculated for each individual from the resting-state fMRI data as the Fisher-transformed pairwise correlation of the concatenated timeseries of all vertices/voxels in the brain. Separately in the PFM, ABCD, and HCP datasets, vertex/voxelwise group-averaged functional connectivity matrices were constructed by first calculating the vertex/voxelwise functional connectivity within each participant as the Fisher-transformed pairwise correlation of the timeseries of all vertices/voxels in the brain, and then averaging these values across participants at each vertex/voxel.

#### Detection of chained functional connectivity patterns (catenation)

Connectivity chains have the appearance of at least two strong connectivity peaks separated by a region of lower connectivity. Thus, we implemented an approach to search for such formations in every individual subject and group average dataset.

First, for every point in the brain, we defined sets of vertices radiating in straight lines (geodesically, along the cortical surface) from that origin point out to a radius of 60mm. These lines represent linear directions along which chains could be present. See Supplementary Fig. 9a for visualization of these lines. 60mm was chosen as a reasonable distance limit to define connected regions as “chains”, given that this distance covers a greater area than any single classically defined cytoarchitectonic region. We also explored 40mm and 80mm distances and found that the localization of chained connectivity was very similar (see Supplementary Fig. 17). Lines were spaced at an angle of 5°, which enabled good coverage of areas distant from the central origin point while reducing overlap for regions proximal to the origin point (Supplementary Fig. 9a).

Second, we defined a set of linear chain-like template connectivity patterns that true connectivity along these lines could match. Each template was modeled as two Gaussians, one centered at the origin point and one centered at some distance from the origin point. With no a priori knowledge of the distance between chained regions or of the size of each chained region, we tested templates that varied the distance between Gaussians (values ranged from 30mm to 60mm) and the sigma value of each Gaussian (values ranged from 6mm to 12mm). See Supplementary Fig. 9c for a visualization of these templates.

Third, the connectivity pattern was computed using the central origin point as a seed (see above “<u>Functional connectivity computation</u>”). For every straight line originating from this seed, functional connectivity values along this straight line were extracted (Supplementary Fig. 9b). Each linear connectivity pattern was then correlated against all of the linear chain-like templates. This produced a large number (# lines X # templates) of correlation values representing possible matches to a chain-like connectivity pattern radiating in some direction from the origin point. In principle, the presence of chain-like connectivity at that origin point could be assessed as the strongest correlation among all tested matches. In practice, we found that the highest correlation value among all line matches was somewhat unstable to parameter variations, due to the susceptibility of correlations with low sample sizes (i.e. few numbers of points) to extreme values. By contrast, the 90^th^ percentile highest correlation match value was much more stable. As such, we assigned the 90^th^ percentile value to the central origin point as the presence of chain-like connectivity seeded from that point.

This procedure was repeated for all cortical vertices, creating a “catenation” map of correlation values representing the presence of chained functional connectivity across the cortex in each individual and group average dataset.

We note that this procedure is optimized for finding chains of two patches in a line. We acknowledge that the existence of a third patch in a straight line within the 60mm limit would actually reduce the fit of the double-gaussian template patterns. In practice, we note that three-patch chains are rarely in a perfectly straight line (as opposed to a curvilinear line, which is more common; see Extended Data Figure 1), and so this is an infrequent fail condition for the procedure. Further, even in this fail condition, catenation can still be detected in lines projecting from the middle patch outward to the outer ones.

#### Statistical testing of catenation maps

*Comparison vs a null model:* The chain detection approach (above) searches for punctate, spatially distributed functionally correlated regions. However, such spatially distributed correlations could in principle also emerge from noise, if correlations that emerge by chance are spatially smoothed. To ensure that the detected catenation maps do not reflect such noise-driven correlations, we developed a null model of catenation consisting of spatially smoothed random correlations. This was accomplished by phase randomizing the unsmoothed data from each vertex in each individual, which destroys any extant structured functional correlations, and then smoothing the resulting phase-randomized data to the same degree as the original data (sigma = 1.7mm). We then repeated the chain detection approach above to create a catenation map from the phase-randomized null data.

To test whether catenation maps created from real data systematically differed from those created from null data, we first converted the catenation values (Pearson’s R) to Z values using Fisher’s Transformation to ensure normality. We then conducted a two-tailed paired t-test across participants at each cortical vertex comparing real to null catenation values.

We corrected these vertexwise tests for multiple comparisons using a cluster-based correction approach, as follows: we first thresholded the map of P values at p < 0.001 uncorrected and computed the sizes of all resulting clusters of significant vertices. We then created a null model for these cluster sizes by randomly permuting the data labels of each map (that is, swapping the real and null labels) 1000 times. In each iteration, vertexwise two-tailed paired t-tests compared the permuted real against the permuted null maps, and cluster sizes were computed after thresholding at p<0.001 uncorrected. This approach establishes a null expectation for significant clusters if significance was solely due to chance. The final cluster size threshold was the cluster size at which fewer than 5% of null iterations had a significant cluster at least that large (i.e., corrected p < 0.05). This value was 17 vertices. The cluster size threshold was then applied to the original paired t-test statistical map.

*Relative prevalence across cortical lobes:* We tested whether catenation maps exhibited higher values in some cortical lobes than others. For each adult subject, we computed the average catenation map value within each cortical lobe (frontal, insula, parietal, temporal, occipital). Paired t-tests compared catenation map values between each pair of lobes, with Bonferroni correction for the 10 tests conducted.

*Relative prevalence across large-scale networks:* We tested whether catenation maps exhibited higher values in some large-scale networks than others. For each adult subject, we defined large-scale brain networks using the infomap graph theoretical community detection approach^13,199^, following procedures described in^6,168^. We then computed the average catenation map value within each of 14 networks. Paired t-tests compared catenation values between each pair of networks, with Bonferroni correction for the 91 tests conducted.

*Individuals vs. group averages:* We tested whether catenation maps computed in adult individuals were systematically different from those computed in group averages. For each group average catenation map, we conducted a vertexwise one-sample t-test in which the catenation values at each vertex from all individual adults were compared against the catenation value at that vertex from the group average. Vertexwise tests were Bonferroni corrected for the 59412 vertices tested.

*Adults vs. pediatric subjects:* We tested whether catenation maps computed in adult individuals were systematically different from those computed in neonates, infants, and children. We conducted a vertexwise two-sample t-test comparing adults to neonates. We then applied a permutation-based cluster-wise correction for multiple comparisons^65,200^, as follows: vertex-wise t-values were thresholded at uncorrected p<0.001, and the sizes of resulting clusters of supra-threshold vertices were computed. Group identities of individuals (adults vs. neonates) were then randomly permuted 1000 times. T-values and supra-threshold clusters were re-computed to generate a null distribution of cluster sizes. Clusters from the true adults vs. neonates comparison survived cluster-based correction if they were larger than any cluster found in 95% of randomly shuffled tests. This approach established a cluster extent threshold of 40 vertices corresponding to an overall corrected level of p<0.05. This approach was then repeated for the adults vs. children comparison, which established a cluster extent threshold of 29 vertices corresponding to an overall corrected level of p<0.05.

As we only had one infant dataset available, we compared the adults to the infant using a one-sample rather than a two-sample t-test. Permutation-based testing was not available with only one individual, so we employed the same cluster-based threshold found to be significant in the adults vs. neonates comparison above.

#### Comparison to superficial white matter

To test whether the superficial white matter system preferentially supported high-catenation cortex, we downloaded the ORG white matter atlas^89^ from (https://github.com/SlicerDMRI/ORG-Atlases). All fiber clusters within superficial tract groupings were first converted to a volumetric fiber density map using the “wm_tract_to_volume.py” script in the White Matter Analysis package provided with the ORG atlas (https://github.com/SlicerDMRI/whitematteranalysis)^89^. A linear registration was computed between the ORG-space population average T1 provided within the atlas and the MNI152 linear template brain. This registration was applied to each superficial fiber cluster density volume, and the resulting MNI-space fiber density maps were summed to create an overall superficial white matter density map.

To compare the (cortical, surface-based) catenation map with the (subcortical, volumetric) superficial white matter map, we mapped catenation values in each cortical vertex to the white matter regions directly beneath that vertex. To accomplish this, we first created interior surfaces 6mm underneath cortex by projecting the vector from the pial to the gray-white surface 6mm into white matter. We then mapped the cross-PFM-participant average catenation map into the volumetric ribbon between the gray-white surface and the 6mm-under surface using Connectome Workbench tools (wb_command-metric-to-volume-mapping)^195^. We computed the average catenation value within the portion of the under-cortex ribbon that overlapped with dense superficial white matter fibers (fiber density > 2). This reflected the catenation value of cortex that was specifically over superficial white matter. Finally, to test whether this catenation value over superficial white matter was higher than would be expected by random chance, we created a null model by randomly rotating the catenation map around the cortical surface 1000 times following^40,201^, thus preserving the spatial distribution of the catenation values while randomizing their location. We then projected each rotated null catenation map into white matter and computed the average catenation value within superficial white matter regions, as above. Significance was assessed as the proportion of null rotated maps that exhibited greater average catenation values within superficial white matter regions than the real catenation map. Notably, we excluded insular regions from this test, as the external capsule white matter tract running directly under insula was not categorized as “superficial” in the ORG atlas.

#### Moran’s I

As an alternative method for quantifying the presence of local chain-like connectivity patterns, we computed Moran’s I^84^ on the connectivity pattern seeded from each vertex. Moran’s I is a measure of spatial autocorrelation that generalizes to a space with any dimensionality. First, we computed the connectivity pattern using each vertex as a seed (see above “Functional connectivity computation”). To characterize local connectivity patterns in particular (e.g., chains in the local area), we restricted the computation of Moran’s I to the connectivity values within 60mm of the seed point (as with the catenation metric). Moran’s I was then computed on this pattern as:

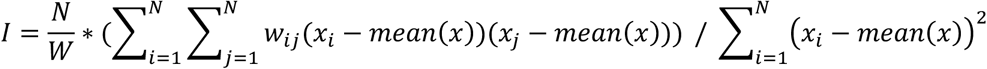

Where x is each connectivity value within 60mm, indexed by *i* and *j*; N is the total number of connectivity values; w is an NxN spatial weights matrix; and W is the sum of all elements in the spatial weights matrix.

The spatial weights matrix describes which elements are considered to have a neighboring spatial relationship. Connectivity maps are intrinsically spatially autocorrelated, but we wanted the Moran’s I value to be sensitive to the presence of distributed connectivity at a scale beyond that baseline autocorrelation. Therefore we used a spatial weights matrix identifying pairs of vertices that were up to fourth-level adjacent (i.e., adjacent to an adjacent to an adjacent to an adjacent vertex) as neighbors in the weights matrix. Practically, varying the level of adjacency in the weights matrix did not strongly affect the spatial topography, but instead had the effect of compressing or expanding the distribution of I values relative to a value of 1 (e.g., including only first-level adjacencies would cause most I values to be between 0.9 and 1.0).

#### Definition of discrete connectivity chains

To define and map each chain of functional connectivity, we entered all vertex-seeded connectivity patterns into the Infomap algorithm^199^, a data-driven graph-theoretic community detection approach. When applied to functional connectivity data, Infomap groups points that are interconnected with each other into communities. We have used this approach in the past to define communities that represent large-scale brain networks^6,13,65,100^. Here, we modify the approach by 1) restricting analyzed cortical vertices to those with a chain match value (see “Detection of chained functional connectivity patterns”, above) at r=0.70 or higher; 2) thresholding connections as the top 1% of connections from each vertex, to define chains only as regions linked by very strong connectivity; 3) clustering each thresholded connectivity map and removing the cluster containing the seed vertex, to prevent connections arising from local autocorrelation of BOLD data; and 4) removing all connections outside of the 60mm distance limit (see above), to restrict defined communities to local connectivity chains only. We explored other thresholding values from 0.2% to 2%, and we found that values below 1% produced increasingly sparse sets of chains, while values at 1% and above produced very similar results (see Supplementary Fig. 18).

#### Connectivity to subcortex

*Chain specificity of subcortical-seeded connectivity:* To test whether subcortico-cortical connectivity also exhibited chain-like patterns, for each subcortical structure among caudate, putamen, thalamus, hippocampus, and cerebellum, we computed a cortical connectivity map in each subject seeded from every voxel in that structure. We then identified the chain (as defined above, in “Definition of connectivity chains”) that exhibited the strongest connectivity to that voxel, in a winner-take-all fashion. If the voxel was more strongly connected to uncatenated brain regions than to any chain, that voxel was excluded from further analysis.

We reasoned that if subcortical-seeded connectivity was chain-like, connectivity seeded from that voxel to that distributed chain would be stronger than connectivity from the voxel to patches from other chains that lie interdigitated between the selected chain’s patches (e.g. a less chain-like, more blob-like connectivity pattern). As such, we computed the average connectivity between that voxel and all patches in all other chains interdigitated between the patches of the most-connected chain.

To avoid circularity, we conducted this analysis in a cross-validated fashion.

Specifically, each individual’s BOLD data was divided into two halves, and the identity of the most-connected chain was defined from the first half of the data, while computation of connectivity in the most-preferred patches and the in-between chain patches was conducted in the second half.

We then averaged the within-chain connectivity strength across the voxels within the subcortical structure, and averaged the between-chain connectivity strength across the voxels within that structure. Finally, in each structure we conducted a paired t-test across subjects testing for significant differences between the within-vs between-chain connectivity strength, with Bonferroni correction for the five tests conducted.

*Effect of catenation level on subcortical connectivity:* To explore whether catenated regions exhibited different connectivity into subcortex than uncatenated regions, in each individual in the PFM, HCP, and ABCD datasets, we defined high-catenation and low-catenation vertices in the cortex by performing a median split on the subject-specific catenation map. In each individual we then computed the average functional connectivity of all high-catenation vertices, as well as the average connectivity of all low-catenation vertices, with every subcortical voxel. Finally, within each dataset, we computed a voxelwise paired t-test comparing high-catenation vs. low-catenation seeded connectivity in each voxel. Because PFM individuals were not constrained to all have the same subcortical mask, voxels included in the subcortical mask of fewer than 10 participants were excluded from testing. Follow-up paired t-tests within each dataset compared high-vs low-catenation-seeded connectivity within striatum (anatomically defined as the combination of caudate, putamen, accumbens, and pallidum), as well as within hippocampus.

*Chain-like connectivity patterns in subcortical structures:* To test whether connectivity patterns within subcortical structures also exhibited chain-like patterns, for each subcortical structure among caudate, putamen, thalamus, hippocampus, and cerebellum, we examined the voxels most strongly connected in each subject to each chain from the winner-take-all procedure above (in “Chain specificity of subcortical-seeded connectivity”). We first identified discrete clusters of voxels with different chain identities, applying a cluster size threshold of 4 voxels. We then determined whether each cortical chain’s winner-take-all voxels exhibited a chain-like distribution (i.e., multiple distinct clusters within the subcortical structure) or not (only one distinct cluster within the structure). We then computed the proportion of each structure with a chain-like organization (i.e., number of voxels comprising multiple distinct clusters of winner-take-all assignment, divided by total number of voxels in the structure). Finally, we conducted paired t-tests across subjects testing for significant differences between each pair of structures in the proportion of chain-like organization, with Bonferroni-correction for the 10 tests conducted.

#### Distance dependence of within-chain connectivity

To test whether connectivity between within-chain patches was stronger when the patches were closer together—which would indicate some degree of anatomically-dependent ordering within chains, rather than each chain being a mutually interconnected net—we computed the functional connectivity between all pairs of within-chain patches.

We also computed the geodesic distance between the centroids of all pairs of within-chain patches. We then correlated within-chain connectivities against distances, both combining across all participants and within each participant.

To ensure that the connectivity-distance relationships we observed were chain dependent, and not just a general principle of functional connectivity, we also computed connectivity and geodesic distance between all pairs of cross-chain patches. These values were computed only for patch pairs within the same hemisphere, as geodesic distance cannot be computed across hemispheres. We then correlated cross-chain connectivities against distances within each participant. Finally, we tested whether within-chain connectivity was more distance-dependent than cross-chain connectivity. We applied the Fisher transformation to each participant’s within-and across-chain connectivity-vs-distance R values (to improve their normality), and we entered them into a paired t-test comparing within-against cross-chain Z(R) values.

#### Temporal ordering of signals within chains

To test whether a consistent temporal ordering of fMRI signals exists in the anterior-posterior span of chains within dorsal medial prefrontal cortex, we first identified the distinct chains within this region of the brain in every subject (using chains identified from “Definition of connectivity chains”, above). For each run, we then computed time delays between the separate regions within each chain, following the lag detection approach described in^202^. Briefly, vertex timecourses were interpolated into motion-censored frames and temporally upsampled. A lagged cross-covariance function was estimated between every vertex and the mean timecourse of the chain, with a maximum lead/lag of 2.0 seconds. We then computed the temporal offset that resulted in the strongest covariance with the average chain signal; this was taken as the signal lead/lag of that vertex relative to the rest of the chain. Vertex lead/lag values were then averaged within each discrete region of the chain.

One individual subject had data with very high temporal resolution (“Individual-specific PFM data – healthy Sungkyunkwan University control”: TR = 0.46sec), which can be expected to improve the accuracy and reliability of lead/lag estimates. Within this subject, we tested whether regions within each of four chains exhibited significant differences in their lag across scanning runs. Within each of the chains, lead/lag estimates for each region computed from each run were entered into a one-way ANOVA, with the chain region as the factor of interest. Significance was Bonferroni corrected for the four ANOVAs run.

Results from the high temporal resolution individual suggested that anterior regions within each chain exhibited later BOLD signals than posterior regions. To test whether this was broadly true, in every individual we averaged lead/lag estimates for each chain region across runs. For every chain, we then computed the centroid (the point in the physical center) across all chain vertices. For each region within the chain, we then computed the mean distance anterior or posterior from that centroid. Finally, across individuals, we correlated all within-chain lead/lag estimates against the anterior/posterior distance from the chain’s centroid.

#### Chained connectivity within face-responsive patches

We tested whether connectivity was chain-like among face-responsive patches. In individual subjects from the Natural Scenes Dataset (NSD), we defined face-responsive patches in each individual as contiguous clusters of vertices within the “faces” functional localizer map with a t-score > 2.0. This threshold defined clusters of face-responsive vertices that were highly convergent with known patterns of face-responsive patches in prior work^4^ and appeared chained. We then defined FFA in each individual as the cluster closest to the peak location of activity associated with the term “face” within the Neurosynth meta-analytic database (neurosynth.org;^12^), which was found to be at MNI coordinates [+40-48-20]. To determine whether this putative FFA exhibited strong connectivity to these chain-like patches, we computed a connectivity seed map from the FFA and computed the average connectivity to all other face-responsive patches. To test whether FFA connectivity to the other face-responsive patches was stronger than would be expected by chance, we created a null model by randomly rotating the face-responsive patches around the cortical surface 1000 times following^40,201^, thus preserving the spatial distribution of the face patches while randomizing their location. We then computed the connectivity of the FFA to the rotated face patches. Within each subject, significance was assessed as the proportion of null face patches that exhibited stronger connectivity to the FFA than the real face patches.

#### Comparison of chains to a priori architectonic areas

We downloaded the Julich Brain Cytoarchitectonic Atlas v3.1^72^ from (https://julich-brain-atlas.de/atlas/probabilistic-maps). We removed from the atlas all vertices labeled “GapMap” (including Frontal-to-Temporal-I, Frontal-to-Temporal-II, Frontal-to-Occipital, Temporal-to-Parietal, Frontal-I, and Frontal-II GapMaps), which indicate as-yet-uncharted regions that have not yet been divided into architectonic areas.

For all distinct cortical chains in every individual (defined as in “Definition of connectivity chains”, above) we defined the patches of each chain as the discrete contiguous clusters of at least 10 vertices. We then computed the centroid of each patch within each chain. We computed the number of patch centroids within each architectonic area in each individual, as well as the density of patches, in patches per cm^2^, within each area.

For each patch, we determined whether the chained patches fit within the architectonic areas by computing the proportion of each patch that fell within a single architectonic area (the area that contained the centroid). We then determined whether this assessed patch-to-area fit was better than would be expected given randomly located cortical areas. We created a null model by randomly rotating the cytoarchitectonic area map around the cortical surface 1000 times following^40,201^, thus preserving the sizes, spatial distribution, and relative adjacencies of the areas while randomizing their location. For each of the null model rotated area maps, we recomputed the within-area fit of each chained patch within each individual. Significance was assessed across individuals using a paired t-test that compared the average patch fit within real architectonic areas to the average fit (across patches and iterated rotations) within null model areas.

Finally, for each chain, we then determined whether all of the patch centroids were within different cytoarchitectonic areas. We computed the proportion of chains that had all of their patch centroids within different cytoarchitectonic areas. Chains were not considered if they contained fewer than two centroids in defined architectonic areas. We then determined whether the distribution of chains across areas was higher than would be expected given randomly located architectonic areas created by rotating the area map around the surface 1000 times, as above. For each of the rotated area maps, we again computed the proportion of chains that had all of their patch centroids within different (rotated) cytoarchitectonic areas. As above, significance was assessed across individuals using a paired t-test. Here, the test compared 1) the proportion of chains that had all patches within different real architectonic areas to 2) the mean proportion of chains that had all patches within different rotated null areas, averaged across iterated rotations.

