## Extended Data and Supplemental Materials for "Parallel processing chains span cytoarchitectures to organize association cortex"

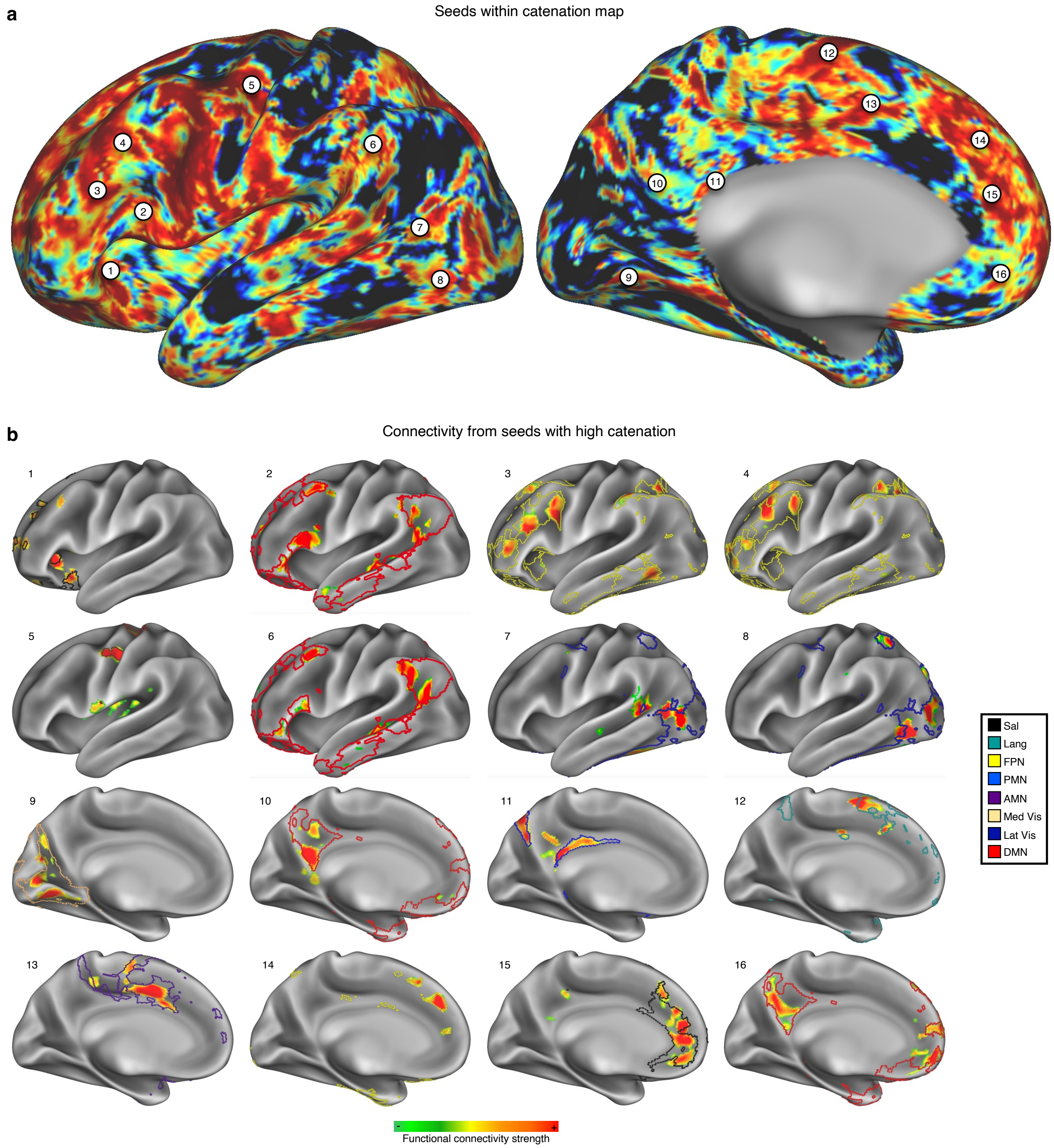


**Extended Data Figure 1| Chains fit within large-scale functional brain networks.** In an example participant (P1), **a,** sixteen cortical seeds were chosen as locations with high catenation. **b,** Connectivity maps generated from each seed in this participant did exhibit chained connectivity. Further, each connectivity chain fit within a large-scale functional brain network (colored outlines). Sal – salience network; Lang – language network; FPN – fronto-parietal network; PMN – parietal memory network; AMN – action-mode network; Med Vis – medial visual network; Lat Vis – lateral visual network; DMN – default-mode network.

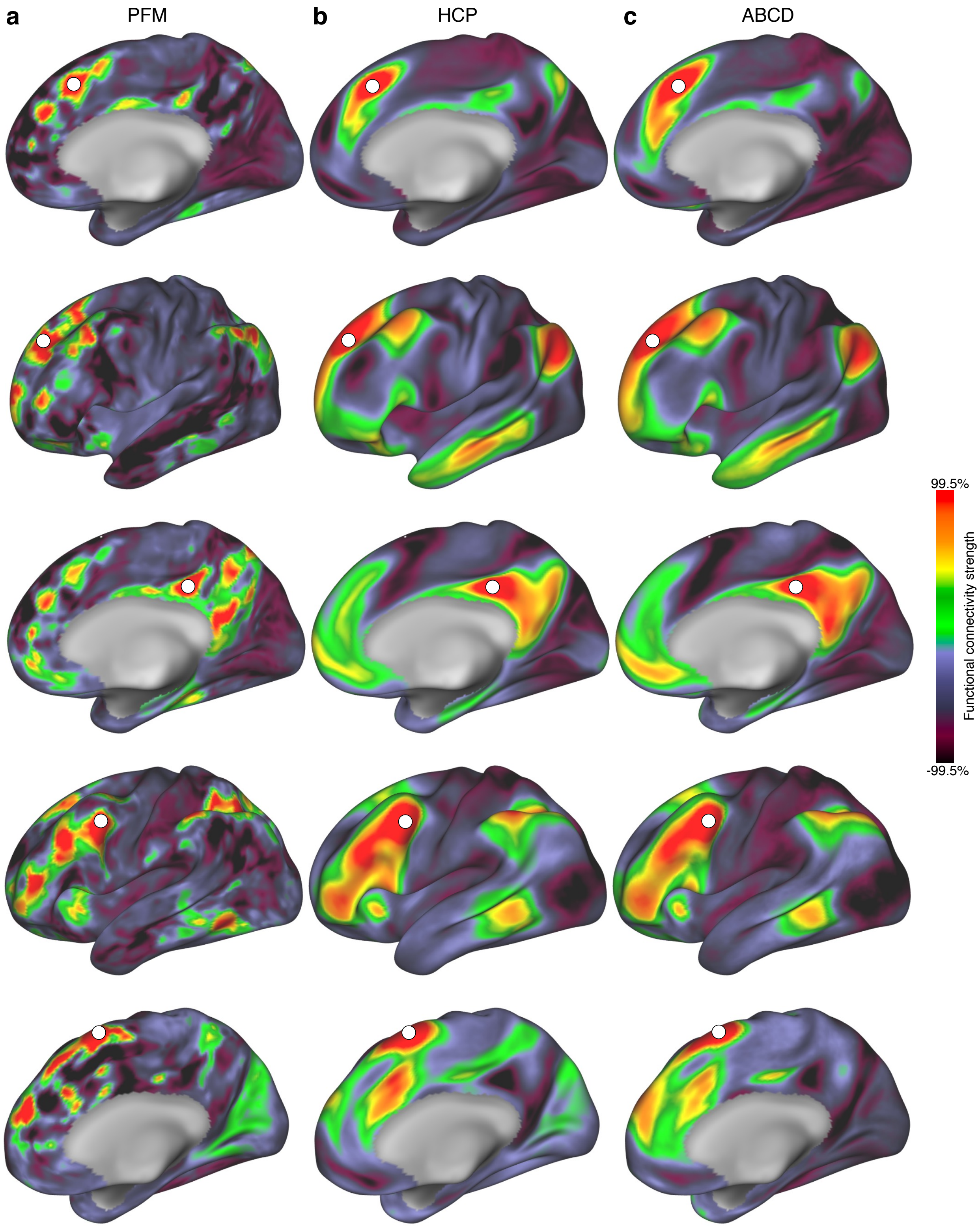


**Extended Data Figure 2| Chained connectivity in individual and group-averaged data.** Five example connectivity maps seeded from the same location in **a,** a single PFM individual (P1), as well as in group-averaged data from **b,** the Human Connectome Project (*n* = 1015) and **c,** the Adolescent Brain Connectivity Development project (*n* = 3928). Each seed elicits a similar overall pattern of connectivity, but chains that are evident in the individual are obscured in the group-averaged data.

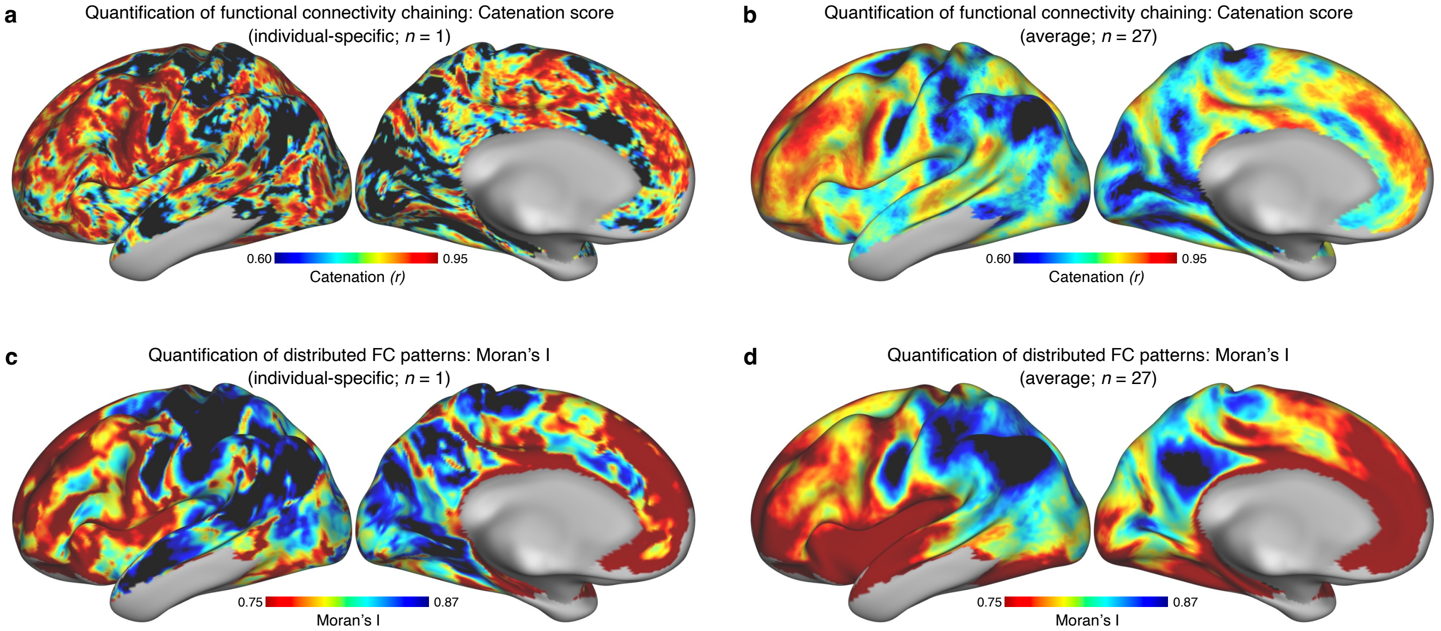


**Extended Data Figure 3| Catenation converges with Moran's I. a,** The catenation map computed from a single individual (P1, as in Fig. 2a) and **b,** catenation maps averaged across individuals (as in Fig. 2b) both converge closely with **c,** a map of Moran's I computed on each vertex's FC map in a single individual (P1) and **d,** Moran's I maps averaged across individuals. To aid interpretation, color scales are reversed in **c-d** relative to **a-b**, because the catenation score increases as FC patterns become more distributed and chain-like, while Moran's I decreases with increasing spatial distribution of FC patterns.

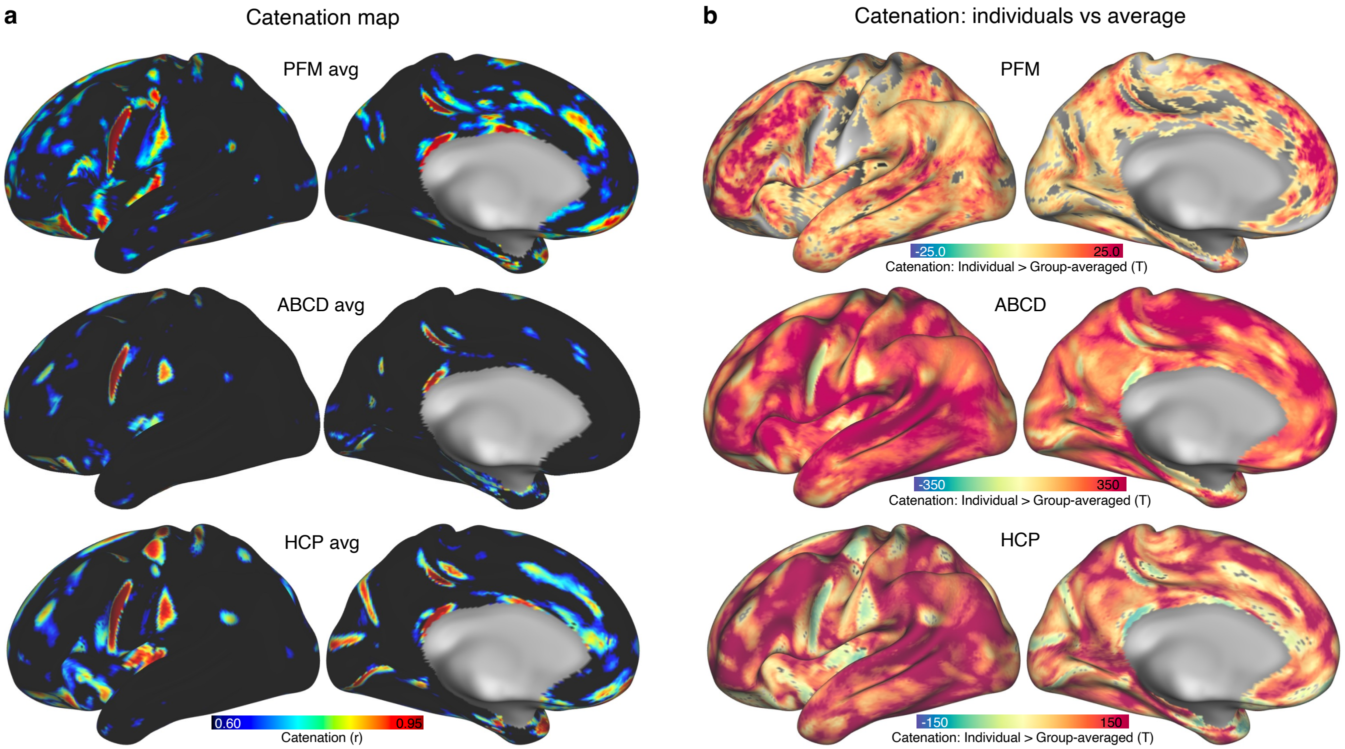


**Extended Data Figure 4| Catenation in group-averaged data**. **a,** Catenation was computed (as in Fig 2) in three group-averaged datasets: the average of the 27 PFM individuals, the Adolescent Brain Cognitive Development data (ABCD), and the Human Connectome Project data (HCP). Relative to individuals (example in Fig 2a), group-averaged data exhibited broadly reduced catenation across much of cortex **b,** T-tests comparing catenation from all individual participants in each dataset against the group-averaged catenation maps from **a**. Individuals exhibited significantly greater catenation across nearly the entire brain than the group averages. Maps are thresholded at $P < 0.05$, Bonferroni corrected for the number of vertices tested.

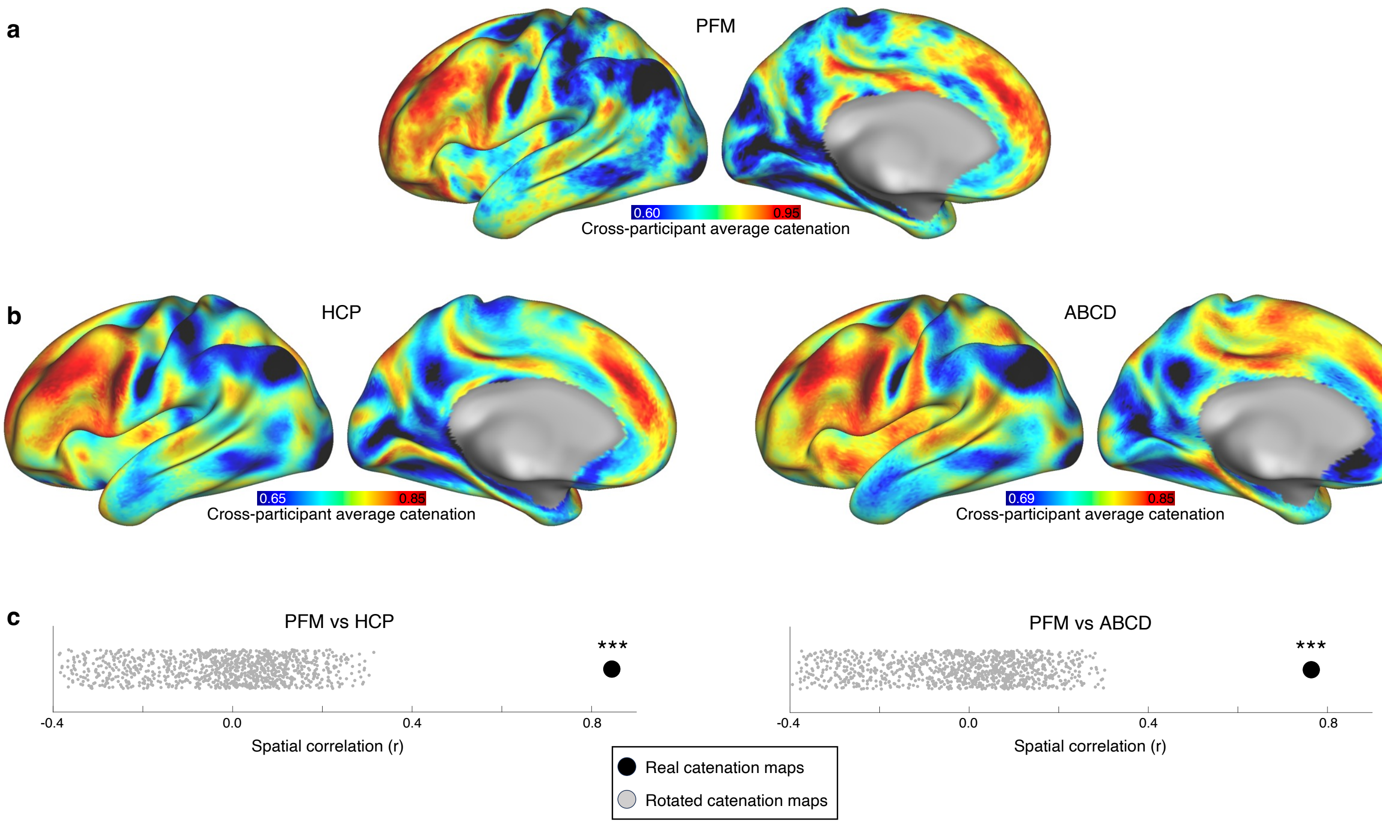


**Extended Data Figure 5| Catenation patterns replicate in large datasets. a,** Average of individual-specific catenation maps in PFM data (from Fig 2b). **b,** Maps generated by computing catenation in individuals and then averaging, without averaging the raw data, for HCP (left) and ABCD (right), rescaled to emphasize spatial similarity to the PFM map. **c,** Similarity (spatial correlation) of the PFM catenation map with the HCP (left) and ABCD (right) catenation maps (black dots). These maps were much more similar than would be expected by chance (null rotated HCP and ABCD catenation maps: grey dots). *** indicates $P < 0.001$.

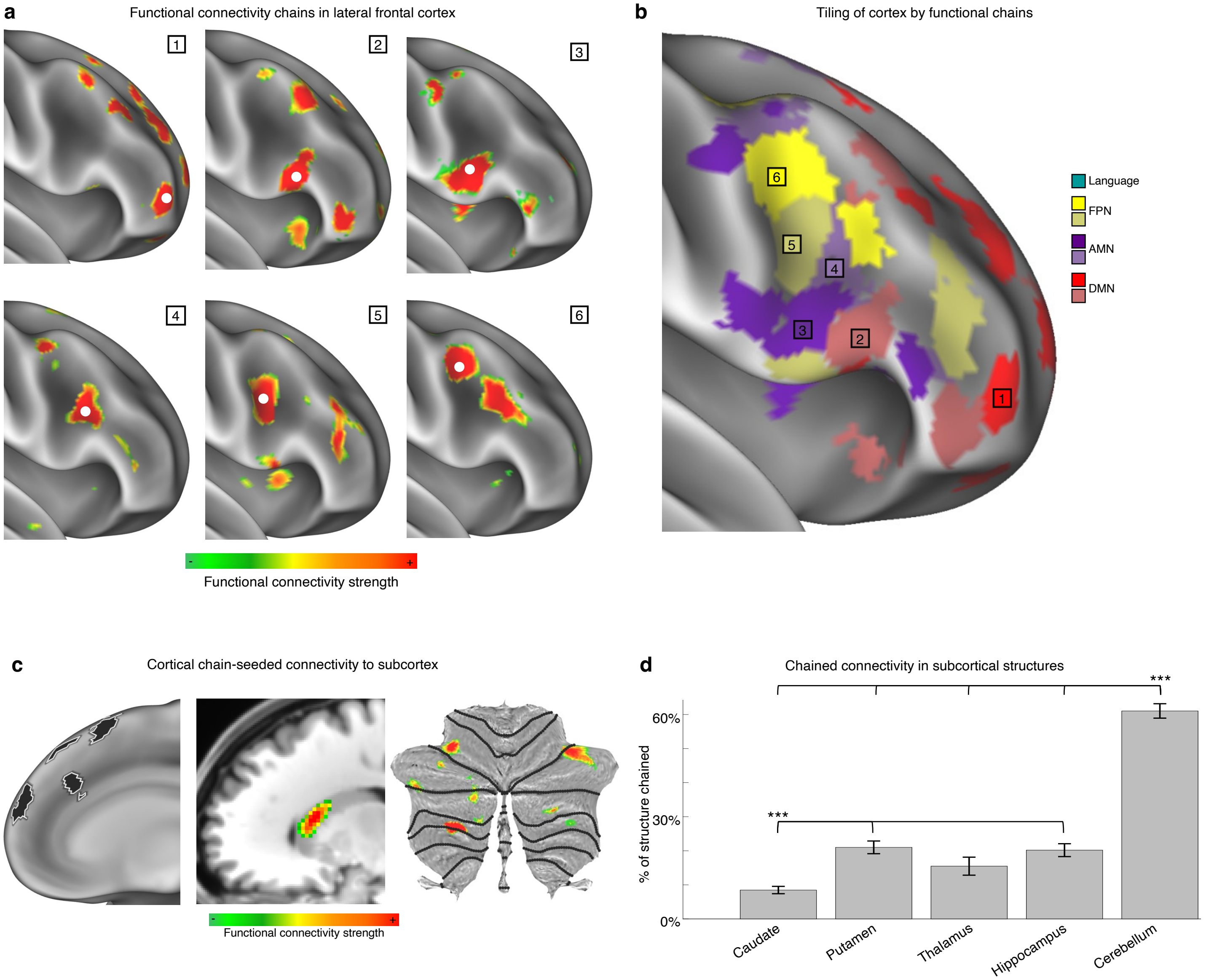


**Extended Data Figure 6| Densely interleaved lateral prefrontal cortex chains and cortico-to-subcortical connectivity. a**, Connectivity seeds (1-6) placed in lateral prefrontal cortex of the exemplar participant (P1) revealed a series of discrete, interleaved functional connectivity chains. **b**, A community detection algorithm (see Methods) defined these chains (1-6) in a data-driven manner, revealing that connectivity chains closely abut each other and together tile most of the lateral prefrontal cortex. Chains are color-coded according to the large-scale canonical functional networks they are part of (language = teal, fronto-parietal network (FPN) = yellows, action-mode network (AMN) = purple, default-mode network (DMN) = red. **c,** Seeding all regions of a cortical chain (left) identified a single location in caudate exhibiting strong functional connectivity (middle) (P1). However, it identified multiple distributed regions exhibiting strong connectivity in cerebellum (right). **d,** Across all cortical connectivity chains in all participants, regions of strong connectivity to a chain frequently constituted multiple chained regions in cerebellum but infrequently constituted multiple regions in caudate, putamen, thalamus, and hippocampus. *** indicates $P < 0.001$, corrected.

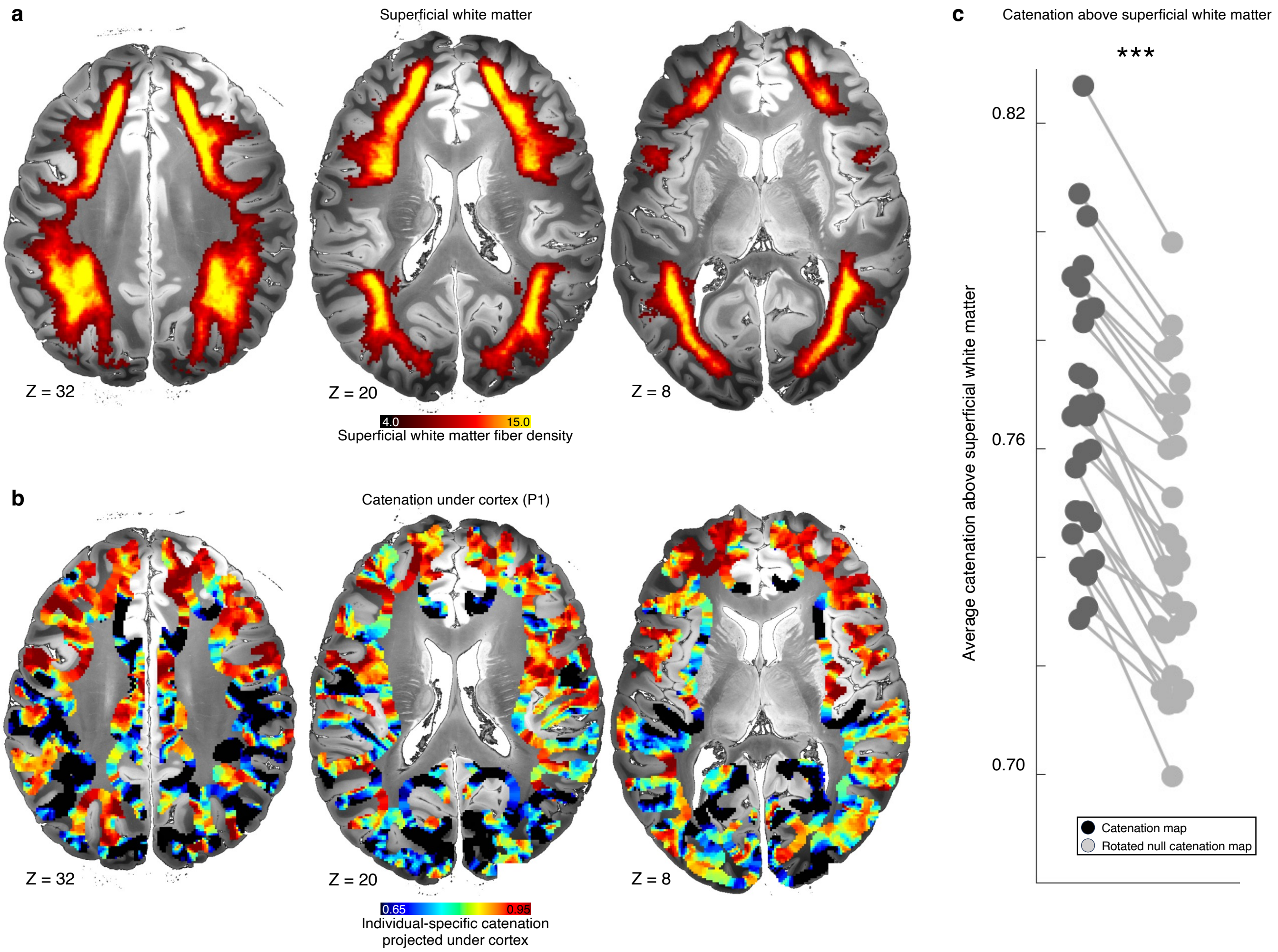


**Extended Data Figure 7| Relationship of superficial white matter tracts to highly catenated cortex. a,** Group-averaged superficial white matter tracts from the ORG atlas (Zhang et al., 2018). **b,** Catenation values from a single exemplar participant (P1) from cortical regions above superficial white matter tracts, projected into the white matter directly below cortex. **c,** In each participant, mean true catenation value above superficial white matter tracts (dark gray dot) relative to mean null model catenation values (rotated around the cortical surface 100 times before projecting into white matter) above superficial white matter tracts, averaged across rotations (light gray dots). Real catenation values were higher than null values in every individual; the difference between the two was significant (*** indicates $P < 0.001$ in a paired t-test).

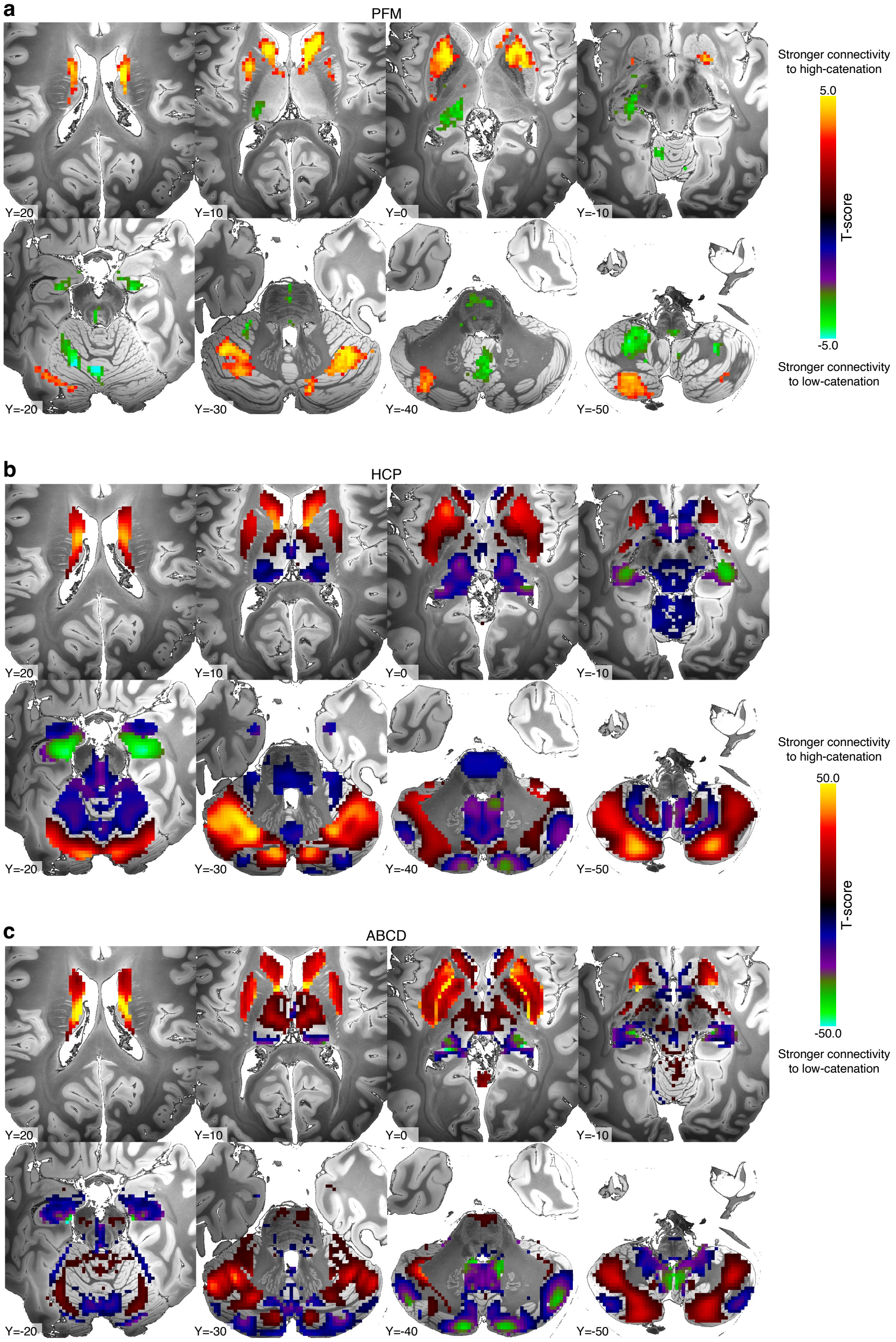


**Extended Data Figure 8| Subcortical connectivity differs by catenation level.** To examine which subcortical regions preferentially exhibited connectivity to catenated vs uncatenated cortex, in **a,** the PFM, **b,** the HCP and **c,** the ABCD datasets we defined high-catenation and low-catenation cortical vertices as a median split-half of each individual's catenation map. In each individual we then computed subcortical connectivity maps seeded from the combination of all high-catenation cortical regions and from all low-catenation cortical regions. Within each dataset we conducted a voxelwise paired *t*-test across individuals between high- and low-catenation-seeded connectivity. Results are thresholded voxelwise at $P < 0.005$ uncorrected for the PFM data and $P < 0.05$, Bonferroni-corrected for the number of voxels tested for the HCP and ABCD data.

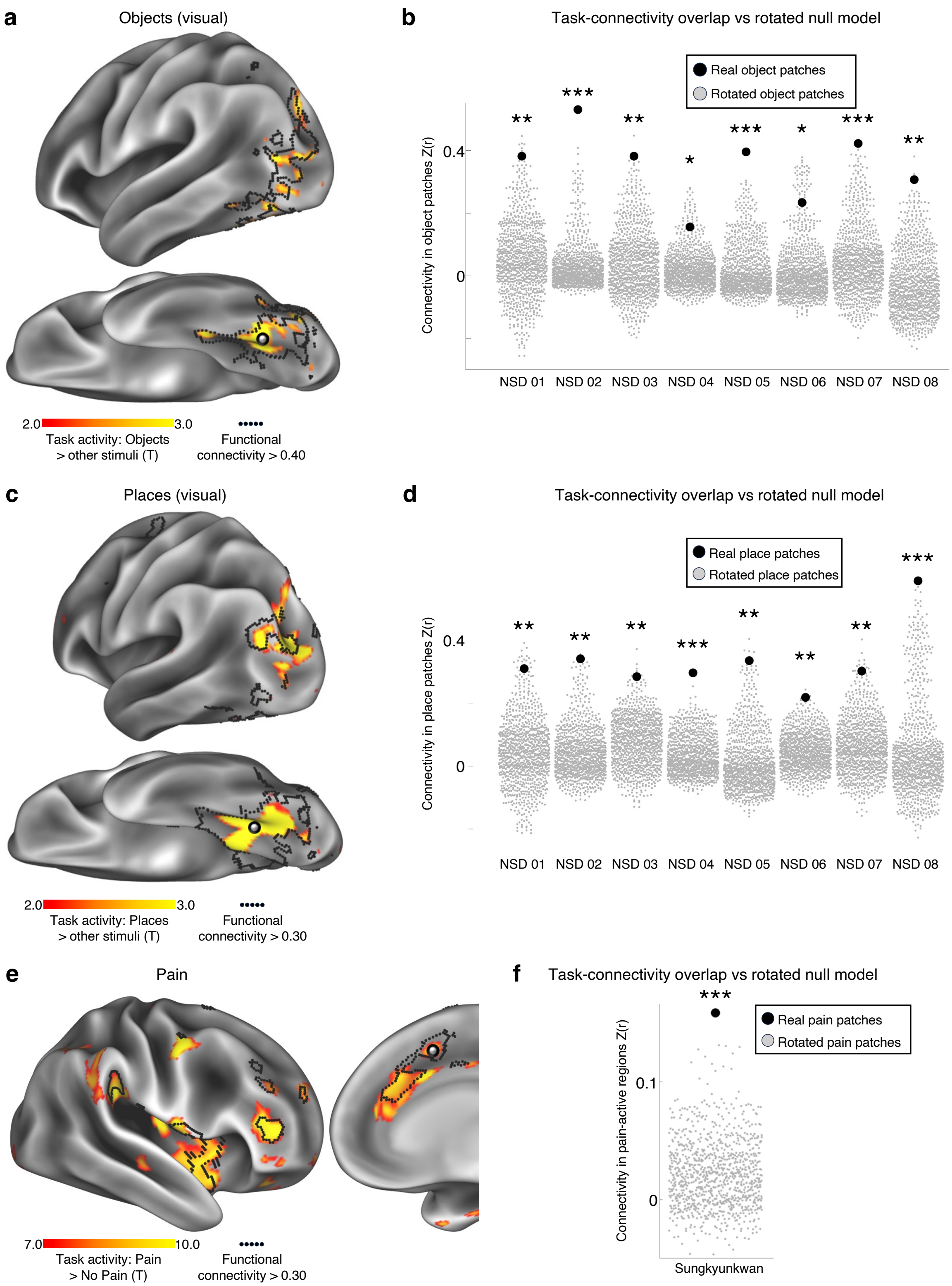


**Extended Data Figure 9| Match between individual-specific precision task fMRI activation maps and functional connectivity patches and chains for additional tasks. a,** In an exemplar individual (NSD05) from the Natural Scenes Dataset (NSD), a task contrast localizing activity related to visual presentation of objects (object stimuli > all other stimulus types; red-yellow) demonstrated a chained pattern of activity in ventral temporal and lateral occipital cortex that overlaps with chained connectivity seeded from the object-active region of ventral temporal cortex (black outline). **b,** Across NSD participants, connectivity seeded from the object-active region of inferior temporal cortex exhibited strong connectivity to other object-active regions (black dots). In all participants, this chained connectivity pattern was more convergent with the object-active regions than would be expected by chance (null rotated object patches: gray dots). **c,** In the same exemplar individual, a task contrast localizing activity related to visual presentation of places (place stimuli > all other stimulus types; red-yellow) demonstrated a chained pattern of activity in ventral temporal and lateral occipital cortex that overlaps with connectivity seeded from the place-active region of ventral temporal cortex (black outline). **d,** Across NSD participants, connectivity seeded from the place-active region of inferior temporal cortex exhibited strong chained connectivity to other place-active regions (black dots). In all participants, this chained connectivity pattern was more convergent with the place-active regions than would be expected by chance (null rotated place patches: gray dots). **e,** In the individual from the Sungkyunkwan dataset, a task contrast localizing activity related to painful heat (painful heat > non-painful heat; red-yellow) demonstrated a chained pattern of activity in dorsomedial and anterior lateral prefrontal cortex that overlaps with chained patterns of connectivity seeded from the dorsomedial prefrontal cortex (black outline). **f,** This chained connectivity pattern was more convergent with the pain-active regions than would be expected by chance (null rotated pain-responsive regions: gray dots). *** indicates $P < 0.001$; ** indicates $P < 0.01$; * indicates $P < 0.05$.

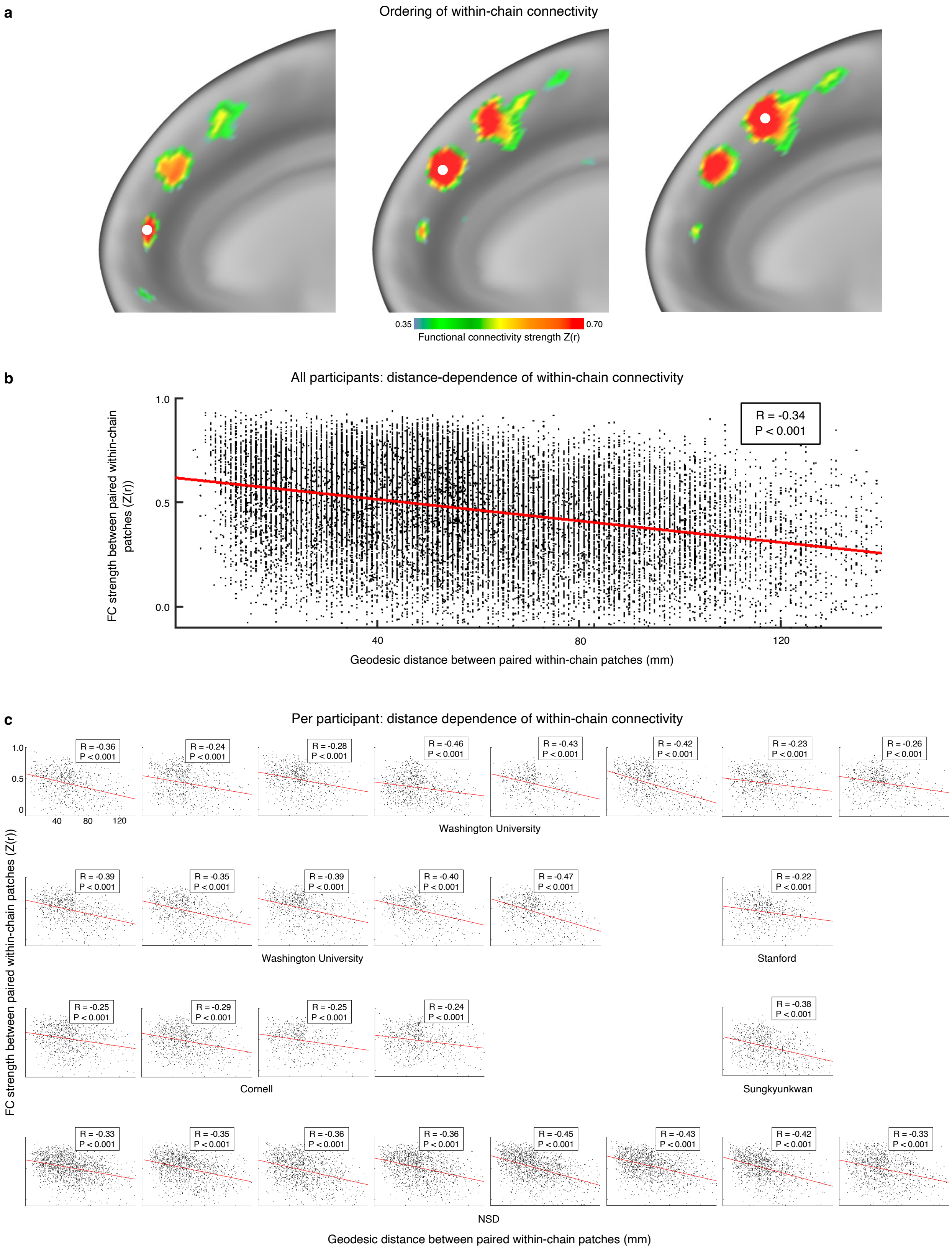


**Extended Data Figure 10| Within-chain connectivity is stronger between closer patches.**
**a,** In an example participant (P1), connectivity seeded from each of three patches of a chain

demonstrated that connectivity was stronger to closer patches of the chain than to more distant patches. **b,** Across all participants and **c,** within each participant, connectivity between all pairs of within-chain patches was stronger when the patches were closer together on the cortical surface. All *P*s < $10^{-6}$, Bonferroni-corrected for multiple comparisons.

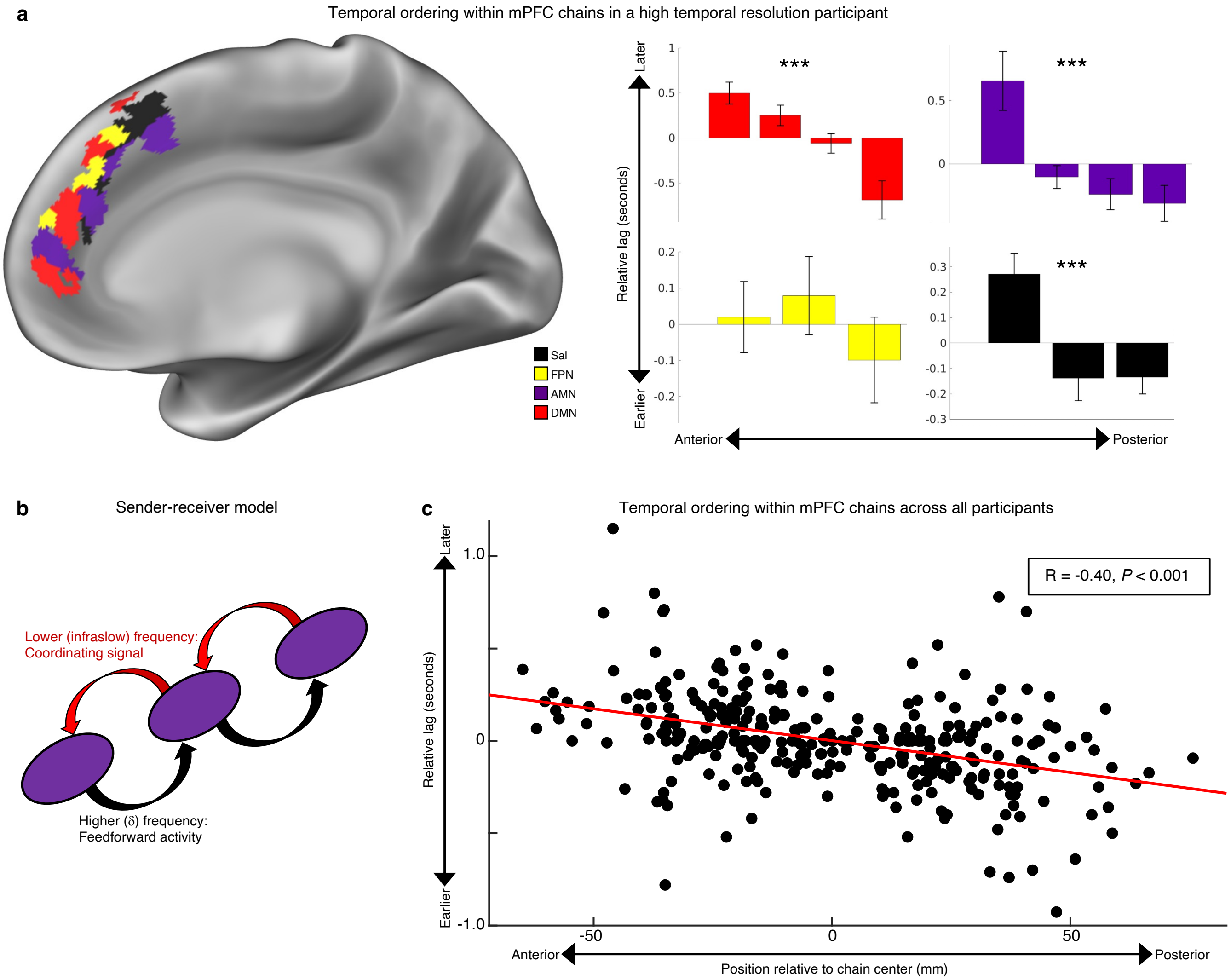


**Extended Data Figure 11| Temporal ordering of signals within medial prefrontal connectivity chains. a,** Lagged connectivity (infra-slow, 0.08-0.1Hz) was estimated within each of four chains identified in the medial prefrontal cortex of a PFM participant with large amounts (169 minutes) of very short-TR data (left). Three of four chains exhibited a significant anterior-posterior ordering, in which signals in more anterior regions of the chain exhibited later signals than those in more posterior regions. The zero-point is arbitrary and indicates timing relative to other regions within each chain. Error bars represent standard error across $n$ = 26 sessions. *** indicates $P$ < 0.001. Sal – salience network; FPN – fronto-parietal network; AMN – action-mode network; DMN – default mode network. **b,** The sender-receiver model for interregional communication[91]. Infra-slow activity (0.08-0.1Hz, detectable by fMRI) propagates from downstream to upstream regions in order to provide a coordinating signal for higher (delta-band, 0.5-4Hz) frequency feedforward signaling representing active processing. **c,** Across all PFM participants, lagged correlations were computed between regions in medial prefrontal cortex chains. Regions of the chain that were more anterior exhibited systematically later signals than those more posterior (Pearson's correlation: $P$ < 0.001). Together, the sender-receiver model suggests that high-frequency activity thus propagates within chains in an anterior-to-posterior direction.

**Supplementary Data**

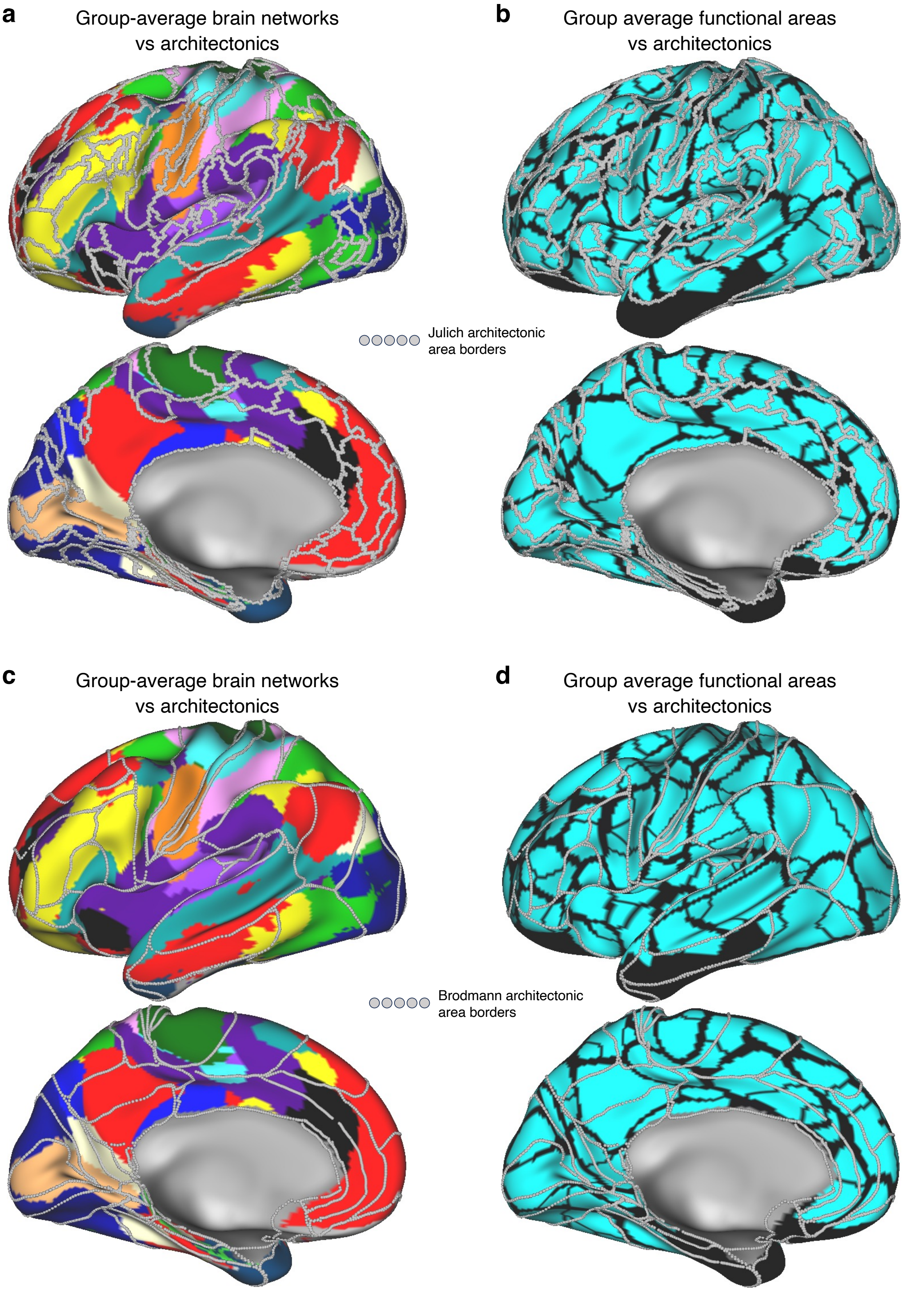

**Figure S1| Relationship between functional and architectonic brain divisions. a,** Overlap between large-scale group-average brain networks (colored brain regions) (Gordon et al. 2017) and modern architectonic area borders (silver) (Amunts et al. 2020). **b,** Overlap between functionally defined areal divisions, identified as locations with strong changes in functional connectivity (black borders) (Gordon et al. 2016) and modern cytoarchitectonic area borders (silver). **c,** Overlap between large-scale group-average brain networks and original Brodmann architectonic area borders (silver) (Van Essen et al. 2012). **d,** Overlap between functionally defined areal divisions and Brodmann architectonic area borders (silver). In all cases, alignment is reasonable in primary (visual, motor) cortex, but borders are strikingly misaligned in association cortex (e.g., lateral and medial prefrontal cortex).

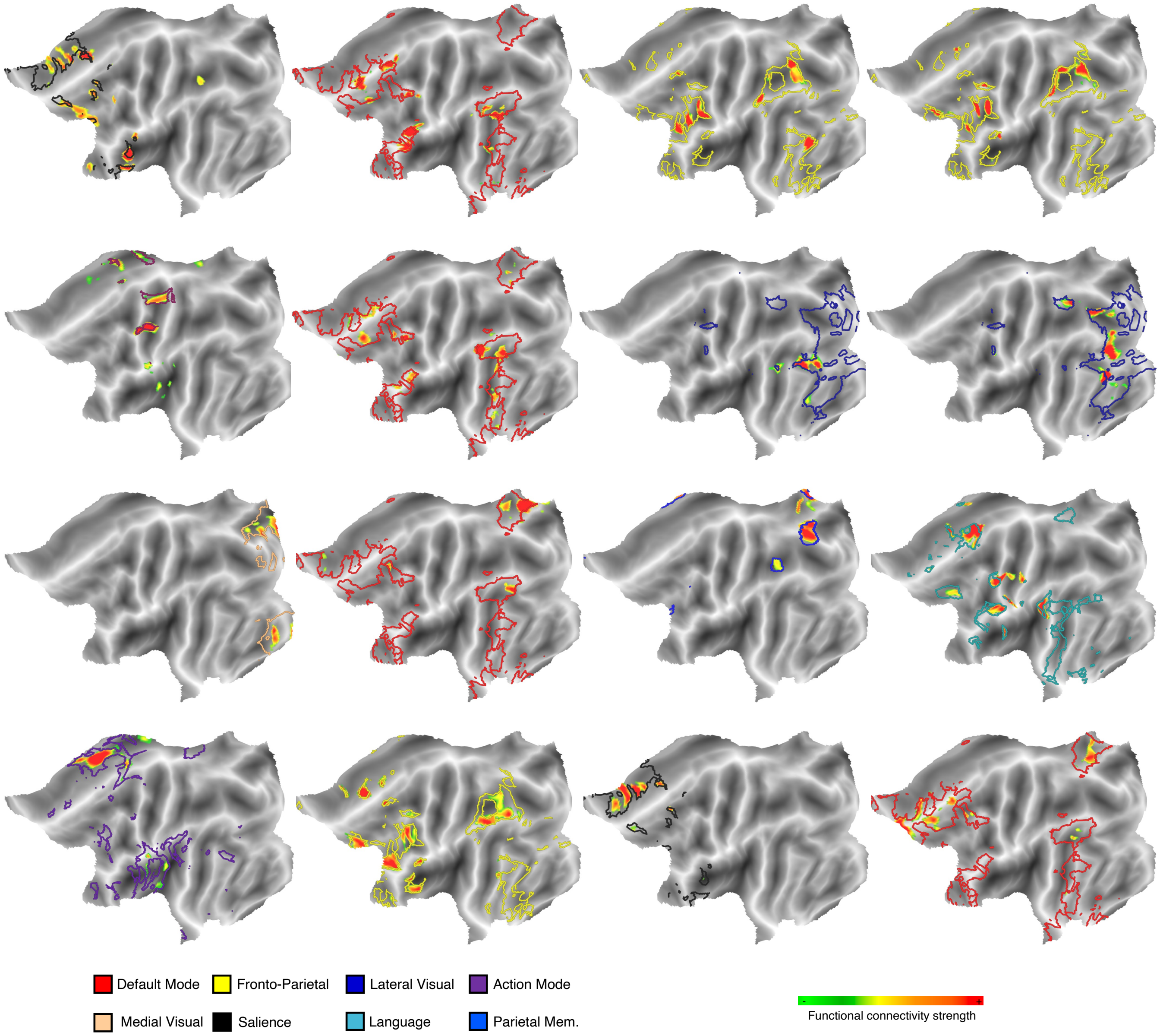


**Figure S2| Flattened surface views of overlap between chains and functional brain networks.** In an example participant (P1), sixteen cortical seeds were chosen as locations with high catenation (same seeds as in Extended Data Fig. 1). Connectivity maps generated from each seed in this participant did exhibit chained connectivity. Further, each connectivity chain fit within a large-scale functional brain network (colored outlines).

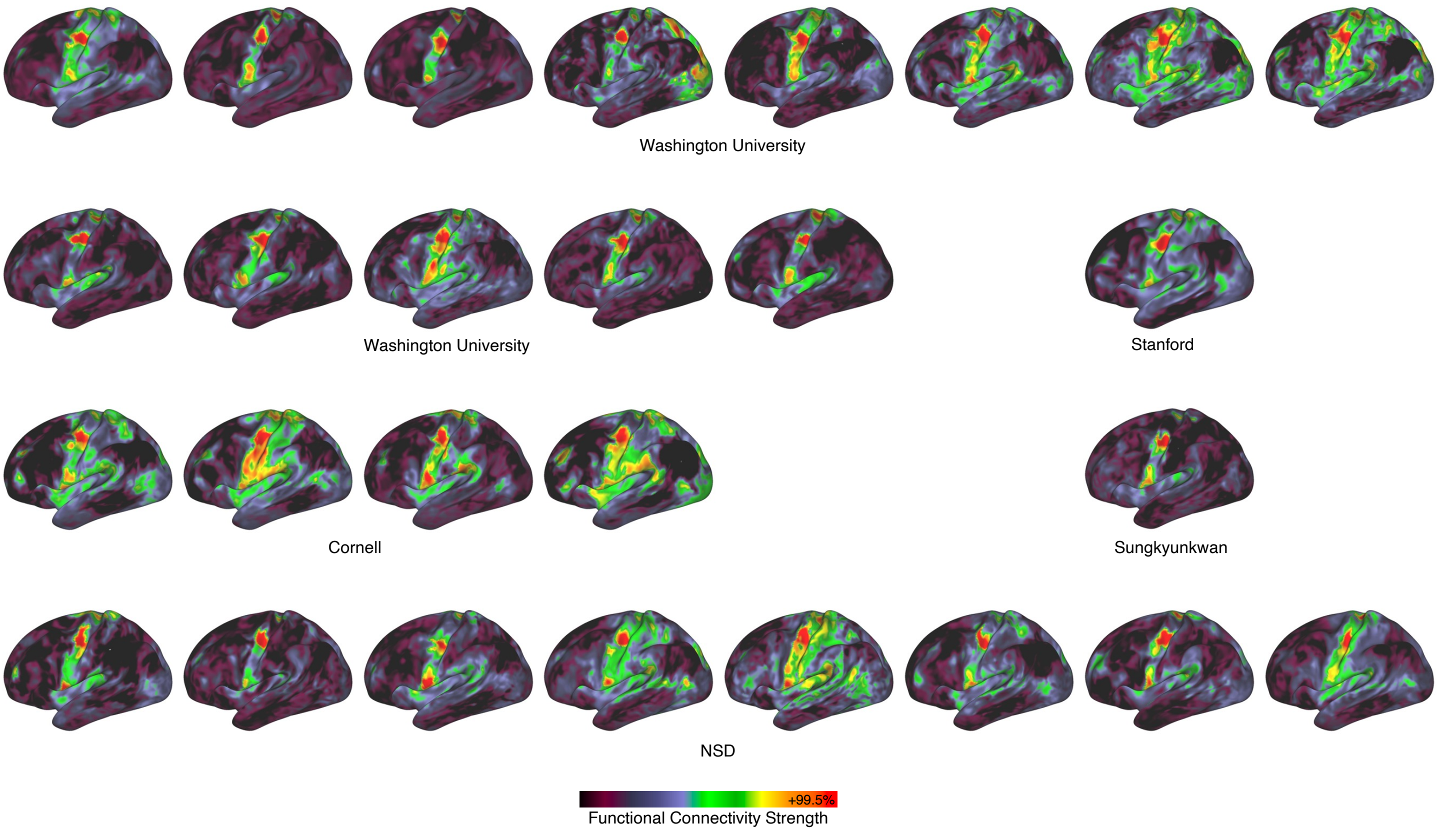


**Figure S3| Precentral gyrus somato-cognitive action network (SCAN) chains in all PFM participants.** The functional connectivity chain illustrated in Figure 1a, shown in all individual PFM participants ($n$ = 27).

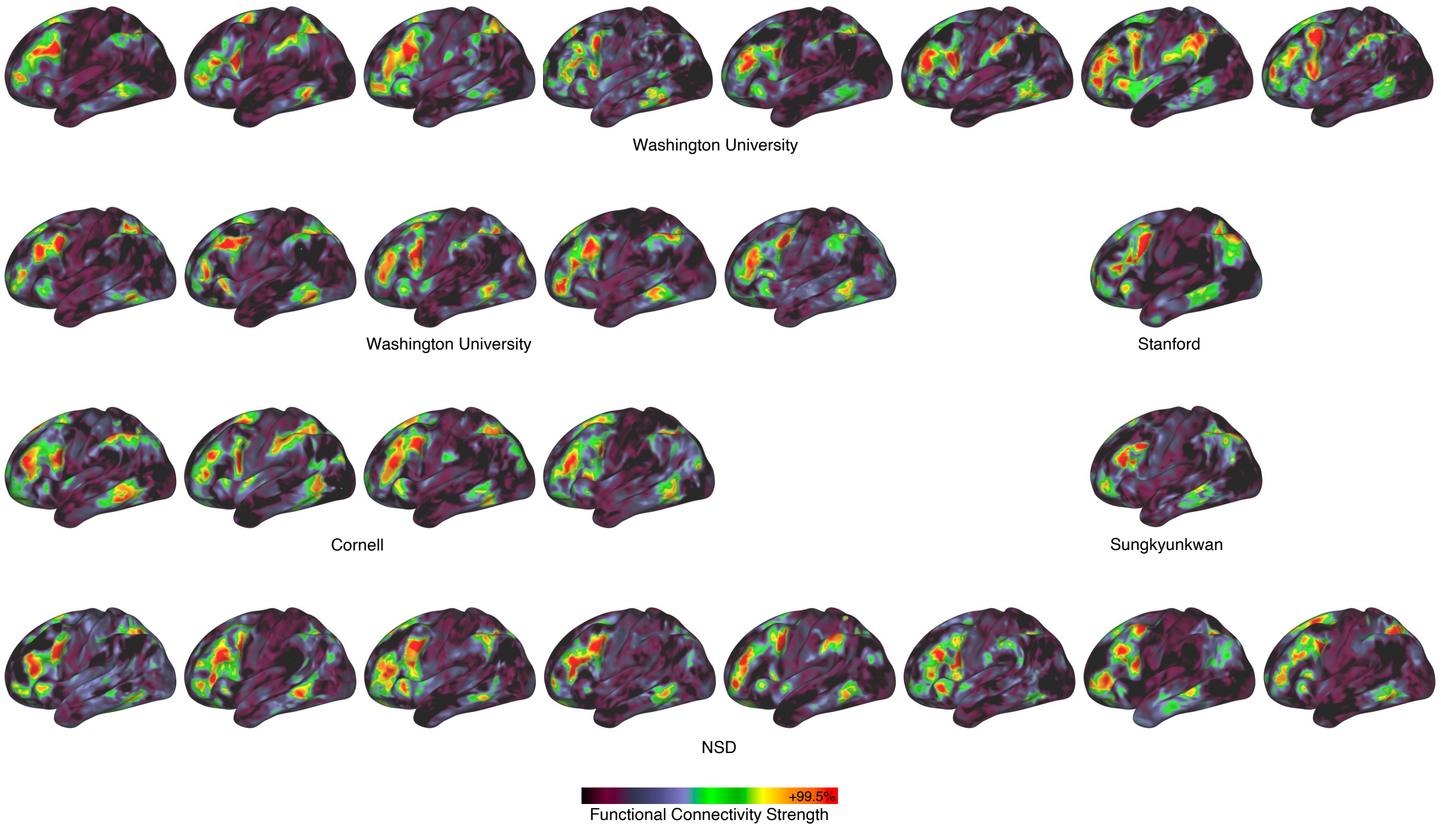


**Figure S4| Lateral prefrontal cortex chains in all PFM participants.** The functional connectivity chain illustrated in Figure 1b, shown in all individual PFM participants (*n* = 27).

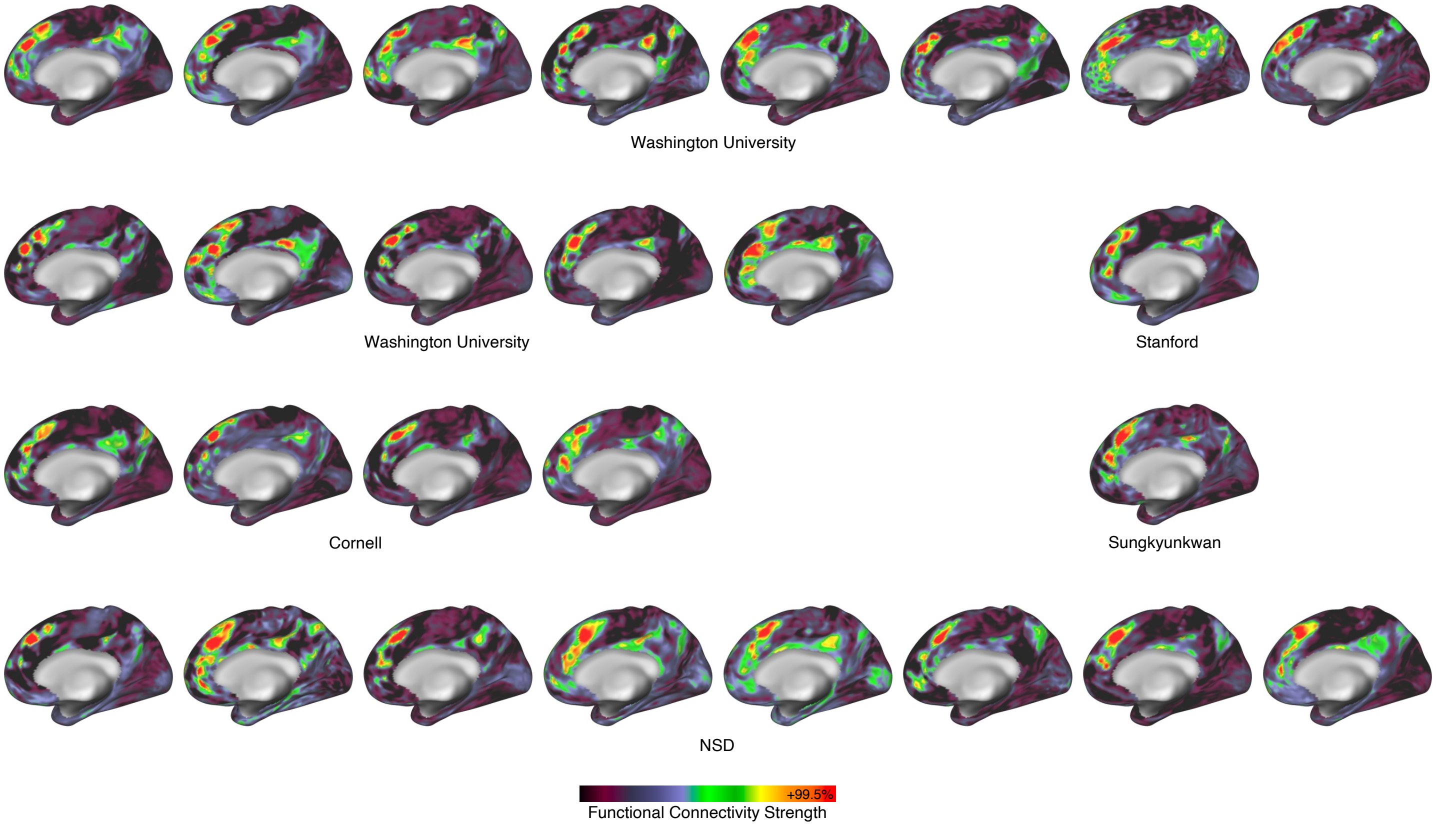


**Figure S5| Dorsomedial prefrontal cortex chains in all PFM participants.** The functional connectivity chain illustrated in Figure 1c, shown in all individual PFM participants (*n* = 27).

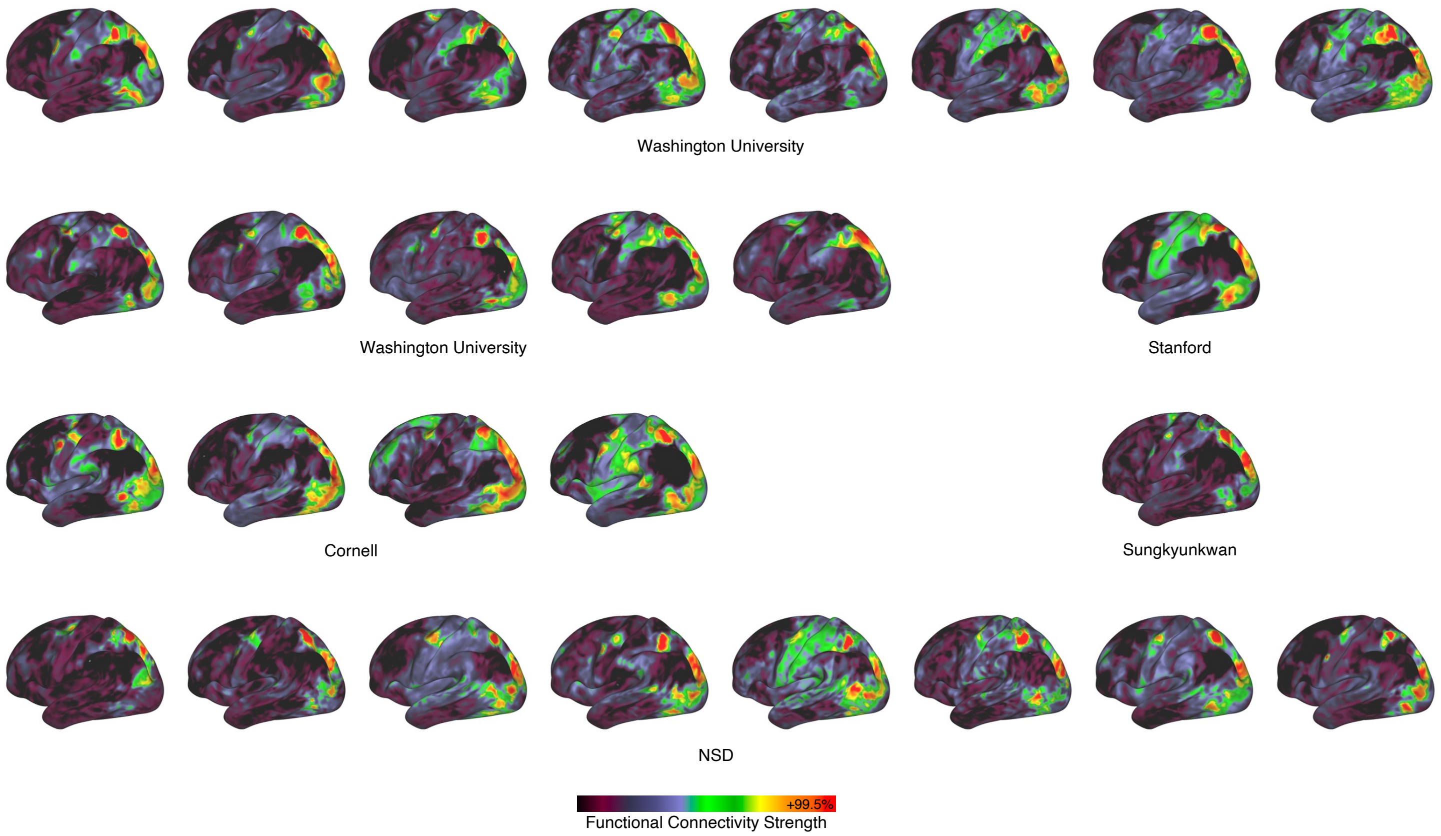


**Figure S6| Superior parietal cortex chains in all PFM participants.** The functional connectivity chain illustrated in Figure 1d, shown in all individual PFM participants (*n* = 27).

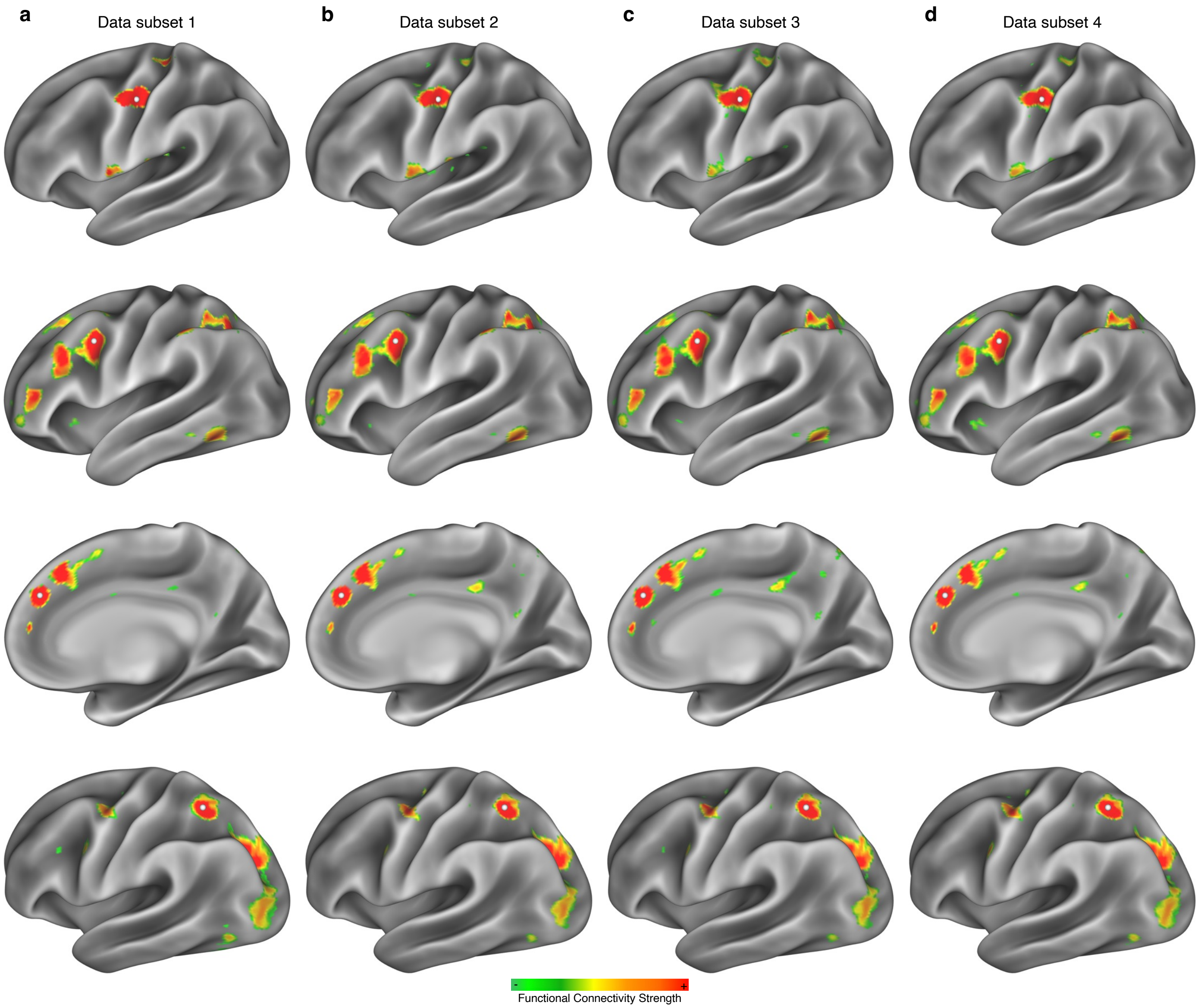


**Figure S7| Chained connectivity patterns are highly replicable within individual.** In a single example individual (P1), 477 minutes of total data were divided into four equal subsets **(a-d).** Connectivity seeded from the example seeds in Figure 1 (rows) exhibited nearly identical patterns across all data subsets.

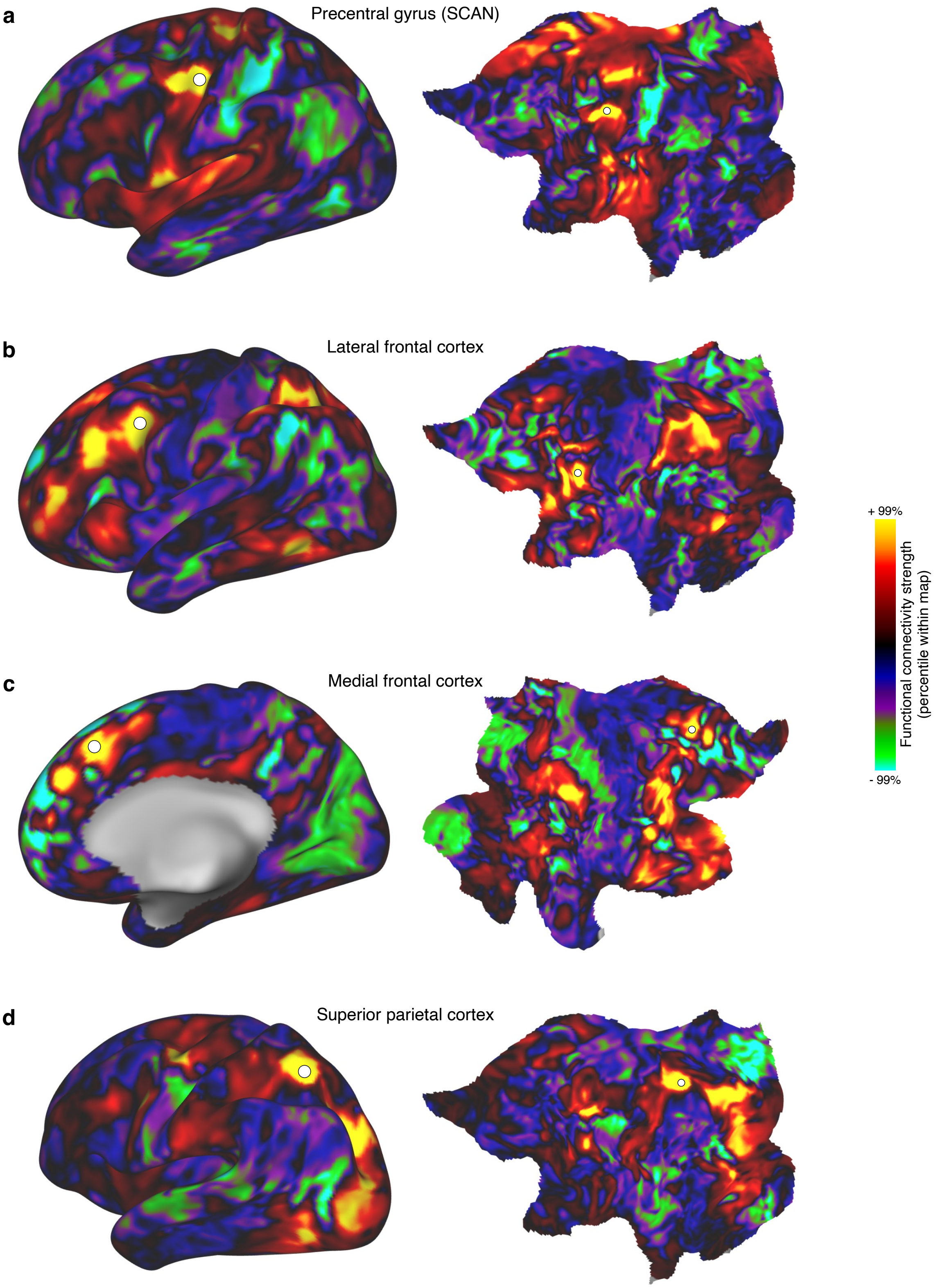


**Figure S8| Unthresholded maps of functional connectivity chains. a-d,** Unthresholded functional connectivity seed maps in a single exemplar individual (P1, as in Fig. 1), displayed on the inflated cortical surface (left) and on a cortical flat map (right). **a,** Somato-cognitive action network (SCAN) alternating with effector-specific regions in precentral gyrus (Gordon et al.,

2023) (seed MNI coordinates [x, y, z] = [-35, 15, 45]); **b,** lateral frontal cortex ([-38, 7, 36]); **c**, medial frontal cortex ([7, 26, 48]), and **d**, superior parietal cortex ([-30, -55, 50], approximate area LIPv [Glasser et al., 2016]). Seed maps revealed strongly dissociated and often anticorrelated connectivity in closely adjacent cortical locations.

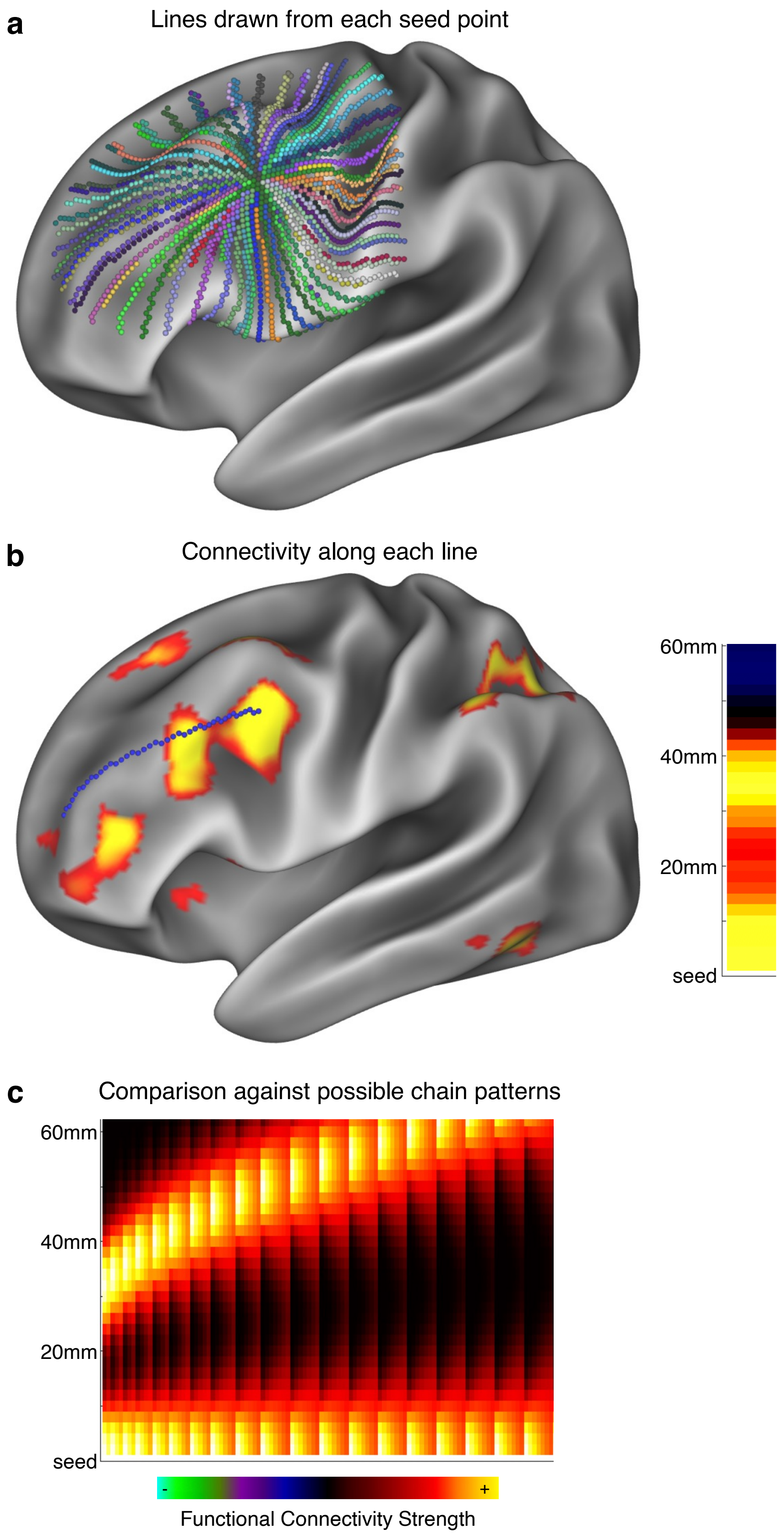
a
Lines drawn from each seed point
b
Connectivity along each line
60mm
40mm
20mm
seed
c
Comparison against possible chain patterns
60mm
40mm
20mm
seed
-
+
Functional Connectivity Strength

**Figure S9| Computing a catenation map as the similarity of functional connectivity to a set of chained patterns. a,** At each point in the brain, 60mm-long straight lines are drawn across the cortical surface with an angular spacing of 5°. See Supplementary Figure S19 for similar results using other distance values. **b,** Vertexwise functional connectivity is computed seeded from the central point (underlay). For each straight line in **a** (example line in blue), the vector of connectivity values along the line is extracted (visual representation on right). **c,** All straight-line connectivity vectors are compared against a set of 60mm-long chained connectivity vectors with two peaks, one at the seed point and one further away. The chain patterns vary in 1) the distance from the seed to the second peak (distances ranged from 30 to 60mm), and 2) the width of the peaks (sigma value of the Gaussians ranged from 6mm to 12mm). Catenation is computed as the 90$^{th}$ percentile strongest Pearson's correlation between a connectivity vector (as in **b**) lying on one of the lines in **a**, and one of the template vectors in **c**.

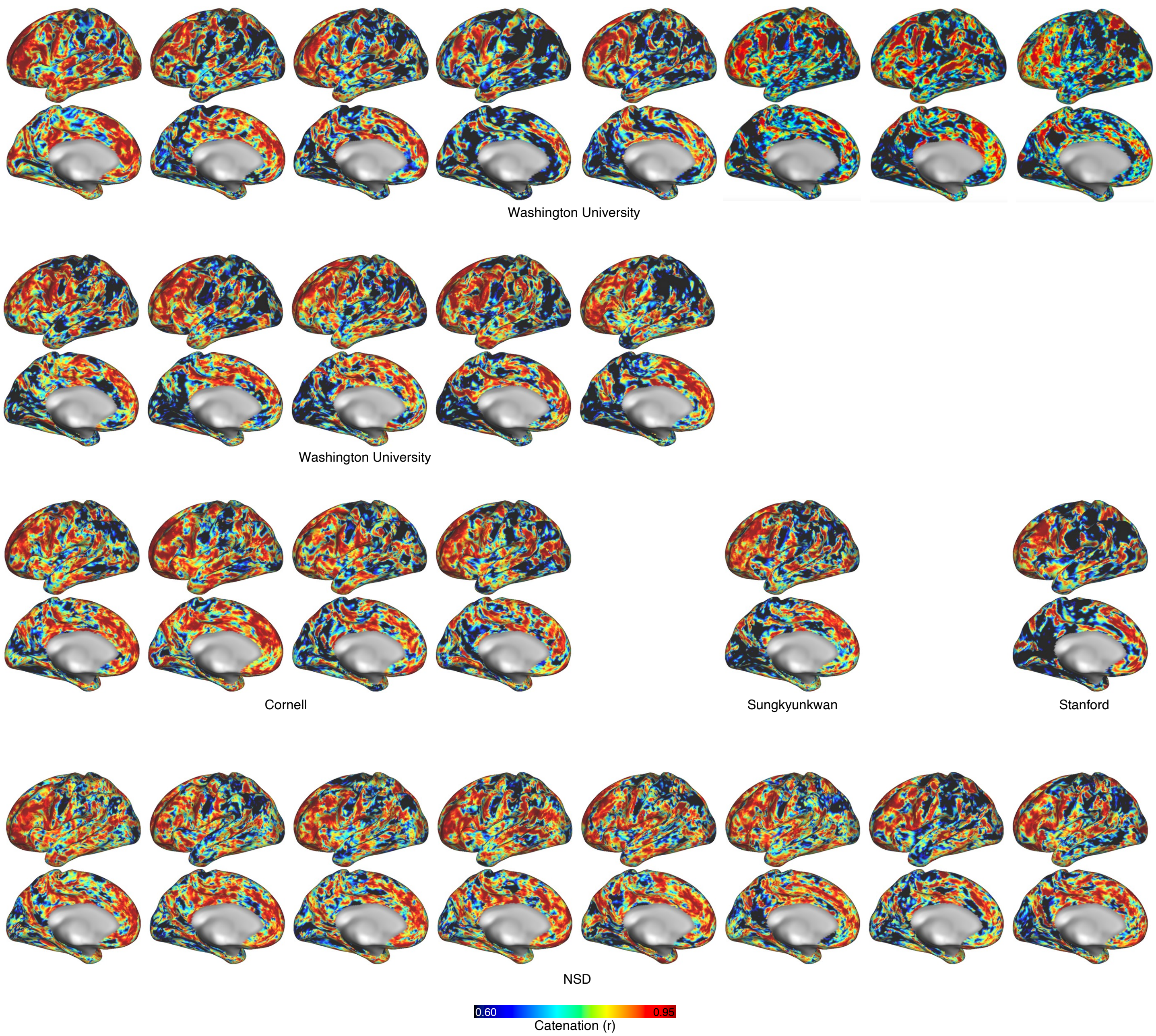


**Figure S10| Catenation maps in each PFM individual.** Catenation maps (as in Figure 2a) shown in all individual PFM participants (*n* = 27).

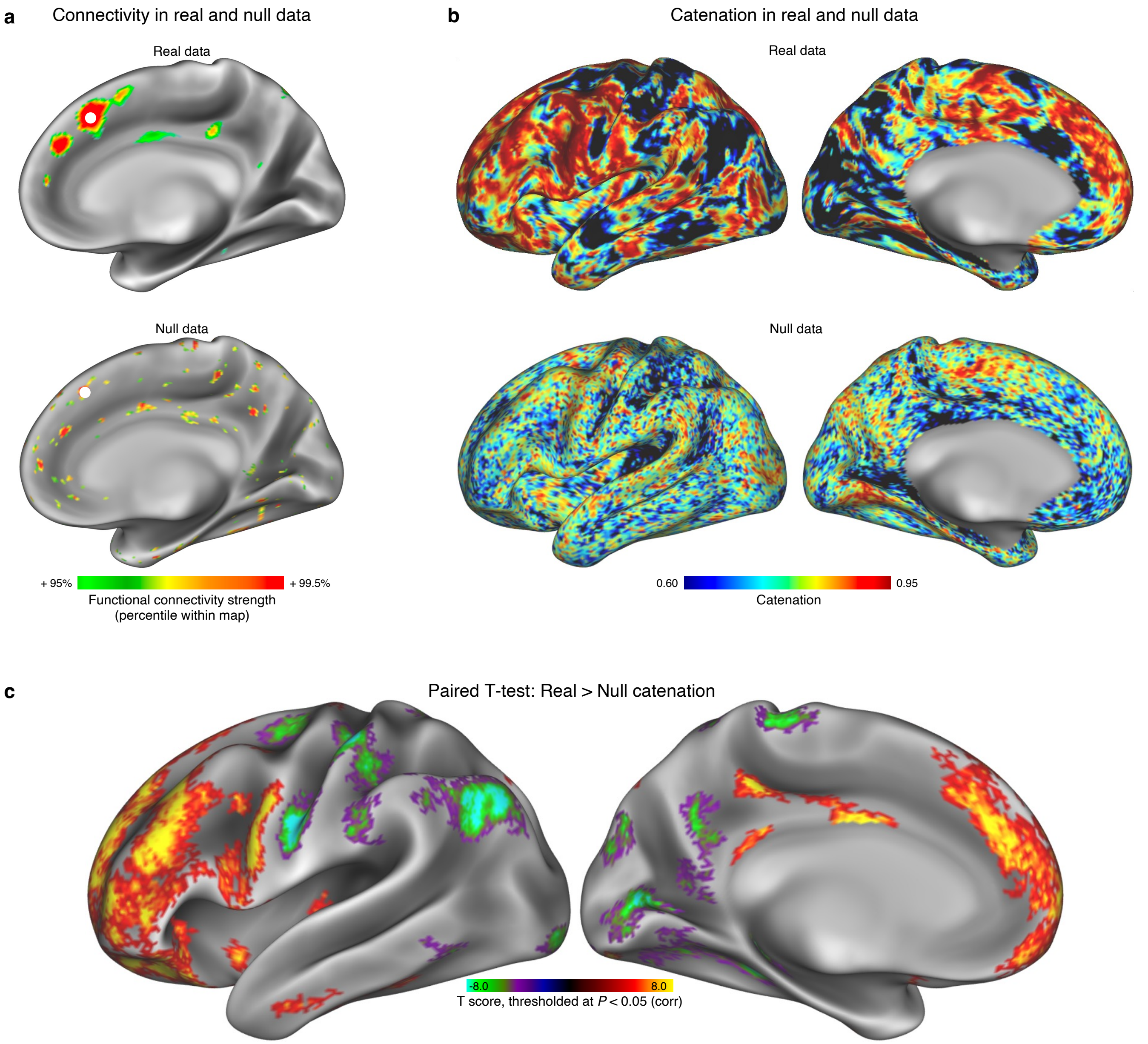


**Figure S11| Catenation relative to the null model.** A null version of each participant's data was created by phase-randomizing the original unsmoothed data to destroy any true correlation structure and re-smoothing the phase-randomized data to the same extent that the original data was smoothed. **a,** In an example participant (P1), correlation maps in real data (top) exhibited clear chains, while those generated from null data did not. **b,** Compared to catenation maps created from real data in the example participant (P1, top), catenation in null data (bottom) exhibited fewer catenated vertices (red) and fewer uncatenated vertices (black). **c,** Across participants ($n$ = 27), a vertexwise paired $t$-test demonstrated that real catenation values were consistently stronger than values computed from null data in frontal cortex and portions of insula, and were weaker (less catenated) in primary cortices, medial parietal cortex, and angular gyrus. Statistical map is thresholded at vertexwise $P$(uncorrected) < 0.001, cluster size threshold = 17, corresponding to $P$(corrected) < 0.05 by cluster-size correction. Note that nonsignificant regions here do not necessarily reflect an absence of connectivity chains, but may instead reflect a lower density of chains combined with inter-individual variability in the locations of those chains.

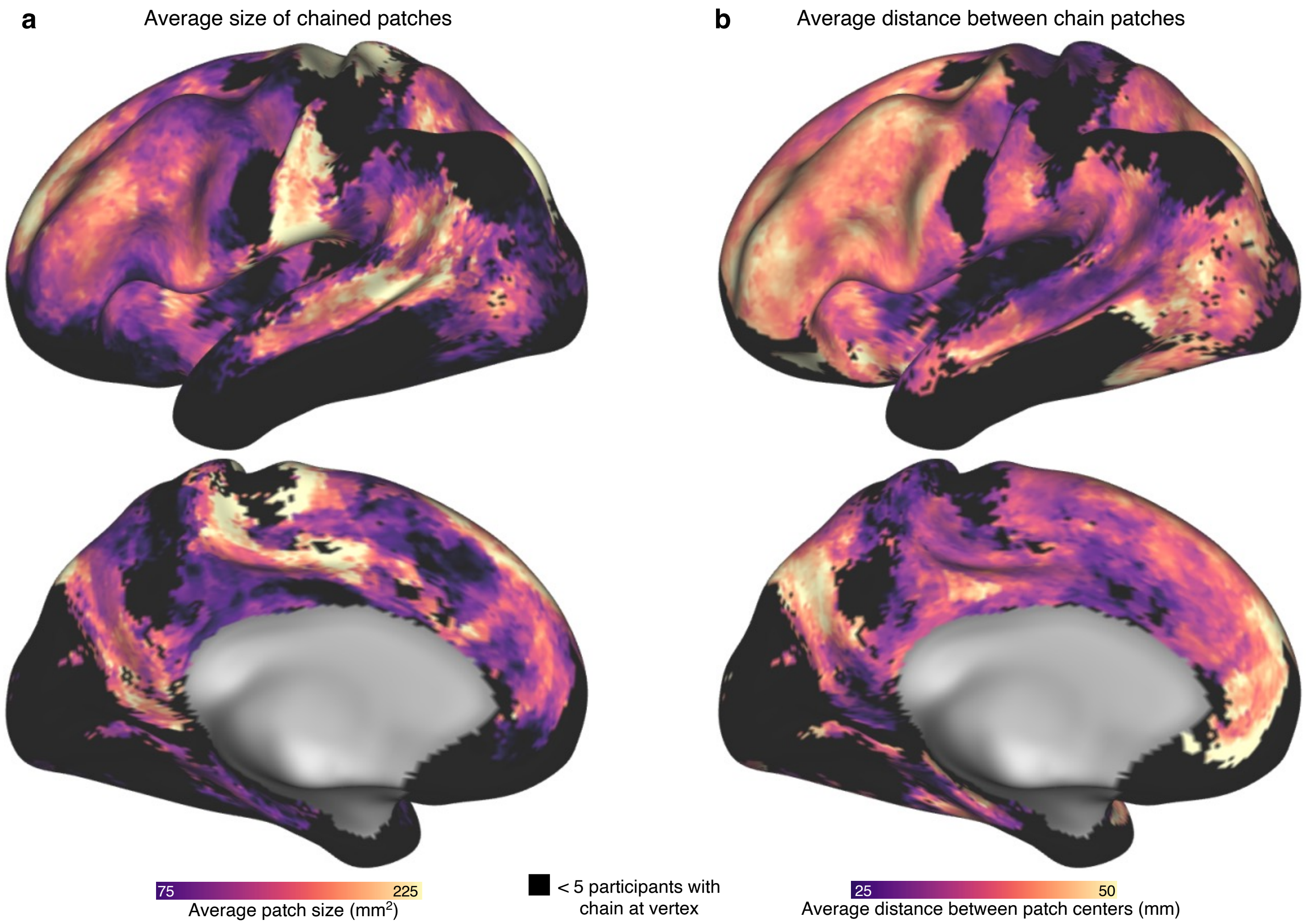


**Figure S12| Size and spacing of chained patches. a,** Average size of chained patches present at each cortical vertex, across PFM participants (*n* = 27). **b,** Average closest distance between the patches of a chain at each cortical vertex, across PFM participants. Vertices containing chain patches in fewer than five participants were zeroed out (black) to reduce instability caused by small sample sizes.

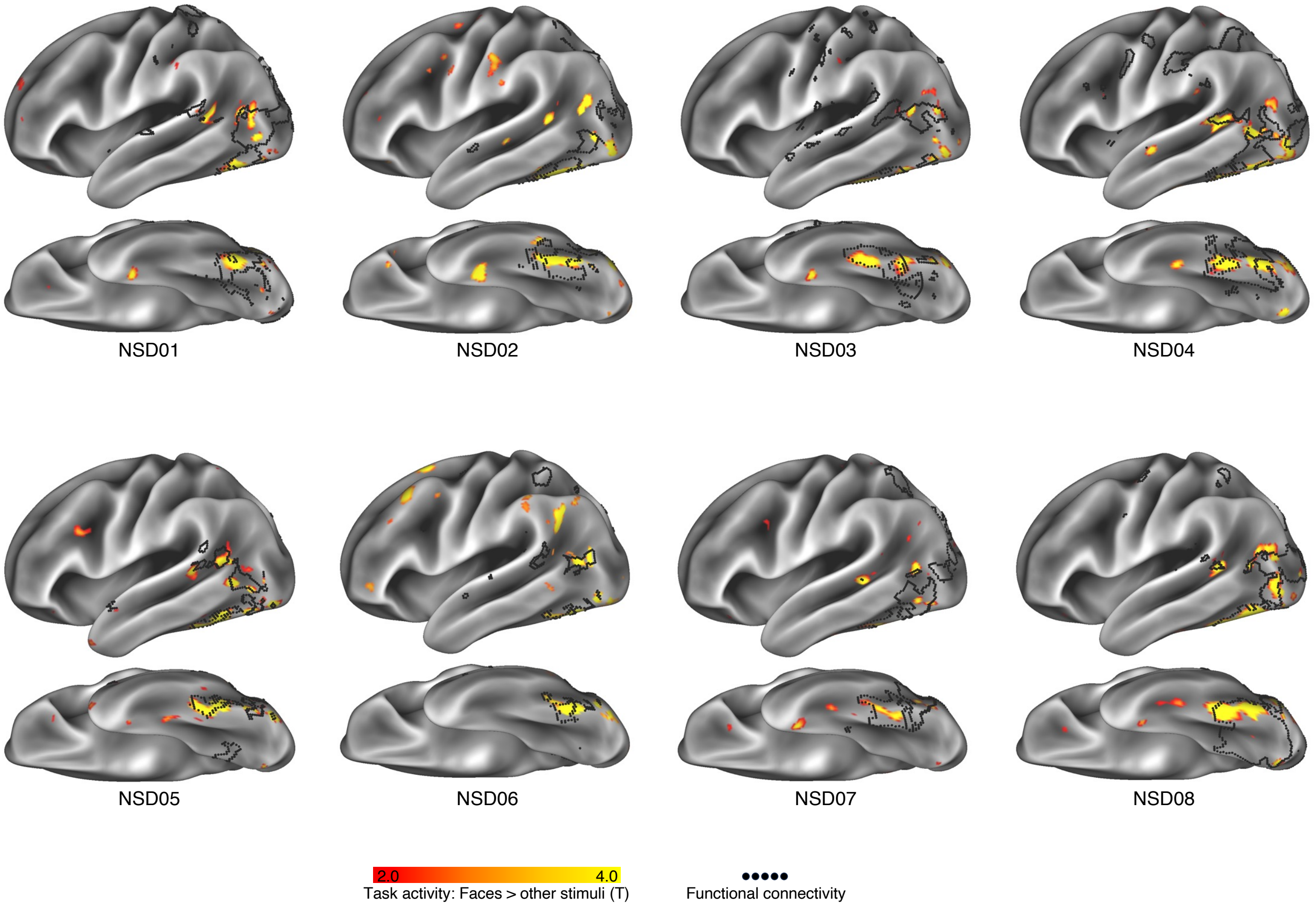


**Figure S13| Task fMRI activations mapped onto functional connectivity chains across participants.** In all participants from the NSD dataset (7 Tesla, 76-169 minutes), the overlap of face-evoked activity (from red-yellow) with functional connectivity (black outline).

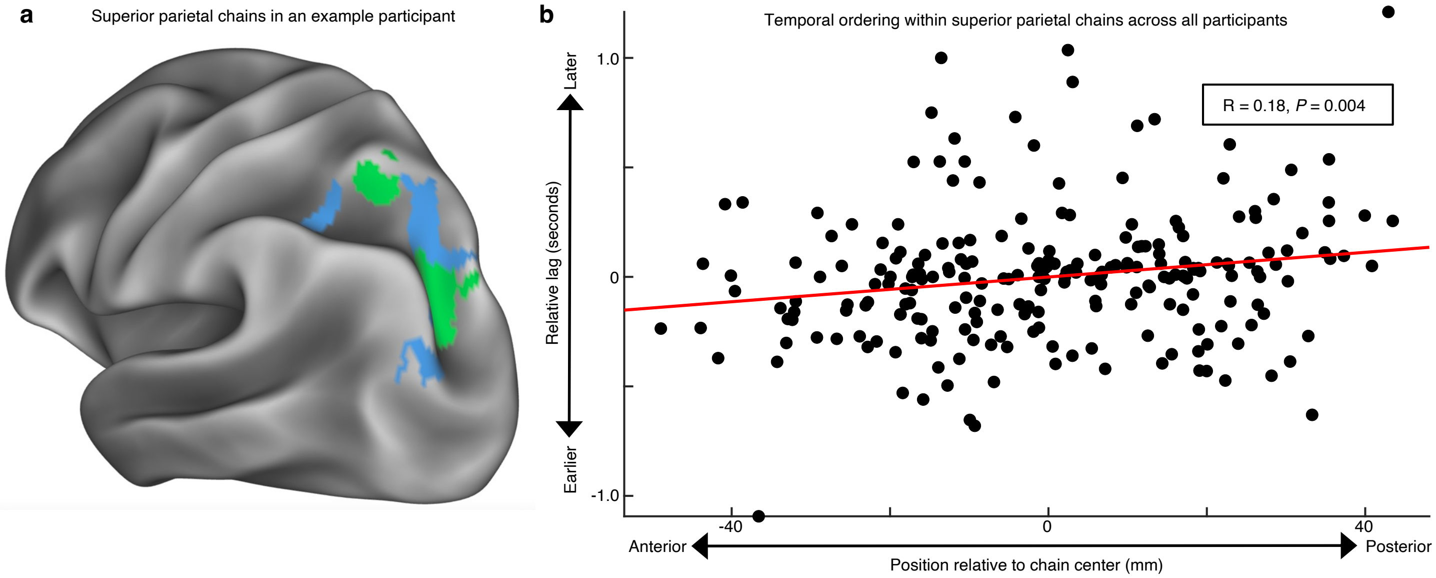


**Figure S14| Temporal ordering of signals within superior parietal connectivity chains. a,** Connectivity chains in superior parietal cortex shown for an example participant (P01). Colors indicate separate chains. **b**, Across all PFM participants, lagged correlations were computed between regions in superior parietal cortex chains. Regions of the chain that were more anterior exhibited systematically earlier signals than those more posterior (Pearson's correlation: $P$ = 0.004). Prior electrophysiology work suggests that later infra-slow (0.08-0.1Hz) activity (here, posterior chain regions) corresponds to earlier delta-band (0.5-4Hz) activity (Mitra et al., 2016).

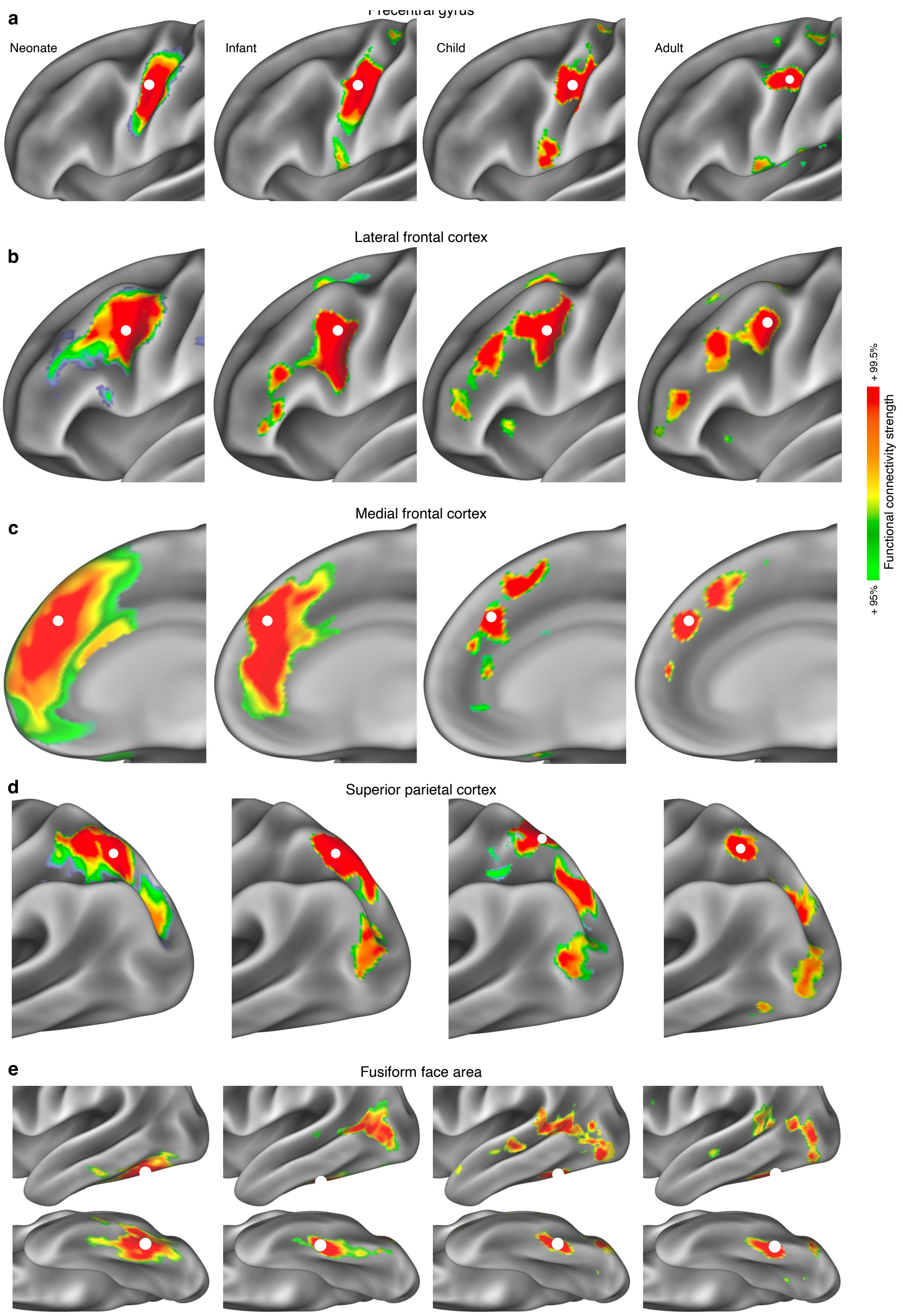
a
Precentral gyrus
Neonate
Infant
Child
Adult
b
Lateral frontal cortex
c
Medial frontal cortex
d
Superior parietal cortex
e
Fusiform face area
+ 99.5%
Functional connectivity strength
+ 95%

**Figure S15| Development of connectivity chains.** In a neonate (37 days old, left), infant (11 months old, left middle), child (9 years old, right middle), and adult (21 years old, right), functional connectivity seeded from **a,** Somato-cognitive action network (SCAN) alternating with effector-specific regions in precentral gyrus; **b,** lateral frontal cortex; **c**, medial frontal cortex, **d**, superior parietal cortex, and **e**, inferior temporal cortex (putative fusiform face area).

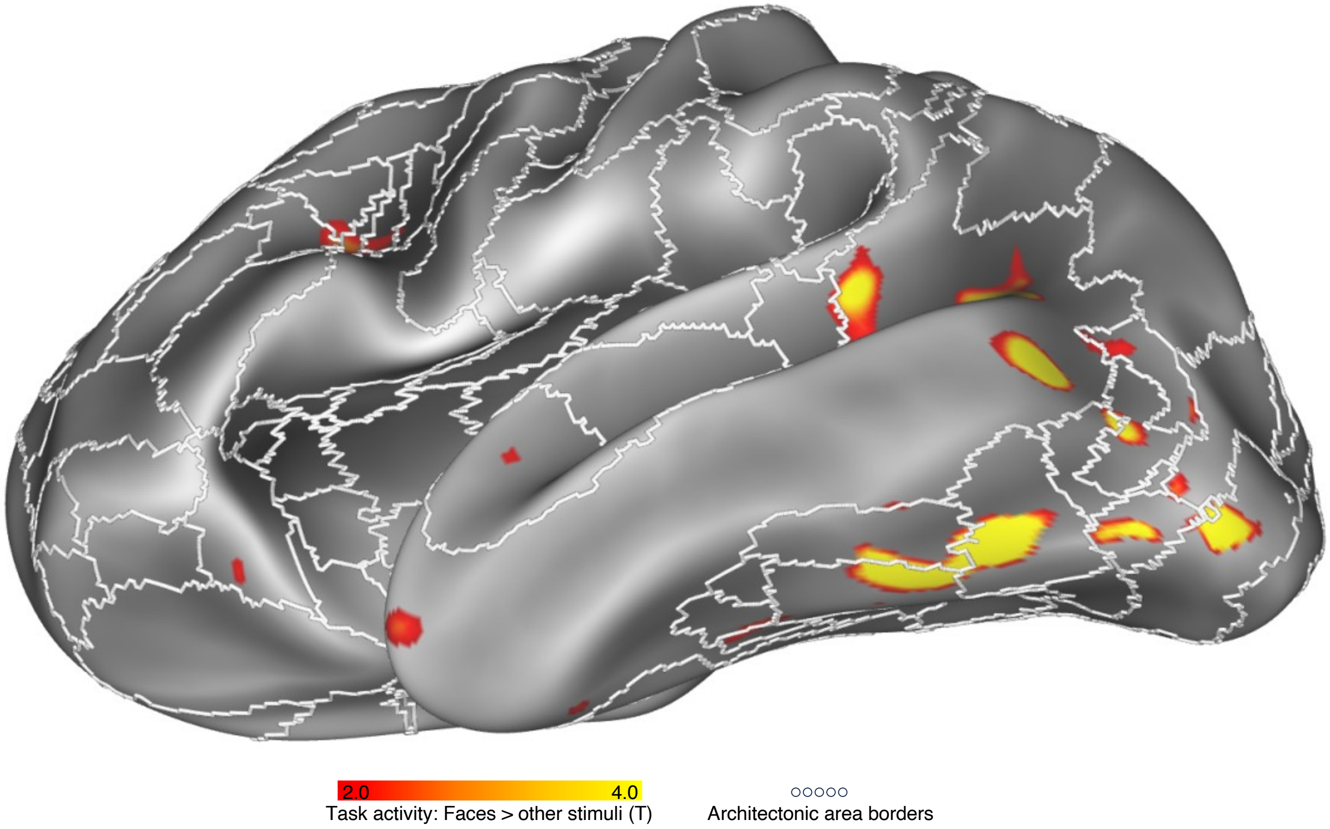


**Fig S16| Chained face patches are distributed across architectonic areas.** In an exemplar individual (NSD05) from the Natural Scenes Dataset, a task contrast localizing activity related to visual presentation of faces (face stimuli > all other stimulus types) is visualized on the ventral surface of the brain. This contrast demonstrates a chained pattern of activity in posterior lateral and ventral temporal cortex that are within different a priori architectonic areas (white borders; from [Amunts et al., 2020]). Patches fall into architectonic areas including human Occipital areas (hOc) 3v, 4lp, 4v, and 5, as well as Fusiform gyrus areas (FG) 2 and 3. More dorsal face patches on middle temporal gyrus and in superior temporal sulcus are within an architectonic gap in which divisions have not yet been delineated.

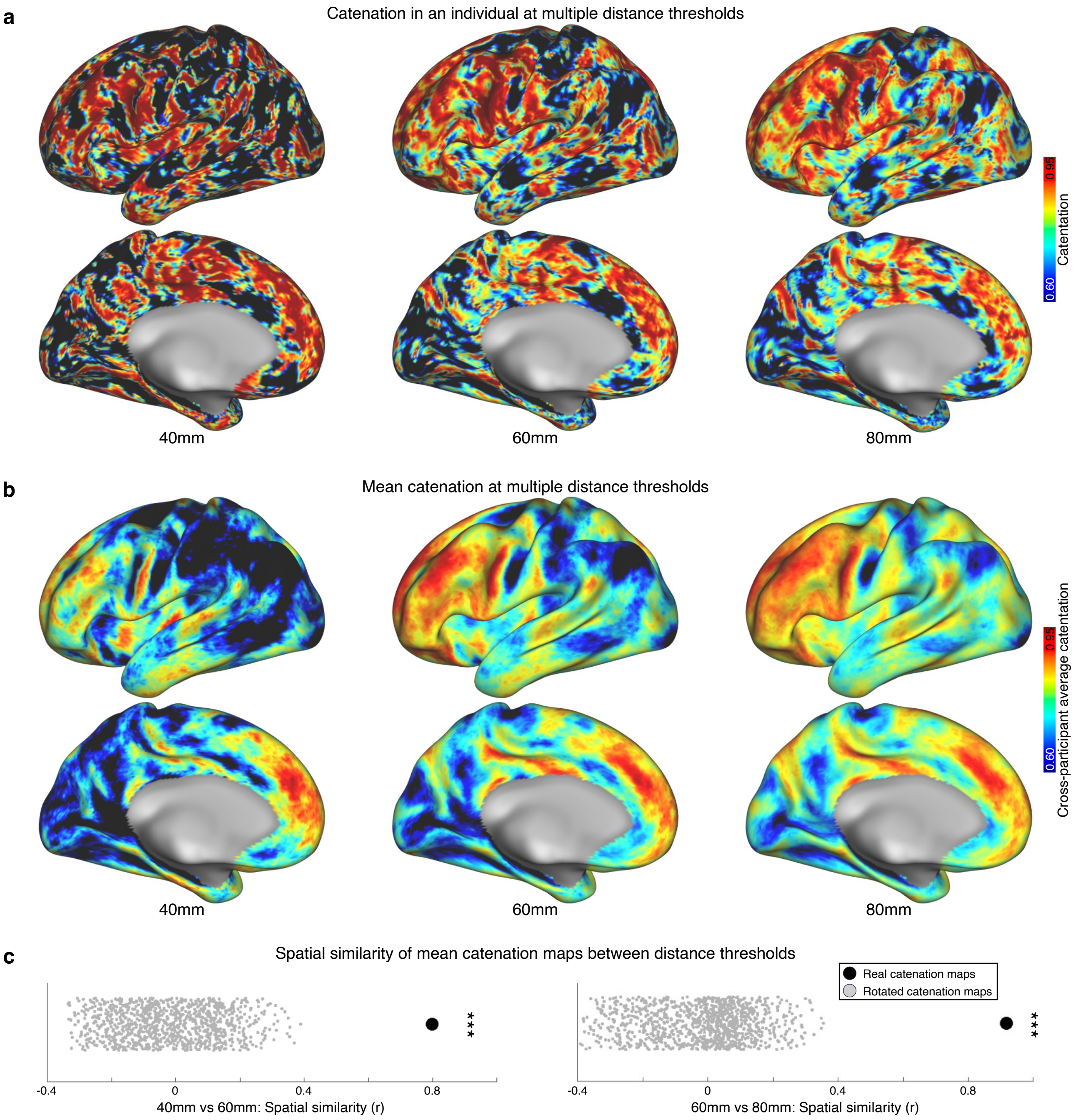


**Fig S17| Effect of distance threshold on catenation maps. a,** Catenation maps in an exemplar participant (P01) computed by examining linear vectors of functional connectivity patterns stretching 40mm, 60mm (as in Main Text), and 80mm across cortex. **b,** Catenation maps computed by examining linear vectors of functional connectivity patterns stretching 40mm, 60mm (as in Main Text), and 80mm across cortex, averaged across PFM participants. In **a** and **b**, longer distances nonspecifically increase catenation values but do not change the pattern of catenation. **c,** Similarity (spatial correlation) between average catenations maps (from **b**) computed at 40mm vs 60mm (left) and 60mm vs 80mm (right). Similarity between maps (black dots) was much greater than would be expected by chance (null rotated catenation maps: grey dots). *** indicates $P < 0.001$.

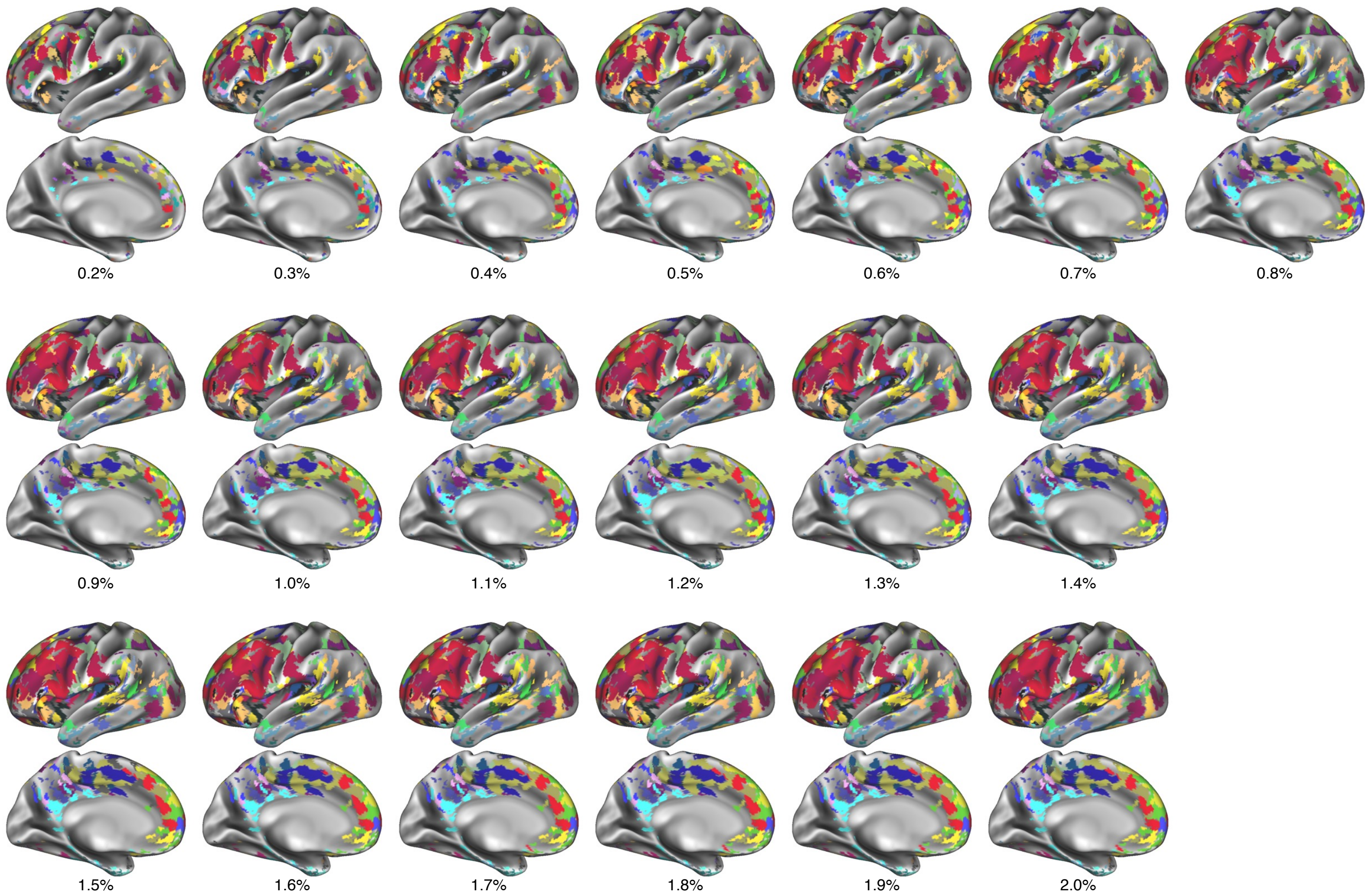


**Figure S18| Chains identified at varying density parameters.** In a single individual (P1), the chain detection algorithm was run using a range of density parameters. The density parameter determines how strong or weak the connectivity between catenated regions must be to represent a connection in the Infomap community detection algorithm. A higher density causes more connections to pass this threshold. The same chains were identified across densities, but higher densities filled the chains out to a greater extent. Different colors indicate different chains.

| | Demographics | | Scan Parameters | | |
|---|---|---|---|---|---|
| **PB ID** | *Post-Menstrual Age at Scan 1 (weeks)* | *Sex* | *SE/ ME* | *Resolution ($mm^3$)* | *TR* |
| PB001 | 42.0 | F | ME | 2.0 | 1.76 |
| PB003 | 43.3 | M | SE | 2.4 | 1.2 |
| PB004 | 44.4 | F | SE | 2.0 | 1.51 |
| PB006 | 41.7 | F | SE | 2.4 | 1.2 |
| PB008 | 42.1 | M | SE | 2.0 | 1.51 |
| PB012 | 40.9 | M | SE | 2.0 | 1.51 |
| PB013 | 43.6 | M | SE | 2.0 | 1.51 |
| PB014 | 43.0 | M | SE | 2.0 | 1.51 |

**Table S1| Demographics and scanning parameters for individual PFM neonates.**
